# A complete diploid human genome benchmark for personalized genomics

**DOI:** 10.1101/2025.09.21.677443

**Authors:** Nancy F. Hansen, Nathan Dwarshuis, Hyun Joo Ji, Arang Rhie, Hailey Loucks, Glennis A. Logsdon, Mitchell R. Vollger, Jessica M. Storer, Juhyun Kim, Eleni Adam, Nicolas Altemose, Dmitry Antipov, Mobin Asri, Sofia Barreira, Stephanie C. Bohaczuk, Andrey V. Bzikadze, Sara A. Carioscia, Andrew Carroll, Kuan-Hao Chao, Yanan Chu, Arun Das, Peter Ebert, Adam English, Mark Fleharty, Laura E. Fleming, Giulio Formenti, Andrea Guarracino, Gabrielle A. Hartley, Katharine Jenike, Jenna Kalleberg, Yu Kang, Robert King, Josipa Lipovac, Mira Mastoras, Matthew W. Mitchell, Shloka Negi, Nathan D. Olson, Keisuke K. Oshima, Luis F. Paulin, Brandon D. Pickett, David Porubsky, Jane Ranchalis, Desh Ranjan, Mikko Rautiainen, Harold Riethman, Robert D. Schnabel, Fritz J. Sedlazeck, Kishwar Shafin, Mile Sikic, Steven J. Solar, Alexander P. Sweeten, Winston Timp, Justin Wagner, DongAhn Yoo, Ying Zhou, Erik Garrison, Evan E. Eichler, Michael C. Schatz, Andrew B. Stergachis, Rachel J. O’Neill, Karen H. Miga, Steven L. Salzberg, Sergey Koren, Justin M. Zook, Adam M. Phillippy

**Affiliations:** Genome Informatics Section, Center for Genomics and Data Science Research, National Human Genome Research Institute, National Institutes of Health, Bethesda, MD, USA; Material Measurement Laboratory, National Institute of Standards and Technology, Gaithersburg, MD, USA; Department of Computer Science, Johns Hopkins University, Baltimore, MD, USA; Center for Computational Biology, Johns Hopkins University, Baltimore, MD, USA; UC Santa Cruz Genomics Institute, University of California, Santa Cruz, CA, USA; Department of Genetics, Epigenetics Institute, Perelman School of Medicine, University of Pennsylvania, Philadelphia, PA, USA; Division of Medical Genetics, Department of Medicine, University of Washington School of Medicine, Seattle, WA, USA; Institute for Systems Genomics, University of Connecticut, Storrs CT, USA; Department of Computer Science, Old Dominion University, Norfolk, VA, USA; Department of Genetics, School of Medicine, Stanford University, Palo Alto, CA, USA; Chan Zuckerberg Biohub, San Francisco, San Francisco, CA USA; Computational Genomics Unit, Center for Genomics and Data Science Research, National Human Genome Research Institute, National Institutes of Health, Bethesda, MD, USA; Graduate Program in Bioinformatics and Systems Biology, University of California, San Diego, La Jolla, CA, USA; Department of Biology, Johns Hopkins University, Baltimore, MD, USA; Google LLC, Mountain View, CA, USA; Beijing Institute of Genomics, Chinese Academy of Sciences, and China National Center for Bioinformation, Beijing, 100101, China; Core Unit Bioinformatics, Medical Faculty and University Hospital Düsseldorf, Heinrich Heine University, Düsseldorf, Germany; Center for Digital Medicine, Heinrich Heine University Düsseldorf, Germany; Human Genome Sequencing Center, Baylor College of Medicine, Houston, TX, USA; Broad Institute, Broad Clinical Labs, Burlington, MA, USA; The Vertebrate Genome Laboratory, The Rockefeller University, New York, USA; Department of Genetics, Genomics and Informatics, University of Tennessee Health Science Center, Memphis, TN, USA; Department of Plant Sciences, University of Cambridge, Cambridge, UK, CB2 3EA; Division of Animal Sciences, University of Missouri, Columbia, MO, USA; Oxford Nanopore Technologies, Oxford, UK; Laboratory for Bioinformatics and Computational Biology, Faculty of Electrical Engineering and Computing, University of Zagreb, Zagreb, Croatia; Coriell Institute for Medical Research, Camden, NJ, USA; Department of Genome Sciences, University of Washington School of Medicine, Seattle, WA, USA; Genome Biology Unit, European Molecular Biology Laboratory (EMBL), Heidelberg, 69117, Germany; Institute for Molecular Medicine Finland, Helsinki Institute of Life Science, University of Helsinki, Helsinki, Finland; School of Medical Diagnostic & Translational Sciences, Old Dominion University, Norfolk VA, USA; Department of Molecular and Human Genetics, Baylor College of Medicine, TX, USA; Department of Computer Science, Rice University, Houston, TX, USA; Genome Institute of Singapore, A*STAR; Department of Biomedical Engineering, Johns Hopkins University, Baltimore, MD, USA; Department of Data Science, Dana-Farber Cancer Institute, Boston, MA, USA; Howard Hughes Medical Institute, University of Washington, Seattle, WA, USA; Brotman Baty Institute for Precision Medicine, Seattle, WA, USA; Department of Molecular and Cell Biology, University of Connecticut, Storrs CT, USA; Department of Genome Sciences, UConn Health, Farmington, CT, USA; Department of Biostatistics, Johns Hopkins University, Baltimore, MD, USA

## Abstract

Human genome resequencing typically involves mapping reads to a reference genome to call variants; however, this approach suffers from both technical and reference biases, leaving many duplicated and structurally polymorphic regions of the genome unmapped. Consequently, existing variant benchmarks, generated by the same methods, fail to assess these complex regions. To address this limitation, we present a telomere-to-telomere genome benchmark that achieves near-perfect accuracy (i.e. no detectable errors) across 99.4% of the complete, diploid HG002 genome. This benchmark adds 701.4 Mb of autosomal sequence and both sex chromosomes (216.8 Mb), totaling 15.3% of the genome that was absent from prior benchmarks. We also provide a diploid annotation of genes, transposable elements, segmental duplications, and satellite repeats, including 39,144 protein-coding genes across both haplotypes. To facilitate application of the benchmark, we developed tools for measuring the accuracy of sequencing reads, phased variant call sets, and genome assemblies against a diploid reference. Genome-wide analyses show that state-of-the-art de novo assembly methods resolve 2–7% more sequence and outperform variant calling accuracy by an order of magnitude, yielding just one error per 100 kb across 99.9% of the benchmark regions. Adoption of genome-based benchmarking is expected to accelerate the development of cost-effective methods for complete genome sequencing, expanding the reach of genomic medicine to the entire genome and enabling a new era of personalized genomics.

## Introduction

Variant benchmarks developed by the Genome in a Bottle (GIAB) Consortium have played a pivotal role in advancing the accuracy of human genome sequencing by providing a truth against which experimental protocols and analysis pipelines can be evaluated^1,2^. Motivated by competitive assessments, the top variant calling pipelines can now achieve F1 scores approaching 0.999 on the latest benchmarks^3^, appearing to leave little room for improvement. However, these variant benchmarks are limited in scope, as they currently exclude from consideration structural variation and the most polymorphic regions of the genome, which are often implicated in human genetic disease. For example, GIAB’s v4.2.1 small variant benchmark for the HG002 genome excluded both sex chromosomes and 12% of the autosomes^4^. The variant-centric framework has limitations, primarily due to the mapping-based approaches used to construct the benchmarks.

Although suitable for many applications, mapping-based resequencing fails to genotype many variants within duplicated or structurally polymorphic regions of the genome. For example, short-read sequencing alone is not able to phase variants by haplotype or resolve highly similar duplications, including multi-copy gene families. Linked or long reads partially address this technical bias^5,6^, and pedigrees can be used to further improve variant filtering in complex regions^7^, but some bias persists whenever variants are called against a reference genome. This arises from errors, gaps, and natural variation in the reference that interfere with read mapping and variant representation, resulting in a lack of variant calls within sequences that are missing or otherwise unalignable to the reference. A complete reference genome, such as T2T-CHM13, addresses the problem of errors and gaps^8,9^ but does not address the issue of structural variation between the sample and reference genomes. A pangenome reference provides a more diverse panel of haplotypes for mapping, thereby reducing reference bias^10^. However, both the development and validation of pangenome resources have relied on benchmarking against incomplete references such as GRCh38^11^. This creates a circular dependency which makes it difficult to establish their accuracy in repetitive and polymorphic regions of the genome—precisely where pangenomes should provide the greatest benefit.

To address these challenges, sophisticated benchmarking tools have been developed to compare different representations of unphased small variants^12^, phased, complex variants^13^, and variants within tandem repeats^14^. However, regions with complex structural and copy-number variation remain difficult to assess. Therefore, even when these regions can be accurately resolved with *de novo* assembly^15^, they remain excluded from benchmarking due to the lack of appropriate standards^16^. For example, GIAB’s latest assembly-based benchmarks for HG002 exclude many Class II MHC genes^17^ as well as 122 medically relevant genes that are considered difficult to genotype due to the presence of complex or copy-number variants, repetitive sequence, and segmental duplications^18^.

To overcome these limitations, we introduce the concept of a telomere-to-telomere “genome benchmark”, which uses the complete diploid genome sequence of the sample as a ground truth, rather than a set of variants against a reference genome (**Figure 1**). Genome benchmarking is more flexible and avoids the inherent biases of variant benchmarking, while simultaneously refocusing the objective away from variant calling and towards “genome inference”, where the goal is to directly infer the haplotypes of the genome being sequenced^19,20^. A complete diploid sequence is the most fundamental representation of a genome, against which single-nucleotide, phasing, and structural accuracy can be reliably assessed either locally or across the entire genome. In fact, a common way to unify complex variant representations has been to compare the underlying sequences themselves^21–23^. Genome benchmarking simply extends this idea to the whole genome and breaks the circular dependency of prior variant benchmarks.

**Figure 1.**
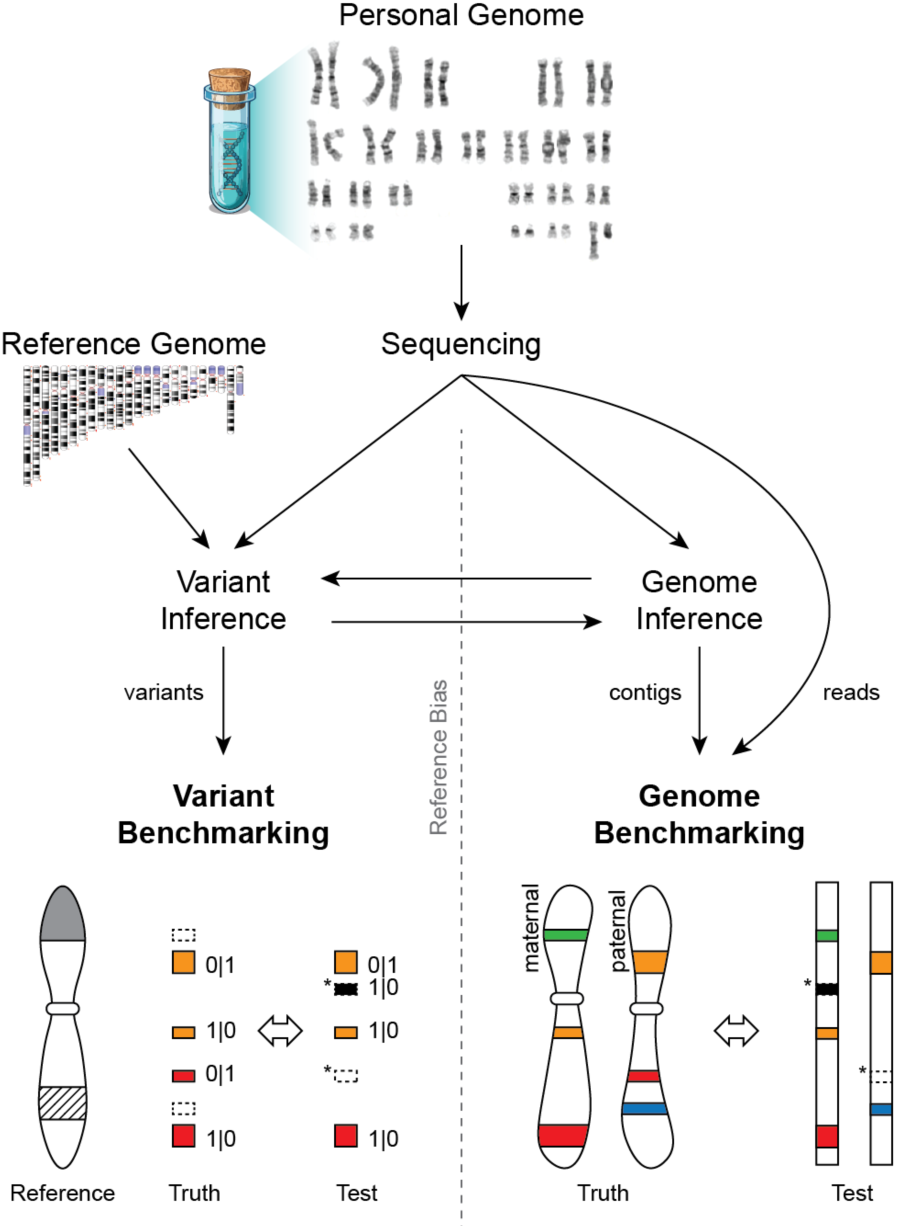
Variant benchmarking versus genome benchmarking. *Variant Inference* (i.e. variant calling) refers to the process of identifying a set of variants (differences) between the sequencing data and a reference genome. *Genome Inference* refers to the process of constructing a genome sequence either *de novo* or with the assistance of a reference genome or pangenome. It is possible to move between the two frameworks, either by aligning contigs to a reference to call variants (right to left) or by applying a set of variants to a reference to construct contigs (left to right). *Variant Benchmarking* compares a set of test variants against a set of known variants to identify false positive and false negative calls (asterisks). However, this approach is typically affected by *Reference Bias*, especially when the reference is missing sequence (gray) or contains complex regions that are difficult to represent in the benchmark (hashed). *Genome Benchmarking* compares a set of sequences (either assembled contigs or reads) directly to the complete, diploid genome they originated from, removing the issue of reference bias and enabling a more comprehensive evaluation.

The field’s reliance on variant benchmarking has been partly driven by the inability to sequence and assemble complete human genomes. However, the Telomere-to-Telomere Consortium recently overcame this barrier with the T2T-CHM13 assembly^8^, completing the final 8% of the genome and revealing amplified gene arrays^8,24^, segmental duplications^25^, and genomic repeats^26,27^ associated with critical genome functions. Recent developments streamlining the assembly of complete, diploid, telomere-to-telomere (T2T) chromosomes^28,29^ have resulted in additional T2T reference genomes for human^30–33^, apes^34,35^, and other vertebrates^36–38^. These technological advances enable the creation of comprehensive genome benchmarks for the thorough evaluation of new sequencing and analysis methods.

Here we present results from the “Q100 Project”, an effort to reconstruct the complete, diploid genome of the widely used GIAB sample HG002 with the goal of essentially perfect accuracy (i.e. a Phred score of Q100, meaning an error rate of less than one per 10 Gb). Version 1.1 of the T2T-HG002 genome benchmark achieves telomere-to-telomere continuity for all 46 chromosomes, with only the interior of the rDNA arrays left unfinished; has been thoroughly validated using at least four different sequencing technologies; is free of detectable errors over 99.4% of the genome; and far exceeds both the completeness and accuracy of all prior benchmarks. In conjunction, we provide the Genome Quality Checker (GQC) software for benchmarking any type of sequence data against this new resource, including *de novo* genome assemblies, raw sequencing reads, phased variant calls, and pangenome-inferred haplotypes. These new resources remove the performance ceiling of past variant benchmarks, allowing all current and future technologies to be compared on equal footing and encouraging the development of improved methods for the sequencing of complete, personalized genomes.

## Results

### A complete, nearly perfect assembly of the HG002 genome

The genome “HG002” is from the son of an Ashkenazi family trio (father-mother-son) originally sampled by the Personal Genome Project (PGP)^39^ and now widely used as a sequencing technology benchmark (NIST Reference Material 8391, Coriell lymphoblastoid cell line (LCL) GM24385, PGP participant huAA53E0). The National Institute of Standards and Technology (NIST) has created a reference material derived from an expansion of the GM24385 cell line, showing a mostly diploid 46,XY karyotype (78% of sampled cells) with lower levels of tetraploidy (18%) and inversions (3%) (46,XY[57]/92,XXYY[13]/46,XY,?inv(3)(q26.3q29)[3], **Supplementary Information**, **Figure S1**). The T2T Consortium initially finished HG002’s X chromosome for comparison to CHM13^8^ and later added HG002’s Y chromosome to the T2T-CHM13v2.0 reference, which lacked its own ChrY^24^. HG002 was also used as a benchmarking genome for the Human Pangenome Reference Consortium (HPRC)^40^ and for the development of automated T2T assembly methods^28^.

To complete the entire HG002 genome, we combined roughly 170× coverage of PacBio circular consensus (HiFi) and 209× of Oxford Nanopore Technologies (ONT) ultra-long reads from the HPRC^41^ and Genome in a Bottle Consortium (GIAB)^42^ (**Table S1**). An initial Verkko^28^ assembly was generated from these reads along with additional Illumina reads from HG002’s parents (HG003 and HG004) for phasing, and Strand-seq^43^ and Hi-C reads^44,45^ for scaffolding across the rDNA arrays (**Supplementary Information**). Validation of the rDNA scaffolding and estimation of the rDNA array sizes were guided by fluorescence *in situ* hybridization^46^, and rDNA consensus sequences were assembled by Ribotin^47^. All other gaps were closed by patching with alternate assemblies or by manual resolution of the assembly graph in a similar manner to CHM13^8^. T2T-HG002v0.7 was released to GitHub in November of 2022.

We iteratively polished, patched, and validated the T2T-HG002v0.7 assembly in three rounds using multiple independent short- and long-read datasets (**Supplementary Information**). Due to the large quantity of available sequencing data for HG002, we adopted a custom polishing approach to better understand the nature of assembly errors, rather than using only automated methods^48,49^. During the first two rounds of polishing, crowdsourced curation of randomly selected examples of several large correction sets was tracked using issues in a public GitHub repository (**Data Availability**), along with curation assignments and analysis of the reliability of different correction categories. Each correction set targeted particular technologies and/or modes of error (e.g. missing heterozygous sites callable with short reads), and those correction sets for which more than half of the curated corrections were validated as correct were then applied to the assembly (**Supplementary Information, Table S2, Table S3**). After the first polishing round, version v0.9 of the assembly was released to GitHub in July of 2023; after a second round of polishing, v1.0.1 was submitted to NCBI GenBank in December of 2023 (GCA_018852605.2, GCA_018852615.2); and, after a third round of polishing, v1.1 in July of 2024 (GCA_018852605.3, GCA_018852615.3). In total, the three rounds of polishing resulted in 38,037 small corrections made to the assembly, including 5,420 single-base substitutions, 26,779 small deletions (50 or fewer base pairs), and 5,628 small insertions (**Figure S2**), along with 210 larger consensus patches of regions totaling 14,182,804 base pairs. Analysis of Strand-seq^50,51^ reads confirmed the overall structural correctness of the v1.1 assembly, showing no evidence of inversion errors (**Supplementary Information**, **Figure S3**). Unless otherwise stated, the rest of the manuscript concerns version T2T-HG002v1.1, or T2T-HG002 for brevity (**Figure 2A**).

**Figure 2:**
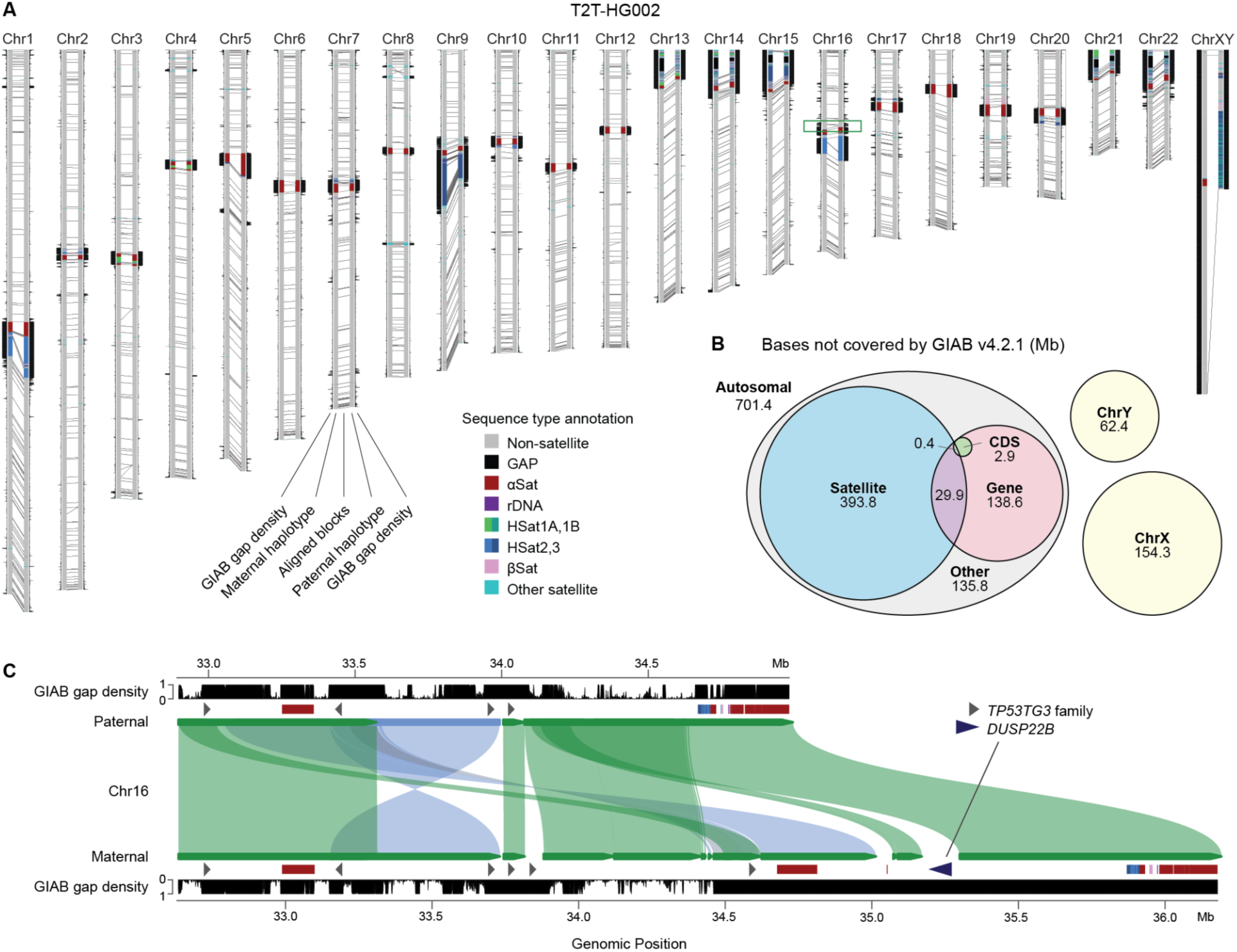
The complete, diploid genome of HG002. **(A)** Genome ideogram of the T2T-HG002 assembly. Each homologous chromosome pair is displayed with its short arms at the top and the maternal copy to the left of the corresponding paternal copy. Satellite repeat annotations highlight the centromeres and other heterochromatic regions. Gray lines connecting the maternal and paternal chromosomes delineate the boundaries of aligned (homologous) blocks. On the outer sides of the haplotypes, density plots show the fraction of bases within 50 kb windows that are *not* included in the GIAB v4.2.1 variant benchmark “high confidence regions” with larger values farther from the haplotype. A maximum value of one indicates a window that is entirely missing from the variant benchmark, such as within the centromeres or on the sex chromosomes, and a value of 0 indicates all bases in a window are covered by the benchmark. 99.7% of all windows genome-wide (119,674 / 120,013) include at least some bases not covered by the variant benchmark. **(B)** Breakdown by sequence class of the bases missing from the GIAB v4.2.1 variant benchmark that are covered by the genome benchmark. For the autosomal bases: Satellite = all DNA satellite annotations. Gene = all annotated genic elements, including introns. CDS = coding sequence. The entire sequence of both sex chromosomes is missing. **(C)** Enlargement of the p-arm pericentromere of HG002 Chr16 showing the complex alignment of paternal and maternal haplotypes (forward: green, inverted: blue), satellite and protein-coding gene annotations, and gaps in the variant benchmark (same “GIAB gap density” plotted in panel A). The *DUSP22B* gene copy only appears on the maternal haplotype and is absent from GRCh38, but present in T2T-CHM13v2.0 (not shown), explaining why the variant benchmark does not cover this region.

We estimated the Phred-scaled consensus quality (QV) and phasing accuracy of the assembly with Merqury^52^ using a database of PacBio HiFi and Element 31-mers from HG002 and Illumina parental 31-mers from HG003 and HG004. After polishing, QV increased from Q63.1 for the v0.7 assembly to Q68.9 for the v1.1 assembly (Q76.2 using 21-mers, **Table S4, Table S5**). This equates to reducing the estimated error rate from roughly one per two million bases to one per eight million bases, which is substantially lower than all prior HG002 assemblies tested^10,53,54^, (**Table S4)**. Interestingly, we noted that certain correction sets resulted in little-to-no QV improvement, due to the limitations of *k*-mer-based methods at such low error rates. This included corrections that restored heterozygous variants, or were made within long homopolymers, tandem repeats, and repetitive sequences, where the *k*-mer-based methods are unable to detect errors. Additionally, while evaluating the assembly for phasing error, we noted large blocks of haplotype switches flagged in the immunoglobulin gene loci. We confirmed this was caused by V(D)J recombination in the LCLs^55^, making the parental-specific markers unreliable in these regions. V(D)J recombination was confirmed in the NIST reference material by comparison to short reads from a PBMC-derived induced pluripotent stem cell (iPSC) of the same individual (Coriell GM27730). Consequently, all immunoglobulin loci were excluded from parental haplotype switch analyses.

Altogether, 99.35% of the diploid genome length is free of any detectable errors in T2T-HG002v1.1. The remaining 3,172 low-confidence regions cover 38.8 Mb (0.65%) of the 6.0 Gb assembly (**Table S6, Table S7**). The majority (86.3%) of these low-confidence bases are composed of rDNA sequence arrays which were flagged in their entirety as low-confidence. These arrays include nine gaps (Ns) spanning 31.1Mb, a resolved rDNA array on the paternal copy of Chr13 spanning 275.9 kb, and 2.4Mb of rDNA sequence flanking the gaps. The remaining 5.3 Mb of low-confidence sequence (0.09% of the assembly) was flagged by either base-level or structural evaluation, erring on the side of caution to ensure the integrity of the benchmark. Included in these flagged regions are 293 regions (183.1 kb) that contain 31-mers not found in either PacBio HiFi Revio SPRQ and Element UltraQ reads, as well as 794 possible haplotype switch errors (31.6 kb) indicated by 31-mers from parental Illumina reads. Mapped PacBio and ONT reads flagged 237 regions (4.9 Mb) with excessive secondary alleles present in the alignments of both technologies, indicative of a possible structural or consensus issue^56^. Lastly, 1,893 regions (486.6 kb) were flagged by DeepTrio^57^ using Element reads; 410 (82.9 kb) by DeepPolisher and DeepVariant^48^; 19 (3.8 kb) by Sniffles^58^; 20 (4.5 kb) via contributed GitHub issue submissions; and 3 (44.7 kb) by T2T-Polish^59^. These low-confidence regions are excluded from the v1.1 benchmark, and their future validation and/or correction will be tracked in the HG002 GitHub issues repository.

Lastly, the assembly was compared to GIAB’s most recent (v4.2.1) HG002 variant benchmark. In addition to the sex chromosomes, T2T-HG002 includes an additional 701.4 Mb (11.7%) of high-confidence autosomal sequence that is absent from the v4.2.1 variant benchmark, mostly covering satellite repeats but also hundreds of Mb of non-satellite, typically segmentally duplicated, sequence (**Figure 2B**). In regions reliably covered by both benchmarks (covering approximately 2.66 Gb of the variant benchmark regions), polishing from v0.7 to v1.1 reduced the number of discrepancies between the genome and variant benchmarks from 5,972 to 219 variants. Nearly all of the remaining discrepancies were found to be GIAB v4.2.1 errors in complex variants or difficult-to-map regions. Manual curation of 20 randomly selected variants outside of segmental duplications indicated only two errors and one possible mosaic variant in the T2T-HG002v1.1 assembly, all within homopolymers and dinucleotide tandem repeats. Within segmental duplications (covering 81 Mb), 1,015 differences remain, but inspection of 20 randomly selected variants suggests these should have been excluded from the GIAB v4.2.1 benchmark due to reference bias; all were within regions that could not be reliably mapped between HG002 and GRCh38 due to repeats, structural variation, or gene conversion. An example of such a region is shown in **Figure 2C**, where the maternal Chr16 pericentromere contains an insertion of *DUSP22* that does not exist on the paternal haplotype or GRCh38, both of which only have *DUSP22* copies near the p-arm telomere of Chr6. As such, this region of the maternal haplotype is entirely absent from prior HG002 variant benchmarks, despite *DUSP22* being a known human-specific duplicated gene family associated with several cancers^60,61^.

### Personalized genome annotation

We generated gene annotations for each haplotype of the T2T-HG002 assembly using Liftoff^62^ to map genes from T2T-CHM13v2.0 (see **Supplementary Information**). In addition, we used LiftOn^63^ and miniprot^64^ to annotate additional gene copies based on protein-to-genome alignment of translated MANE (v1.4) transcripts (a high-quality annotation in which a single representative transcript is selected for each protein-coding gene). Mitochondrial and immunoglobulin genes were excluded from the annotation. The resulting annotation includes 59,380 genes on the maternal haplotype (MAT), including ChrX, and 57,728 genes on the paternal haplotype (PAT), including ChrY (**Table 1**). Since the CHM13 assembly is haploid and contains both sex chromosomes, combining HG002 MAT with both ChrX and ChrY for comparison totals 60,074 annotated genes compared to 57,319 for CHM13, an increase of 2,755 (not including rRNAs). Only 39 distinct genes were absent from both T2T-HG002 haplotypes relative to T2T-CHM13 (**Table S8**), with all three missing protein-coding genes (*HLA-DRB5*, *CT45A8*, *OPN1MW*) known to naturally vary in copy number. For example, HG002 contains only one copy of the X-linked *OPN1MW/OPN1MW2/OPN1MW3* opsin gene family^65^, which Liftoff annotated as *OPN1MW3*.

**Table 1.** Number of genes annotated in T2T-HG002v1.1 and T2T-CHM13v2.0. Gene biotypes were identified using the attribute “gene_biotype” taken from the T2T-CHM13 reference annotation, except for protein-coding genes, which were identified based on whether or not a gene included a CDS annotation. The T2T-CHM13 counts include all autosomes as well as both ChrX and ChrY.

| Gene type | HG002<br>MAT 1–22 | HG002<br>PAT 1–22 | HG002<br>MAT ChrX | HG002<br>PAT ChrY | CHM13<br>All Chrs |
| --- | --- | --- | --- | --- | --- |
| **Protein coding** | 19,117 | 19,084 | 838 | 105 | 19,635 |
| **lncRNA** | 17,889 | 17,818 | 377 | 165 | 17,850 |
| **Other** | 20,049 | 20,132 | 1,110 | 424 | 19,834 |
| **Total** | 57,055 | 57,034 | 2,325 | 694 | 57,319 |

These gene counts exclude all rRNA genes due to the remaining rDNA gaps in the HG002 assembly. The v1.1 assembly contains 49 individual rDNA copies in scaffolds and 100 Ribotin morphs (i.e. consensus sequences for distinct rDNA units) are made available separately as unplaced contigs. Prior work estimates over 600 total rDNA copies in the HG002 genome^46^, with approximately 430 on MAT and 197 on PAT, which is substantially higher than the ∼400 copies in CHM13. In contrast to the 45S rDNA arrays on the acrocentric chromosomes, both 5S arrays on Chr1 are complete in HG002 comprising 91 MAT copies and 63 PAT copies, compared to 99 in CHM13.

The complete, annotated diploid assembly of HG002 allowed us to compare the total gene content between maternal and paternal haplotypes of the same individual. The primary questions we addressed were: (1) how many protein-coding genes are “broken” in each haplotype; and (2) how many genes have different copy numbers between the two haplotypes? We focused our analysis of broken genes on those contained in MANE. This restricted our analysis to 18,976 genes on MAT and 18,231 on PAT. A protein-coding gene was defined as “broken” if the mapped MANE transcript met at least one of the following criteria: (1) an invalid start and/or stop codon; (2) a premature in-frame stop codon; or (3) transcripts whose protein translation was less than 80% identical to the corresponding MANE protein. Using these criteria, we identified 129 broken protein-coding genes on MAT and 121 on PAT (**Table S9)**. Broken genes were enriched within segmental duplications, with 28% (69 / 250) intersecting an annotated segmental duplication, compared to just 8% (2996 / 36,957) of the unbroken genes. One such example, located within a segmentally duplicated region of Chr15, is the human-specific fusion gene *CHRFAM7A*^66^, which is a dominant negative regulator of α7 neuronal nicotinic acetylcholine receptor (*CHRNA7*) function^67^. HG002 PAT contains a common 2 bp frame-shifting deletion in exon 6 that is associated with increased risk of various neuropsychiatric disorders, but the recently duplicated, polymorphic nature of this locus complicates analysis^68^. Contrary to previous literature that suggests the 2 bp deletion allele is typically inverted relative to the wild-type^69^, both HG002 MAT (wild-type) and PAT (Δ2 bp) copies are in the same orientation as the GRCh38 and CHM13 references.

Next, we computed gene copy numbers across both T2T-HG002 haplotypes (**Supplementary Information**, **Table S10**) and identified haplotype-specific genes (i.e. those unique to only one haplotype). After filtering to reduce false positives from closely related genes and excluding sex chromosomes, we identified 14 MAT-only and 12 PAT-only autosomal genes (**Supplementary Information**, **Table S11**). Our filtered set of haplotype-specific genes includes several well-characterized examples of copy-number variable genes, such as *DUSP22* (**Figure 2C**), as well as *CFHR1*, *CFHR3*, *GSTT1*, and *GSTM1*. HG002 PAT includes a co-deletion of *CFHR1* and *CFHR3*, which is a fairly common haplotype (20% of chromosomes in a UK-based cohort) shown to protect against age-related macular degeneration^70^. The glutathione S-transferase genes *GSTM1* and *GSTT1* are another pair of copy-number variable genes in HG002 that play a role in the neutralization of toxic compounds, with homozygous deletions (null alleles) being linked to an increased risk of cancer in certain populations^71,72^. HG002 is heterozygous for both, containing a single copy of *GSTM1* only on PAT and a single copy of *GSTT1* only on MAT. Excluding genes that were entirely absent from one haplotype, we identified 56 genes with a higher copy number on the MAT haplotype and 38 with a higher copy number on PAT. These differences reflect true variation and highlight the value of a haplotype-resolved assembly in structurally variable regions of the genome.

To enable easy access and exploration, both maternal and paternal haplotypes of T2T-HG002 have been organized along with additional data tracks in a UCSC Genome Browser hub^73^. In addition to the diploid gene annotation, we generated sequence-based annotation tracks for GC percent, CpG islands, repetitive elements, segmental duplications, subtelomere repeats, and centromeric satellites. Validation tracks provide the locations of all known assembly issues, depth of coverage for mapped HiFi and ONT reads, and secondary variants reported by NucFreq^56^. Whole-genome alignments are also included, showing the position of all heterozygous sites and enabling quick locus switching and lift over between HG002 haplotypes, T2T-CHM13, and GRCh38. We also provide multiple tracks derived from HG002 long-read functional genomics experiments, including: 5-methylcytosine (5mC) predictions from both HiFi and ONT sequencing, chromatin accessibility and regulatory element predictions from Fiber-seq^74^, and bulk RNA and single-cell transcriptome data from PacBio Iso-Seq Kinnex and MAS-Seq. Uniquely, because this data is based on long-read sequencing and drawn from matched HG002 cells, it can be confidently mapped genome-wide, providing unprecedented coverage of the epigenome and transcriptome across both haplotypes.

Because HG002 cells are openly available for sequencing and research (including iPSC lines), this comprehensive browser resource presents a unique resource for personalized genomics (**Figure S4**). Two companion studies demonstrate the new capabilities enabled by a personalized reference genome: Tullius *et al.* (to appear) apply Fiber-seq to HG002 sperm and lymphoblastoid cells to evaluate the nucleosome-to-protamine transition genome-wide, uncovering epigenetic features governing the intergenerational inheritance of the centromere kinetochore binding region as well as traces of regulatory information retained in the sperm epigenome; while Xu *et al.* (to appear) profile the chromatin architecture of HG002 centromeric satellites, revealing coordinated relationships between mCpG, H3K9me3, and CENP-A occupancy across centromere regions, as well as their epigenetic transmission through cell passaging. Such analyses would be impossible using GRCh38, which is an incomplete mosaic reference not linked to any one biological sample.

### Benchmarking genome assemblies

*De novo* assemblies of diploid human genomes using PacBio HiFi, ONT ultra-long, and Hi-C sequencing reads have begun to yield phased, chromosome-scale scaffolds of complete haplotypes^28,29,75^. Additionally, the development of pangenome databases^10^ has enabled genome inference methods that can predict personalized, haplotypic sequences even from short reads^76^. However, both *de novo* and inferred assemblies typically contain numerous structural errors like repeat expansions or collapses, misjoins, inversions, haplotype switches, and smaller substitution or insertion/deletion (indel) errors^77^. Since the T2T-HG002 assembly is complete and achieves nearly perfect accuracy, it can be used as a genome benchmark to identify likely errors in test assemblies generated by different sequencing and assembly approaches^78,79^.

Errors in alternative assemblies of HG002 can be directly identified by aligning each haplotype to its corresponding location in the genome benchmark and cataloging the differences. However, previously developed tools for assembly quality assessment (e.g. QUAST^80^ and GenomeQC^81^) do not consider the case of a diploid reference genome, and others (e.g. CRAQ^82^) only evaluate read alignments and not how well an assembly reproduces a genome sequence. We developed the Genome Quality Checker (GQC) software to enable comprehensive evaluation of DNA sequences against a highly accurate, diploid genome benchmark. GQC implements a haplotype-aware alignment strategy to collect statistics on continuity, accuracy, as well as phase consistency. Briefly, haplotype-specific *k*-mers are extracted from the diploid benchmark genome and a two-state hidden Markov model (HMM) over these markers is used to determine haplotype blocks in the test assembly (**Supplementary Information**). This information is used to refine the whole-genome alignments and differentiate phasing errors from base calling errors, which is especially helpful when evaluating pseudohaplotypes (i.e. assembled contigs that have not been fully phased). Because every error is assigned a position on the reference genome, all analyses can be stratified by a list of genomic regions to include or exclude.

Using GQC, we evaluated five previous assemblies of HG002 spanning five years of technology improvement against the T2T-HG002 genome benchmark (**Table S12 and Table S13**): the “Ash1v2” haploid reference assembly from 2020^53^, a diploid assembly from the initial “Hifiasm” publication in 2021^54^, a diploid assembly from the “HPRCv1” pangenome release in 2022^10^, a diploid assembly from the initial “Verkko” publication in 2023^28^, and a diploid “ONT LC24” assembly released by ONT at the London Calling conference in 2024 (https://epi2me.nanoporetech.com/lc2024_t2t/). The Jarvis *et al.* assembly^40^ represents an early iteration of the HPRCv1 assembly with slightly worse accuracy and so is included only in the supplementary tables.

Across several measures of completeness and accuracy, the assemblies showed progressive improvement over time (**Supplementary Information**). Specifically, NGAx plots show steady increases in the lengths of long scaffolds that are continuously alignable to the genome benchmark (**Figure 3A**). NGAx is especially powerful in conjunction with an accurate reference since it plots the size-ordered lengths of assembly-to-benchmark aligned blocks versus the percent of the benchmark covered, and, therefore, reflects both the continuity and structural accuracy of the test assembly^79^. Additionally, rates of substitution errors within alignable sequence have fallen over time by more than two orders of magnitude, and indel rates have fallen roughly tenfold (**Figure 3B,C**). There have been more modest improvements in the accuracy of mononucleotide run lengths (**Figure 3D**), with all assemblies continuing to struggle with long homopolymers, especially those greater than 20 base pairs in length.

**Figure 3:**
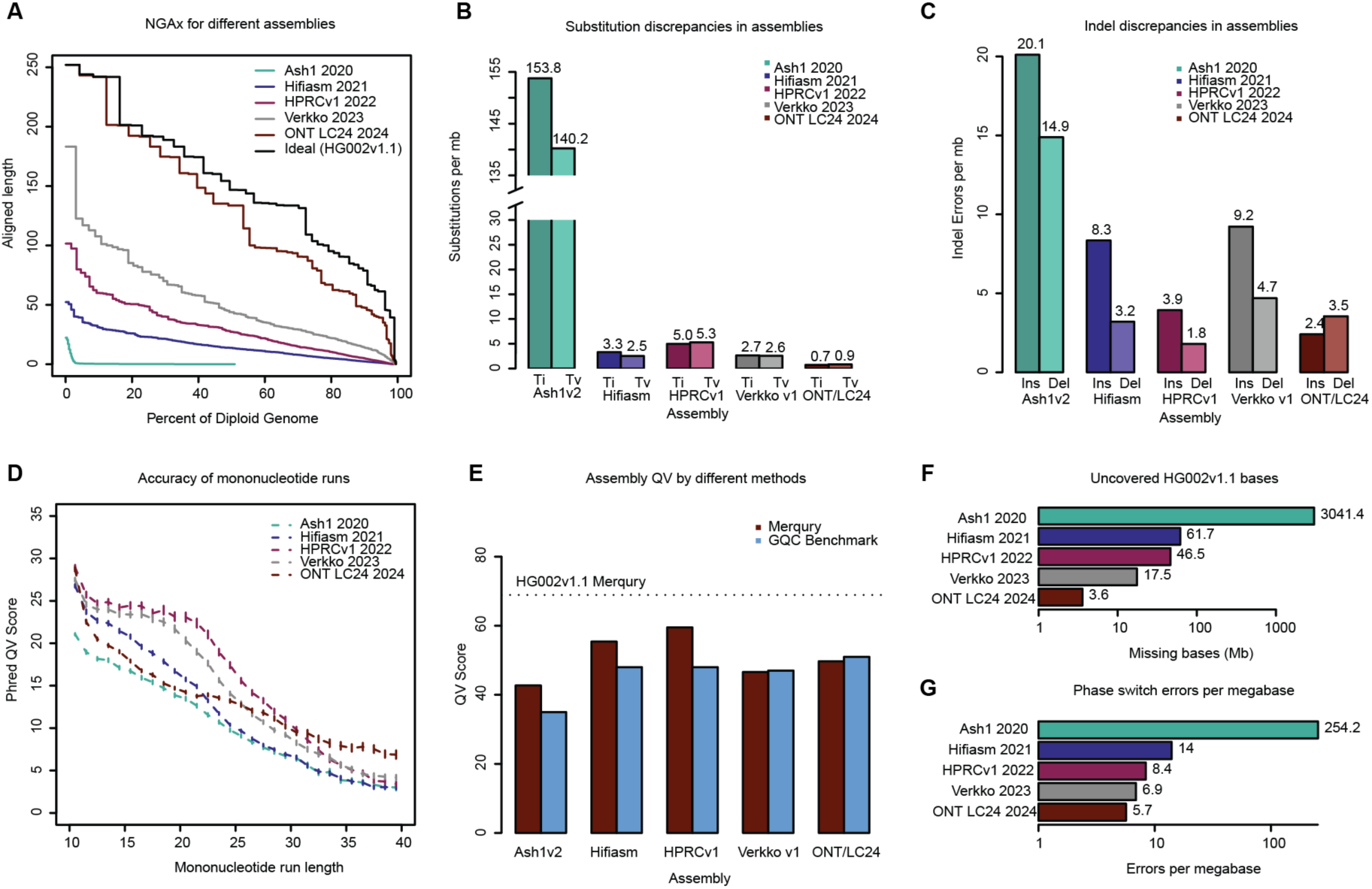
Five assemblies evaluated against the genome benchmark. **(A)** NGAx plot calculated from the lengths of uninterrupted alignments to the T2T-HG002 benchmark. Alignments were broken at locations of indels that were at least 10 kb in length. **(B)** Within-alignment substitution rates plotted with transition rates (Ti) to the left and transversion rates (Tv) to the right. Substitution alleles that match the alternate haplotype at heterozygous sites are not included in these counts, but are instead classified as phase switch errors. **(C)** Within-alignment indel rates, plotted with insertion rates (Ins) to the left and deletion rates (Del) to the right. Alleles that match the alternate haplotype at heterozygous sites are not included. **(D)** Mononucleotide run accuracy is measured by alignments to mononucleotide runs of different lengths in the benchmark and reported as a Phred-scaled quality score: −10*log_10_(#erroneous runs / #aligned runs). **(E)** Phred-scaled quality scores calculated using *k*-mer-based (Merqury) and alignment-based (GQC) methods: −10*log_10_(#discrepancies / #aligned bases). **(F)** Number of non-N bases in the genome benchmark uncovered by primary alignments of the test assemblies to the benchmark. **(G)** GQC phase switch rates within alignments of the test assemblies to the benchmark.

In addition to structural validation, our genome benchmark also measures the precise base accuracy of an assembly without the limitations of *k*-mer-based approaches such as Merqury. These methods are typically based on the detection of “error *k*-mers”, i.e. short sequences that are present only in the tested assembly and not in the raw sequencing data. Comparing base quality measured by GQC and Merqury across the same five assemblies revealed several interesting trends (**Figure 3E**). Using a combined database of 31-mers derived from the HiFi and Element reads, Merqury tended to overestimate the quality of the PacBio assemblies compared to GQC (Δ Ash1+7.9QV, Hifiasm+7.9QV, and HPRC+11.7QV), likely due to a confirmation bias towards the PacBio data and a failure to penalize missing or collapsed sequences. Conversely, assemblies that included ONT data showed a reduced Merqury quality score but an improved GQC score (Δ Verkko-0.4QV, LC-1.4QV), likely due to regions of the genome recovered only by the ONT data and possibly underrepresented in the HiFi and Element reads. This conclusion is supported by the increased completeness of the ONT-based assemblies versus the PacBio assemblies, which suffer from intermittent coverage dropout in GA-rich regions^83^ (**Figure 3F**). Lastly, phase switch errors, as identified by GQC’s HMM method, have also steadily decreased, with the ONT-including assemblies showing the best performance due to their increased read lengths (**Figure 3G**). These results highlight the advantages of a complete genome benchmark and suggest caution when interpreting *k*-mer-based quality scores alone.

### Benchmarking sequencing reads

Recent improvements in genome assembly quality are largely a consequence of ongoing improvements to the length and accuracy of sequencing reads. However, it is not sufficient to rely on the quality values reported by sequencing instruments themselves. Even if reported quality values could be assumed to be accurate, they do not capture other important quality information such as coverage biases (e.g. GC-rich or GA-rich sequence), error biases (e.g. difficulty with homopolymer runs), rates of chimeric reads, or the frequency of different types of errors (e.g. insertions, deletions, or substitutions). A genome benchmark enables us to comprehensively evaluate sequence reads by comparing them directly to the genome from which they originated and ensures that quality is measured across a broad range of metrics and sequence contexts.

Given BAM-formatted reads aligned to their genome of origin, GQC reports total aligned and clipped bases in the reads, as well as rates per megabase of read substitution and indel error within the primary alignments. It further stratifies error rates in homopolymer, dinucleotide, trinucleotide, and tetranucleotide runs of different lengths. Substitution error rates are reported for each of the twelve possible single-base errors, and if base quality scores are included in the BAM file, tallies of substitution and indel errors are reported for each quality score value.

Comparing four recent datasets with GQC shows marked differences in base calling performance and coverage uniformity (ONT “Q28” Ultra-Long Sequencing Kit V14; PacBio Revio HiFi ICSv13 with DeepConsensus v1.2; Element 2×150 Aviti “Q50” UltraQ from 400 bp inserts; and Illumina 2×151 NovaSeq PCR-Free from 300 bp inserts) (**Table S12 and Table S14**). Substitution error rates show that Illumina short reads have five times the rate of transversion errors as the other platforms^84^ (**Figure 4A**), while HiFi and ONT long reads have indel error rates two orders of magnitude higher than the short reads (**Figure 4B**). For homopolymers, Element reads show an order of magnitude lower error rate than all other platforms (**Figure 4C**). It is also possible to compare observed versus reported quality values for each technology by binning their bases by quality score and calculating error rates from the alignments. While short read (Illumina, Element) base quality scores are well calibrated, base calls in the long reads (HiFi, ONT) show lower-than-reported accuracy, on average, for bases with reported quality scores greater than Q20 (**Figure 4D**). Analysis of the distribution of binned coverage shows that all technologies show a wider distribution than the expected Poisson (**Figure 4E)** and that while HiFi Revio is more overdispersed for coverage values lower than the median, Illumina PCR-Free is more overdispersed for higher coverage values. GQC can also report windowed read coverage as a function of GC content, which shows the long reads outperforming short reads for high %AT, and all technologies struggling with high %GC (**Figure 4F**).

**Figure 4:**
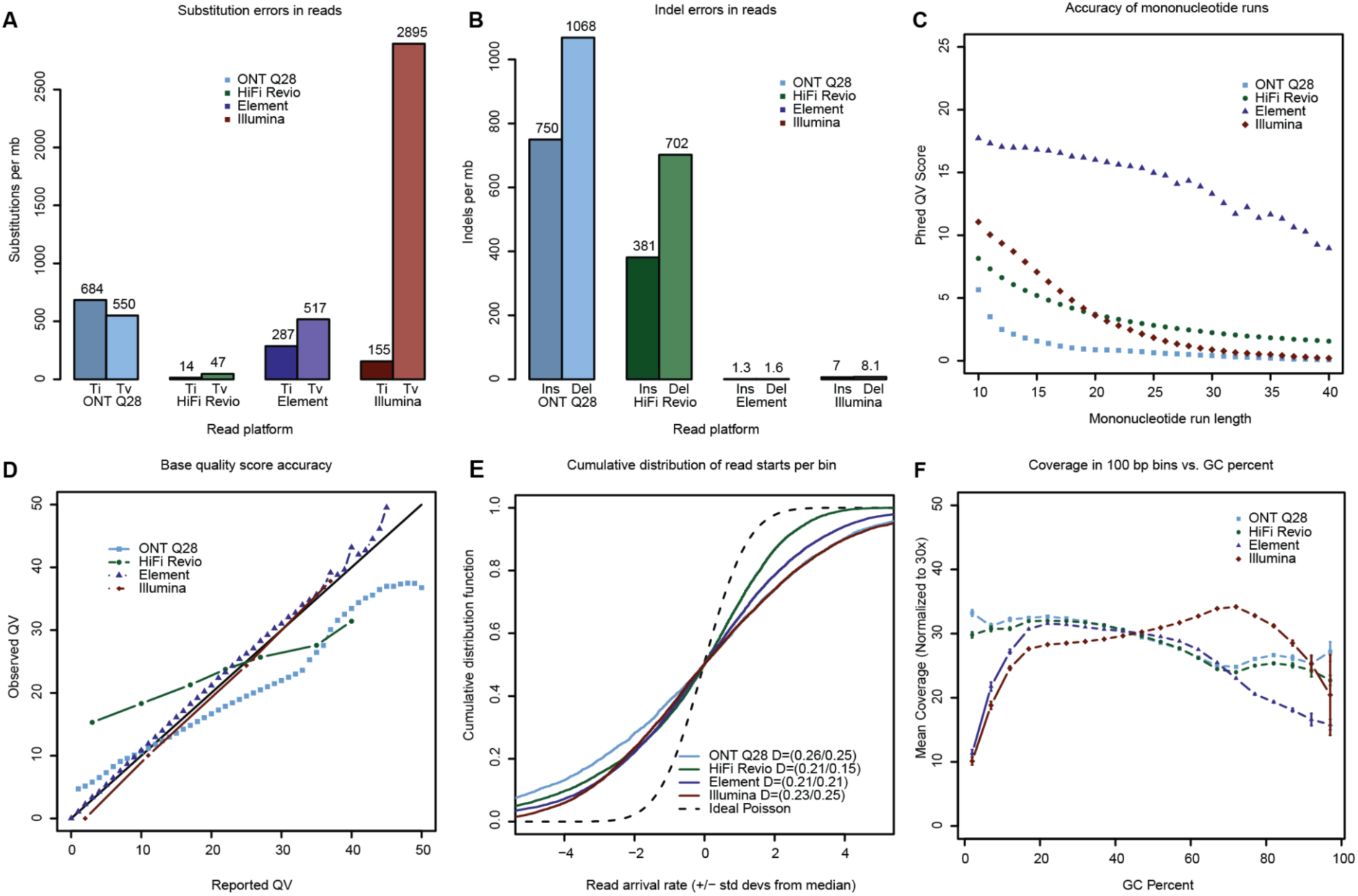
Four sequencing technologies evaluated against the genome benchmark. **(A)** Read substitution rates within alignments of reads to the genome benchmark. “Ti” is the count of transitions and “Tv” is the count of transversions. **(B)** Insertion and deletion rates within alignments of reads to the genome benchmark. **(C)** Phred-scaled quality scores for homopolymer runs of different lengths. To be considered correct, a homopolymer must be the correct length and free of single-nucleotide errors both within the repeat and in the 5 flanking bases before and after the repeat. **(D)** Accuracy of base quality scores calculated from actual alignment match rates (Observed) versus base quality scores reported with the reads (Reported). **(E)** Cumulative distribution of read starts per bin (arrival rate) for different technologies scaled by the expected standard deviation for a Poisson distribution and shifted to put the median at zero for visualization. Kolmogorov-Smirnov distances (D values) correspond to the maximum difference between a curve and the ideal below and above the median, respectively. **(F)** Mean coverage in 100 bp windows as a function of %GC.

### Benchmarking variant call sets

Variant call sets are by far the most commonly benchmarked genomic data type, but do not typically assess the entire genome. Conveniently, it is possible to convert variant sets to genomes and vice versa to better understand their limitations. Tools such as dipcall^15^ and PAV^85^ report variants from the alignment of a diploid assembly to a reference, which can then be compared to a variant benchmark to measure assembly consensus accuracy^86^. Conversely, it is possible to apply a set of variants to a reference genome using BCFtools^87^ and in the process build a “variant constructed genome” by updating a reference genome sequence with the alleles reported in a VCF file. Ideally, a perfect variant set would produce a variant constructed genome which perfectly matches the sample being sequenced, but in practice the output remains limited due to the incompleteness of the reference and/or variant calling.

To measure how much of the genome is being overlooked by state-of-the-art variant calling approaches, we generated multiple HG002 variant constructed genomes against both GRCh38 and T2T-CHM13 references. Since a genome benchmark consists of complete haplotypes, the variant callsets must be at least partially phased for evaluation, and it is necessary to know which regions of the reference genome have been sufficiently covered by the sequencing data, as provided by the gVCF format^88^. Using the best performing single-technology approach^3^, we variant called HiFi long reads with DeepVariant^57^ and phased with HapCut2^89^ using the same data. We then generated diploid variant constructed genomes by applying the called variants from each haplotype to the reference and masked bases that were below genotype quality (GQ) score thresholds of 10, 20, 30, and 40 (**Supplementary Information**). This resulted in eight constructed genomes (two references × four GQ thresholds) which we evaluated by GQC to assess quality and completeness (**Table S13**).

The variant constructed genomes built using GRCh38 cover 93.8–92.9% of the HG002 genome (min GQ 10–40). In contrast, those built using CHM13 cover 97.8–95.7% (**Figure S5**), demonstrating the improved coverage of variant call sets produced by mapping against a complete reference sequence. However, many of these added bases are in difficult-to-call regions of the genome, such as satellite repeats, and the average quality of the consensus sequence suffers slightly as a result. Average consensus QV ranged from 37–40 for the GRCh38-variant constructed genomes (min GQ 10–40), and from 35–39 for the CHM13-variant constructed genomes. In particular, there was a higher rate of single-nucleotide substitution errors when using the CHM13 reference compared to GRCh38 (72.3 / Mb vs. 41.1 / Mb for GQ ≥ 40), while the indel rate was comparable (52.6 / Mb vs. 50.6 / Mb, respectively) (Figure S5). When restricted to only parts of the HG002 genome covered by variant constructed genomes from both references (thus, eliminating most satellites), average QV ranged from 37–41 for the GRCh38-variant constructed genome and 38–41 for the CHM13-variant constructed genome (**Table S13**). In comparison, the best-performing historical HG002 assembly evaluated here, ONT LC24, achieves a QV of 51 with coverage 99.94%. Thus, when averaged across the whole genome, *de novo* assembly can now outperform variant calling accuracy by an order of magnitude and achieve higher genome coverage than mapping to T2T-CHM13.

### Relationship between genome and variant benchmarking

Due to their increased coverage and more natural representation of complex genomic regions, genome benchmarks can also be used to improve variant benchmarks. The first T2T assembly of HG002’s X and Y chromosomes^24^ was used to create phased variant benchmarks by aligning it against GRCh37, GRCh38, and T2T-CHM13v2.0 references to call variants in complex regions and exclude regions of ambiguous orthology^16^. The resulting updated GIAB variant benchmarks are more accurate and comprehensive, enabling the evaluation of sex chromosomes, complex structural variants, large indels, indels within long homopolymers and tandem repeats, and variants within closely related segmental duplications for the first time.

To assess what types of errors may be missed by variant benchmarks and facilitate a transition from variant-centric to genome-centric analysis, we analyzed the relationship between genome benchmarking metrics (e.g. consensus and phasing accuracy) and standard variant benchmarking metrics (e.g. precision, recall, F1, false positives (FPs), false negatives (FNs), and genotype errors). As a case study, we evaluated the previous Jarvis *et al.* diploid assembly of HG002^40^ using both genome and variant benchmarking and compared the errors identified (**Table S15**). To focus our comparison on differences between the two approaches, rather than the underlying benchmarks, we used an updated version of the GIAB variant benchmark that was created by aligning the T2T-HG002 assembly to GRCh38 (https://ftp-trace.ncbi.nlm.nih.gov/giab/ftp/data/AshkenazimTrio/analysis/NIST_HG002_DraftBenchmark_defrabbV0.019-20241113/). Single nucleotide variant (SNV) and indel designations for genome benchmarking were based on comparison of the HPRCv1 assembly directly to T2T-HG002, while variant benchmarking was based on alignment of HPRCv1 to GRCh38 and comparison to the variant benchmark.

Genome benchmarking identified 66,544 errors in the HPRCv1 assembly, of which 50,843 could be lifted to GRCh38 and 30,505 fell within the small variant benchmark’s high confidence regions. Thus, more than half of the total genome benchmarking errors in the HPRCv1 assembly were not assessed by variant benchmarking due to reference bias and a lack of benchmark coverage. Restricting the comparison to only the high confidence regions, FP and FN rates from variant benchmarking largely correlated with error rates from genome benchmarking across various repeat stratifications (**Figure 5A, Figure S6**). For these regions, variant benchmarking reported more errors (sum of FP and FN) than genome benchmarking, except for SNVs within segmental duplications, where the two methods reported a similar frequency of error. We found that each genome benchmarking error often translated into multiple variant benchmarking errors due to complex variants in homopolymers and tandem repeats or differences in how phasing errors were reported, which explained the higher rate of errors from variant benchmarking (**Supplementary Information, Figures S6, S7, S8, S9, and S10**). In this way, genome benchmarking provides a more straightforward interpretation of error without the added complexity of natural variation between the sample and reference. We found that phase switch errors (swapped haplotypes) and haplotype collapse errors (loss of heterozygosity) were often more interpretable in the variant benchmarking results, but this limitation could be addressed by future genome benchmark tooling.

**Figure 5:**
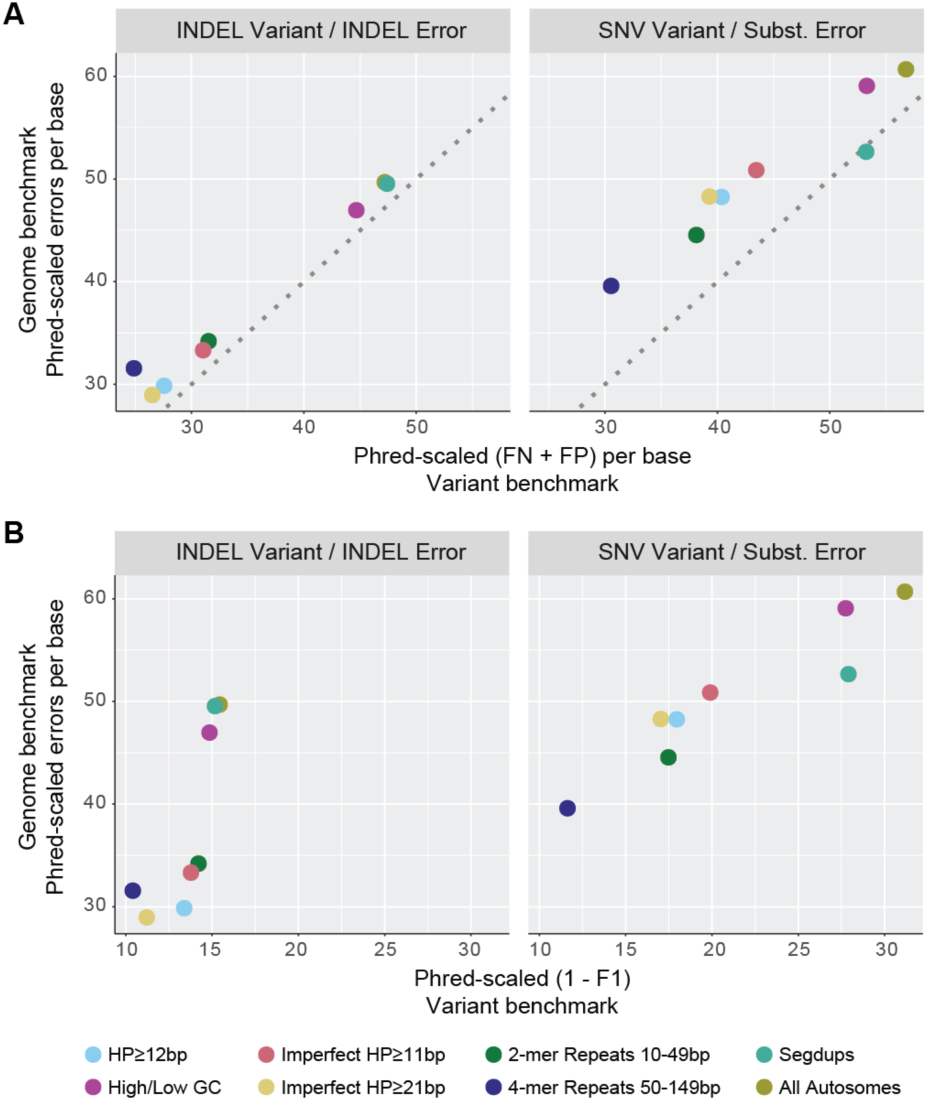
Comparison between variant and genome benchmarking results. Indel and SNV errors were identified in the Jarvis et al. assembly of HG002 using both the GIAB HG002 variant benchmark and T2T-HG002 genome benchmark, with results stratified by various sequence contexts (e.g. light blue = homopolymers ≥12 bp). For comparison, genome benchmarking errors were restricted to those which could be successfully projected onto GRCh38 GIAB small variant confident regions (30,505 of 66,544 genome benchmarking errors). **(A)** SNV and INDEL variant benchmarking false negatives and false positives per base of GIAB confident regions vs. substitution and INDEL genome benchmarking errors per base in the Jarvis et al. assembly (normalized by the number of bases in each stratification). Error rates are plotted as Phred-scaled quality: −10*log_10_(P_error_). Points above the dotted line indicate contexts for which variant benchmarking reported more errors than genome benchmarking. **(B)** Variant benchmarking F1 score vs. genome benchmarking errors per base. F1 scores are not directly relatable to per-base error rates because they are a fraction of true variants rather than a fraction of bases, but the Phred-scaled (1−F1) is shown on the same log scale to depict the relationship of these commonly-used performance metrics. Individual plots of FP, FN, precision, and recall are shown in **Figure S6**.

Genome benchmarking error (or consensus accuracy) is not directly comparable to more commonly used benchmarking metrics like precision and recall, because these metrics use different denominators. Whereas genome benchmarking measures accuracy at every base in the genome, precision/recall only measure known variant sites. For example, variant-based indel F1 scores do not necessarily correlate with genome benchmarking error rates because most true indels in the variant benchmark fall within homopolymers and tandem repeats (due to their naturally higher mutation rate). However, such repeats make up only a small fraction of the total genome, so their stratified error rate is elevated in genome benchmarking compared to the genome-wide average (**Figure 5B, Figure S6A**). These comparisons point to the need for standardized genome benchmarking metrics and stratifications to enable downstream users to interpret performance in a familiar context. One compromise could be to identify a set of clinically important or known variant sites in the diploid HG002 genome and report precision/recall measurements across these positions, which would be akin to variant benchmarking but without the confounding reference bias.

## Discussion

Building on the widely used GIAB HG002 reference material, T2T-HG002v1.1 establishes the most comprehensive genomics benchmark ever constructed. Compared to CHM13, HG002 is more suitable as a benchmarking genome due to its availability as a NIST reference material^42^, previously established variant benchmarks^1,18^, commercially available LCL and iPSC lines from the NIGMS Human Genetic Cell Repository at Coriell^90^, and consent for open data sharing^39^. This makes HG002 an ideal testbed for developing personalized genomics, where an individual’s complete, T2T diploid genome serves as the basis for analyses. Towards this goal, the T2T-HG002 genome benchmark will be essential for measuring progress and is already being used to guide the development of new genome technologies such as sequencing^91^, polishing^48^, and variant calling methods^13,92,93^. We also anticipate the development of functional genomics benchmarks for RNA-seq, methylation, chromatin and other assays to be anchored on the resource presented here.

No genome assembly is infallible, and this includes T2T-HG002v1.1. The “Q100” project name invokes an aspirational goal of creating a genome benchmark with a Phred quality score of Q100, corresponding to a virtually error-free 6 Gbp diploid human genome. However, this level of accuracy is exceedingly difficult to achieve and validate. First, the majority of sequencing data generated here was derived from cultured HG002 LCL cells, and some degree of somatic variation is expected relative to the NIST reference material DNA that was extracted from the same cell line but from a different batch of cells^42^. Even within the reference material itself, 85 high-confidence mosaic SNVs have been previously identified with variant allele fractions between 5–30%^94^. Second, benchmarking always suffers from a bootstrapping problem, as it is difficult to validate newly resolved regions of the genome that are not accessible to all sequencing technologies. Here we show that *k-*mer-based methods for QV estimation miss certain classes of errors (e.g. loss of heterozygosity, long homopolymer errors, repeat expansions/collapses) and can report false positive errors if the *k*-mer database is incomplete (e.g. due to coverage bias in the raw sequencing data). Although it is possible to artificially increase *k*-mer quality by removing the known erroneous *k*-mers, this risks removing true heterozygous or paralog-specific variants. Thus, to construct a reliable benchmark, we chose a more conservative polishing approach that based decisions on an understanding of the errors being corrected and only used the *k*-mer-based methods to measure progress. In fact, most of the corrections we made had little impact on *k*-mer-based QV estimates due to their limitations in dealing with repetitive regions and heterozygous variation. The remaining regions of lower confidence include long homopolymers (i.e. >20 bp in length) and repetitive regions of the genome that could not be reliably mapped and validated with short reads (e.g. rDNAs, satellites, and segmental duplications). Given the size and potentially high mitotic recombination rate of rDNAs^95^, further improvements in sequencing technology are likely needed for their confident reconstruction. Excluding rDNA arrays, 99.9% of the diploid genome is covered by high-confidence genome benchmark regions in v1.1, compared to 85.2% with the v4.2.1 variant benchmark (**Table S16)**.

We have demonstrated that prior variant-based benchmarks are limited in scope, prone to reference bias, and leave large parts of the genome unrepresented. Additionally, variant benchmarking stratifies errors into categories (SNVs, indels, complex variants) based on their representation *relative to the reference*, which can obscure the true nature of the errors, particularly in repetitive regions where a single error can manifest as multiple variant calls of different types. Genome benchmarking fundamentally simplifies this problem and isolates the errors from any natural differences between the sample and reference. Ideally, all GIAB variant benchmarks can be updated and improved with a corresponding genome benchmark. However, there is a near-term need for improved variant benchmarks since variant calling remains the primary output of most genomics workflows, especially in clinical settings where the endpoint is a set of variants annotated for their clinical significance and the performance metrics are similar to those used for clinical testing. As methods to interpret the clinical significance of personal genome assemblies are developed, new approaches will be needed to understand how genome benchmarking performance metrics impact clinical interpretation. This human genome benchmark will be used to improve existing variant benchmarks, particularly in highly repetitive and copy number variable regions of the genome, and help guide the formulation and interpretation of genome benchmarking metrics, which are not yet standardized.

T2T genome sequencing and assembly methods have enabled discoveries within previously hard-to-analyze regions of the genome, such as segmentally duplicated genes^25,61^, amplified gene families^96^, satellite DNAs^97,98^, and tandem repeats^99,100^. Continued progress towards the routine sequencing and analysis of complete genomes is necessary to fully illuminate the dark genome across population scales^76^. In the near term, such analyses would require a high-quality pangenome^41^ comprising a large number of samples from which the entire genome of an individual could be inferred using less expensive sequencing methods^101–103^. However, the rigorous validation of such pangenome-based methods has been impossible without a ground truth that encompasses the entire genome. The complete, diploid T2T-HG002 genome benchmark provides this missing validation framework, removes the performance ceiling of previous variant benchmarks, and will drive the development of new methods for low-cost, complete genome inference.

We expect the routine inference of T2T genomes to mark a new era of personalized genomics, whereby all analyses are performed in the context of an individual’s complete, diploid genome. In this way, an individual’s somatic and functional genomics data can be unambiguously mapped to the genome it was derived from, eliminating reference bias. An open collection of T2T genomes with matched, long-read functional data would allow for the training of sequence-based computational models of the entire genome that, combined with existing knowledgebases and literature, could be used to annotate rich metadata such as gene annotations, regulatory elements, population allele frequencies, and disease risk predictions directly onto the personalized genome. This would ultimately allow clinical genome assessment to move beyond the interpretation of individual variants and towards the interpretation of whole genomes.

## Resource Availability

### Lead contact

Requests for further information and resources should be directed to and will be fulfilled by the lead contact, Adam M. Phillippy.

### Materials Availability

No new materials were generated from this study. The sample is publicly available as NIST Reference Material 8391, Coriell lymphoblastoid cell line (LCL) GM24385, PGP participant huAA53E0.

### Data and Code Availability

The T2T-HG002v1.1 assembly is available from NCBI GenBank under the HPRC BioProject PRJNA730823 with accession numbers GCA_018852605.3 (paternal) and GCA_018852615.3 (maternal). No new sequencing data was generated for this project, but all data used here is organized in **Table S1** and available from the project GitHub along with links to the browser hub, issue tracker, and associated analysis software (https://github.com/marbl/hg002). Any additional information required to reanalyze the data reported in this paper is available from the lead contact upon request.

## Acknowledgements

This work was supported, in part, by the Intramural Research Program of the US National Human Genome Research Institute, National Institutes of Health (NFH, SK, AR, DA, JK, BDP, SJS, APS, and AMP). The contributions of NIH authors are considered Works of the United States Government. The findings and conclusions presented in this paper are those of the authors and do not necessarily reflect the views of the NIH or the U.S. Department of Health and Human Services. This work is also supported, in part, by NIST Intramural Funding (ND, NDO, JW, and JMZ). Certain commercial equipment, instruments, or materials are identified to specify adequately experimental conditions or reported results. Such identification does not imply recommendation or endorsement by the National Institute of Standards and Technology, nor does it imply that the equipment, instruments, or materials identified are necessarily the best available for the purpose. Several authors are supported by NIH grants 5R01HG010169 (to EEE), R00GM147352 (to GAL), 2T32GM007454, 1K99GM155552, and 1U01HG013744 (to MRV), R01-HG006677 and R35-GM156470 (to K-HC), UM1DA058229 (to LFP), R01-HG006677 and R35-GM130151 (to SLS), 1UG3NS132105 and 1U01HG011758 (to FJS), U24HG010263 (to MCS), 1DP5OD029630, UM1DA058220, and 1U01HG013744 (to ABS), and UM1HG010971 and R01HG011274 (to KHM), as well as USDA NIFA 2023-67015-39261 (to JK and RDS) and HATCH MO HAAS0001 (to RDS). NA is a Chan Zuckerberg Biohub Investigator and is supported by an HHMI Hanna H. Gray Fellowship and a Pew Biomedical Scholar Award. RJO is funded by the John and Donna Krinicki Endowment and by the Colossal Foundation. EEE is an investigator of the Howard Hughes Medical Institute. KHM was supported by the Searle Scholars Program. This work utilized the computational resources of the NIH HPC Biowulf cluster (http://hpc.nih.gov).

## Author Contributions

Assembly generation: NFH*, AR*, DA, MR, SK*, Curation and validation: NFH*, ND*, AR*, HL*, GAL*, EA, DA, MA, SB, SCB, AVB, SAC, AC, YC, AD, PE, MF, LEF, GF, AG, KJ, YK, JKim, RK, JL, MM, SN, KKO, LFP, BDP, DP, DR, HR, FJS, KS, MS, SJS, APS, JW, SK*, JMZ, AMP. Annotation: NFH*, HJJ*, AR*, HL*, MRV*, JMS*, EA, NA, DA, KHC, GAH, JR, DR, HR, WT, DY, YZ, ABS, RJO, KHM, SLS. Cytogenetics: MWM. Variant benchmark: ND*, AE, NDO, JW, JMZ. Software: NFH*. Analysis: NFH*, ND*, AR*, JK, JKim, PE, KS, SLS, JMZ, AMP. Manuscript figures: NFH*, ND*, JKim, AMP*. Manuscript writing: NFH*, ND*, HJJ*, AR, GF, AG, JK, RDS, EG, EEE, MCS, RJO, SLS, KHM, SK*, AMP, JMZ. * denotes group leaders: SK assembly generation, NFH curation, ND variant benchmark, HJJ gene annotation, AR validation, HL satellite annotation, GAL centromere validation and annotation, MRV Fiber-seq annotation, JMS repeat annotation. All authors read and approved the final manuscript.

