## Supplementary Material for "A complete diploid human genome benchmark for personalized genomics"


Sequencing and initial assembly of HG002 3
- HG002 samples used for this project 3
- Sample G-banded karyotyping of the GM24385 cell line 3
- Sequencing and initial assembly of HG002 (v0.7) 3
- Strand-seq-based clustering of acrocentric chromosomes 6

Polishing and patching of T2T-HG002v0.7 to create T2T-HG002v1.1 6
- Crowd-sourced curation 6
- Read alignments 7
- Categories of issues/corrections to v0.7 (round 1) 7
  - Small corrections 7
    - Phase-switched and falsely homozygous sites 7
    - Small errors discovered from short-read homozygous calls 9
  - Corrections to larger regions 12
    - T2T-Polish-discovered issues 12
    - Issues discovered with Flagger/SecPhase 13
    - Patches to problematic regions using a newer assembly (HG002v0.8) 14
  - Final assembly corrections applied to v0.7 14
- Categories of issues/corrections to v0.9 (round 2) 15
  - Corrections to problem areas in “difficult” regions of T2T-HG002v0.9 15
    - Correction of errors identified in centromeres 15
    - Correction of errors identified in telomeres 16
    - Extension of v0.9 rDNA flanking sequences into the rDNA gap 17
    - Lengthening of paternal Chr13’s rDNA array using spanning ONT reads 19
  - Correction of small errors using trio-based Onso, standard and long-insert Element, Illumina, and HiFi DeepVariant calls 19
  - Final assembly corrections applied to v0.9 20
- Categories of issues and corrections to v1.0.1 (round 3) 21
  - Patching v1.0.1 with consensus from two alternative assemblies 21
  - Polishing and patching v1.0.1 to create v1.1 22
  - Methods used to discover regions for attempted patching 22
- Total corrections made in three rounds of polishing 25

Evaluation of T2T-HG002v1.1 25
- Strand-seq assembly evaluation 25
- k-mer based evaluation 26
- Mapping based evaluation 28
  - T2T-Polish issues 29
  - NucFlag 29

Visualization of short-read alignments 30
Prior known issues 31
Consolidated Issues Track 31
Annotation and browser resources for T2T-HG002v1.1 32
Gene annotation 32
Identification of haplotype-specific genes 34
FIRE analysis of Fiber-seq data for HG002 35
Repeats and transposable element sequences 35
Centromeric satellite sequences 36
Segmental duplications 37
Subtelomeric regions 37
Representation of additional HG002 rDNA units 38
Creation of a T2T-HG002v1.1 ideogram track 38
Heterozygous variants and estimated windowed heterozygosity 39
Chain files to and from the HG002v1.1 assembly 40
Fraction of HG002v1.1 inaccessible to variant-based benchmarks 41
GIAB v4.2.1 missed regions in various annotated regions of HG002v1.1 41
Use of HG002v1.1 as a genome benchmark to evaluate assemblies, reads and variant call sets 42
Evaluation of test assemblies using GQC 42
Pre-phasing diploid assemblies with GQC 42
Evaluation of assembly's long-range continuity 43
Evaluation of assembly accuracy within alignments to the benchmark 43
Evaluation of mononucleotide run accuracy 43
Assemblies used for comparison and evaluation with HG002v1.1 44
Evaluation of sequencing read accuracy using GQC 44
Datasets used to demonstrate read accuracy benchmarking 44
Calculation of substitution and small insertion/deletion discrepancies 45
Coverage assessment 45
Assessment of short tandem repeat accuracy 45
Evaluation of base quality scores 46
Evaluation of variant call sets via "constructed genomes" 46
Converting gVCF files to VCF files with high confidence region BED files 46
Phasing variant call sets 47
Creating and benchmarking a variant-constructed genome 47
Direct comparison between genome-based and variant-based benchmarking errors 49
Overall comparisons between genome benchmarking QV and variant benchmarking precision/recall 50
Equivalencies between variant benchmarking errors and genome benchmarking errors 52
Categories of variants and their relation to the genome errors provided by GQC 52

### Sequencing and initial assembly of HG002

#### HG002 samples used for this project

##### Sample G-banded karyotyping of the GM24385 cell line

The lymphoblastoid cell line GM24385 was obtained from the NIGMS Human Genetic Cell Repository at the Coriell Institute for Medical Research and expanded to a total culture size of $2x10^{10}$ cells to create the HG002 lot of cells housed at NIST[42]. G-banded karyotype analysis was performed on this expansion of cells, harvested at passage 2 (post-cell line establishment). During the g-banding analysis, 73 metaphase cells were counted, and 12 metaphase cells were analyzed and karyotyped. Chromosome analysis was performed at a resolution of 400 bands or greater. DNA used for microarray was isolated from a frozen cell pellet (2x107 cells) using the Autopure LS instrument (Qiagen). DNA was genotyped using the Affymetrix Human SNP Array 6.0 (Affymetrix, Inc.).

Cytogenomic analysis of the g-banding and microarray results showed a karyotype of 92,XXYY[13]/46,XY,?inv(3)(q26.3q29)[3]/46,XY[57].arr[hg19](1-22)x2,(XY)x1 in this expansion lot of GM24385/HG002 cells (**Figure S1**). While the microarray analysis resulted in a normal, human male karyotype, the extensive g-banding data show a mostly diploid (78%) (**Figure S1A**), with some tetraploid cells (18%) (**Figure S1B**), and a very small number (4%) of cells showing an inversion on chromosome 3 (**Figure S1C**).

##### Sequencing and initial assembly of HG002 (v0.7)

The assembly, polishing, patching, and validation of the HG002 genome made use of sequencing data from short, long, and ultra-long datasets from the Human Pangenome Reference Consortium (HPRC)[41] and Genome in a Bottle Consortium (GIAB)[42] (**Table S1**). In addition to 170x coverage in HiFi Sequel II reads and 166x of ONT reads longer than 100kb used for the first assembly, roughly 200x coverage of HiFi Revio reads called with DeepConsensus v1.2 became available and were utilized during the polishing stage of the project. 82x coverage of R10 ONT reads greater than 100kb in length were also used for polishing.

The initial assembly of HG002 used for this manuscript, labeled "v0.7", consists of a manually-curated set of scaffolds derived from verkko version 1.0 (commit f69fa7c0f62d4f1eaf2288a22c7e9e58fd1cacd7) assemblies (v0.1 through v0.6) run during the summer of 2022[28]. Roughly 170x coverage in HiFi Sequel II reads and 166x of ONT reads >= 100 kb were assembled, using short read k-mer "hapmers" determined using Illumina sequence (**Table S1**) from HG002, HG003, and HG004 (HG002 and its parents) for phasing. The v0.7 release was the last unpolished release in a series of curated verkko assemblies.

The initial assembly (assembly v0.1) had 25 T2T contigs, 28 T2T scaffolds, and 18 gaps. The initial QV was 63.1 measured with hybrid 31-mers (see Evaluation section). It was aligned to the

CHM13 T2T reference to assign chromosome names to the graph and assembled sequences. This assembly resolved Chromosomes 1 maternal, 2 maternal, 3 both, 4 both, 7 both, 8 both, 9 both, 10 both, 11 both, 12 paternal, 16 both, 18 both, 19 both, 20 paternal, and Y. The three T2T scaffolds were Chr 1 paternal, Chr 12 maternal, and Chr 17 paternal with 1 gap each.

The assembly was resolved T2T based on manual evaluation and gap-filling following the protocols in Nurk et al[8]) and using the sg_sandbox repository archived in this manuscript's software repository[104]. Briefly, Chromosome X was re-used from the original HG002 ChrY publication[24]. ChrY from the publication was not used because it has a known inversion which the v0.1 assembly corrected. Chromosome 12 maternal was also previously resolved and re-used. The gap in Chromosome 1 paternal was due to an un-popped bubble and was resolved by inspection of the graph. Chromosome 13 paternal was resolved by scoring multiple traversals of a gap and selecting a winner as in Nurk et al. This assembly was named v0.1. Assembly v0.2 was identical to v0.1 except resolved chromosome orientations were updated to match the CHM13 T2T reference.

Chr 5 maternal, Chr 13 maternal q-arm, and Chr 20 maternal had overlaps between the contigs at the assembly breaks. To confirm these overlaps we generated an independent assembly using Flye[105]. All ONT reads > 100 kb were trio-binned using merqury[52] and assembled independently. In all cases, the Flye assembly confirmed the overlap size. This assembly after joining overlapping sequences was called v0.3.

Chromosome 14 paternal q-arm was resolved by comparing traversals through the graph and selecting the highest scoring one. Chromosomes 2 paternal also had a gap which was patched using a consensus from three ONT reads. The remaining chromosomes could not be resolved using this initial assembly graph. To resolve them, an additional assembly was generated using experimental MBG options (copycount_resolve commit 8e03e068ca64d986a9f983aab35ce4b1d4c4176, available as option --copycount-filter-heuristic since MBG v1.0.13) which performs extra simplification by removing low-coverage nodes during multiplex resolution. Chromosome 6 maternal and paternal were resolved based on the best-scoring traversal in the new assembly graph. Chromosome 17 maternal and paternal were patched to close gaps in the original assembly using the best-scoring traversals from the new assembly. This assembly was called v0.4.

Chromosome 5 paternal was resolved by taking the best traversal from the new assembly and lifting it back to the original graph with GraphAligner[106] alignment. The assembly was screened for Epstein-Barr virus (EBV) (AJ507799) sequences and mitochondrial human DNA (NC_012920). Any matching nodes were removed from the assembly to avoid redundancy. Representative mitochondrion and EBV sequences were selected by coverage and trimmed (if needed) to remove any redundant sequence on the ends. This assembly was called v0.5.

Nodes matching rDNA were identified using mash screen with the commands:

```
mash sketch -I unitigs.hpc.fasta -o sketch.msh
mash screen sketch.msh KY962518.fasta |awk '{if ($1 > 0.9 && $4 < 0.05) print $NF)' >
target.screennodes.out
```

where KY962518.fasta is the KY962518 reference rDNA sequence. The matching nodes were collapsed to a single node and telomere added to the graph to annotate ends of chromosomes. The script to perform this analysis (remove_nodes_add_telomere.py) is archived in this manuscript's software repository[104]. This simplified the graph for the acrocentric chromosomes, allowing the distal regions to be resolved. In all except two cases, the majority of the distal arm was resolved in a single node. Two paternal distal regions (later determined to be Chr 13 paternal and Chr 22 paternal, see below) were tangled together in the graph with approximately 1.6 Mbp of 1.8 Mbp (in homopolymer-compressed space) shared between the two chromosomes. The two best paths through this region based on ONT alignments were selected to represent the distal regions. This assembly was called v0.6.

Two independent methods were used to assign the acrocentric chromosome ends to their correct chromosome. Sequence for all ten chromosome ends was generated from the assembly graph by aligning the rDNA repeat unit and DJ sequence to identify paths. These were assigned to a parental haplotype using trio markers. The paths could be resolved unambiguously except in the case of two paternal haplotypes which were nearly identical with few bubbles. For these two (Chr 13 and Chr 22), two paths were generated, each using the shared paths and alternating in the bubble, but the phasing between bubbles is likely inaccurate. This resulted in 10 candidate distal regions, 5 for each haplotype. Then, pstools[107] was used to identify k-mer based mappings of Hi-C sequences to the paternal and maternal assembly and the respective distal regions, independently. For each possible pair of distal region and q-arm in a haplotype, the pair with the maximum read support as well as the second-best pair were reported. Independently, Strand-seq data (**Table S1**) was used to identify best-buddy matches between the ends and q-arms of the chromosomes, again within a haplotype (see "Strand-seq-based clustering of acrocentric chromosomes", below). The two methods agreed with the exception of Chr13/21 paternal where Hi-C swapped the assignments versus Strand-seq. The second-best matches in the Hi-C pstools results agreed with Strand-seq. FISH was used to identify characteristic WaluSat repeat arrays on the chromosome ends to disambiguate this assignment[46]. The Hi-C assignment would indicate there is a weak WaluSat signal present on both Chr21 and Chr22 paternal while Strand-seq indicates that it would be present on Chr13 and Chr22. The images better support the signal on Chr13 and Chr22 and thus the Strand-seq assignment for these chromosomes was selected. The presence of WaluSat on the other chromosomes was also confirmed to be consistent with the assignments using Hi-C and Strand-seq data (**Table S1**). The rDNA on Chr13 paternal was short enough to be manually resolved. For the other acrocentric chromosomes, gaps corresponding to the FISH-estimated size of their rDNA arrays[46] were inserted into the sequence, resulting in the v0.7 assembly that was released in November, 2022 and used as the starting assembly for the polishing described in "Polishing and patching of HG002v0.7 to create HG002v1.1".

#### Strand-seq-based clustering of acrocentric chromosomes

As reported before, Strand-seq has a unique ability to preserve the strand inheritance of the whole maternal and paternal homologs[50]. This information can be used to to assign assembly contigs to their respective homologous chromosomes. We used Strand-seq (**Table S1**) to assign short and long arms of all acrocentric chromosomes (13, 14, 15, 21, and 22) to their respective paternal and maternal homologs. To do this, we extracted the last 5% of Strand-seq reads at the end of the short acrocentric arm (tail) and first 5% of Strand-seq reads from the beginning (head) of the long acrocentric arm. Only uniquely mapped reads (mapping quality >= 10) were considered. We do this across multiple single-cell Strand-seq libraries (n=65) so we observe multiple independent assortments of acrocentric chromosomes in daughter cells. We then compare the Strand-seq strand-state ('ww' - Watson-Watson, 'wc' - Watson-Crick, and 'cc' - Crick-Crick) between tail and head of each pair short and long acrocentric arm across all five acrocentric chromosomes. We assign short and long acrocentric arms based on the best agreement between tail and head strand-states. This means, the strand-state of short and long acrocentric arms that likely belong to each other share the same strand-strand state more often across multiple Strand-seq cells than the other pairs.

### Polishing and patching of T2T-HG002v0.7 to create T2T-HG002v1.1

#### Crowd-sourced curation

In the initial two rounds of polishing and patching, randomly selected examples of different sets of potential corrections (see "Categories of issues/corrections to v0.7" and "Categories of issues/corrections to v0.9") were assigned to volunteer curators using GitHub's issue interface. Using a github repository called "HG002-issues"[108] and python scripts calling github's "hub" API[104], quality issues with the v0.7 or v0.9 assembly were recorded as issues, assigned to curators, discussed in the comments, and eventually closed. Curators were supplied with XML-formatted session files to load into IGV, allowing identical aligned read data and genome annotations to be viewed against the v0.7 or v0.9 assembly in each problem region. In general, if a majority of the issues in a particular category were confirmed by curators to be genuine and correctable, that category's corrections were applied in creating the next T2T-HG002 release.

The scripts used to populate, change the status of, and eventually close issues are publicly available in the software repository for this manuscript (Nancy F. Hansen. (2025). nhansen/github_issues: Release v0.1 software for managing github issues (v0.1)[109].

#### Read alignments

For all rounds of polishing, we aligned newly available sequencing reads using the alignment methods described in McCartney *et al.*, 2022[59]. Briefly, short Illumina, Element Biosciences, and PacBio Onso reads were aligned with BWA-MEM (version 0.7.17-r1188) using default parameters, after which BAM files were sorted with samtools, mate pair coordinates were filled in with samtools fixmates, and PCR duplicates were marked with samtools markdup. Longer ONT and PacBio reads were aligned with Winnowmap2 (version 2.03) using a repetitive k-mer database of 15-mers for down-weighted seeding[110]. The scripts used to run Winnowmap2 and

BWA-MEM are part of the T2T-Polish software package[59] and are included in the manuscript's software archive[104].

### Categories of issues/corrections to v0.7 (round 1)

#### Small corrections

##### Phase-switched and falsely homozygous sites

In an effort to identify locations in the T2T-HG002v0.7 assembly where sequence from one of HG002's parental haplotypes was present on the opposite HG002 haplotype, we ran Merqury on short read (Illumina) data from HG002 and its parents HG003 and HG004, and we performed Strand-seq analysis to phase the assembly and identify heterozygous sites with alleles on the wrong-haplotype. These regions were discovered by first running Merqury[52] on the HG002v0.7 assembly using 21-mer meryl databases of Illumina HiSeq2500 PCR-free 2x250 reads (~350bp insert size) from HG002, HG003, and HG004 (64x for child, 50x for parents) (**Table S1**), with which Merqury creates a database of hapmers and predicts phase blocks (num_switch=100, short_range=20000). Phase-switched Merqury regions were identified as locations where a maternal block is found on a paternal HG002v0.7 chromosome or a paternal block is found on a maternal HG002v0.7 chromosome.

The Strand-seq-based phasing was performed with the v14 parameterization of the PGAS pipeline[85,111]. Briefly, the strand-specific signal of Strand-seq was leveraged to identify phase-informative regions in the assembly. These regions were then phased with StrandPhaseR version #8b93668[112] into sparse whole-chromosome haplotypes. These sparse haplotype blocks then served as the backbone in the final step with WhatsHap v1.1 (Sven Schrinner *et al.* (2020). WhatsHapv1.0[113] that added additional phase information from the long reads to create more dense and reliable haplotype blocks.

We obtained 58 "high confidence" regions by intersecting the Merqury- and Strand-seq-flagged regions detected in the analysis above, and assigned them to volunteers for curation. Curators examined PacBio HiFi, ONT ultra-long, and Illumina reads in IGV, annotated with the locations of haplotype-informative assembly k-mers and heterozygous sites in the assembly. Notably, when long read data were examined by curators, roughly half (25) of the issues were found by curators to be false positives (i.e., the v0.7 assembly consensus was found to be correct), so this set of corrections was not applied, and we pursued an alternative correction discovery method.

Curators had observed that the 23 true positive regions in the abandoned correction set described above nearly always had ONT reads that were long enough to stretch from correctly-phased heterozygous sites of the assembly into regions where the assembly had alleles on the wrong haplotype. Therefore, we used DeepVariant v1.5 to call variants using alignments of R10 ONT reads (**Table S1, Table S2**) to the entire diploid assembly (subsequently referred to as "all-to-all" alignments), filtering the results for homozygous PASS calls with >=95% alternate allele frequency, >=10 reads coverage, and no variant call present on the alternate HG002 haplotype, keeping only the variants that were within regions of homozygosity of at least 5000 bases in the assembly:

```
# call variants withDeepVariant version 1.5.0:
# output hg002v0.7_matpat_r10_simplex_DV_1.5.vcf.gz

# create bed files of all variants in homozygous regions >=5 or 10kb
# with genotypes quality at least 10 and 95% allele fraction:
awk -F"\t" '$3-$2>=5000 {print}'\
hg002v0.7.haplotypemapping.pri.wm.withsimscores.hom.sort.uniq.bed > \
hg002v0.7.haplotypemapping.pri.wm.withsimscores.hom.ge5kb.sort.uniq.bed

bedtools intersect -a\
hg002v0.7.haplotypemapping.pri.wm.withsimscores.hom.ge5kb.sort.uniq.bed -b \
hg002v0.7_matpat_r10_simplex_DV_1.5.allvars.bed -wo > \
hg002v0.7_matpat_r10_simplex_DV_1.5.vars.hom.ge5kb.sort.bed

awk -F"\t" '$12>=10 && $13>=0.95 {print}'\
hg002v0.7_matpat_r10_simplex_DV_1.5.vars.hom.ge5kb.sort.bed >\
hg002v0.7_matpat_r10_simplex_DV_1.5.vars.hom.ge5kb.gq10.af95.bed

# bed files with coordinates of the variants, as well as four column bed files for
lifting:
awk -F"\t" '{OFS="\t"; print $7, $8, $9, "ID"NR, $10, $11, $12, $13, $14}'\
hg002v0.7_matpat_r10_simplex_DV_1.5.vars.hom.ge5kb.gq10.bed >\
hg002v0.7_matpat_r10_simplex_DV_1.5.vars.hom.ge5kb.gq10.varlocs.bed
awk -F"\t" '{OFS="\t"; print $1, $2, $3, $4}'\
hg002v0.7_matpat_r10_simplex_DV_1.5.vars.hom.ge5kb.gq10.varlocs.bed >\
hg002v0.7_matpat_r10_simplex_DV_1.5.vars.hom.ge5kb.gq10.varlocs.forlifting.bed

# lifted locations on opposite haplotypes of hq variants in hom regions:
liftOver \
hg002v0.7_matpat_r10_simplex_DV_1.5.vars.hom.ge5kb.gq10.varlocs.forlifting.bed\
../../chainfiles/v0.7_mat_vs_pat/hg002v0.7.bothdirs.chain \
hg002v0.7_matpat_r10_simplex_DV_1.5.vars.hom.ge5kb.gq10.varlocs.althap.bed\
hg002v0.7_matpat_r10_simplex_DV_1.5.vars.hom.ge5kb.gq10.varlocs.althap.unmapped\
sort -k1,1 -k2,2n -k3,3n \
hg002v0.7_matpat_r10_simplex_DV_1.5.vars.hom.ge5kb.gq10.varlocs.althap.bed >\
hg002v0.7_matpat_r10_simplex_DV_1.5.vars.hom.ge5kb.gq10.varlocs.althap.sort.bed\

# divide alt hap (lifted) variant positions between those that have a variant
# on the alt hap and those that don't -- note that these bed files are in the
# coordinates of the correct alternate location, not the homogenized variant!
bedtools intersect -a \
hg002v0.7_matpat_r10_simplex_DV_1.5.vars.hom.ge5kb.gq10.varlocs.althap.sort.bed -b\
hg002v0.7_matpat_r10_simplex_DV_1.5.allvars.sort.bed -v > \
hg002v0.7_matpat_r10_simplex_DV_1.5.vars.hom.ge5kb.gq10.no_vars_on_alt_hap.bed
bedtools intersect -a \
hg002v0.7_matpat_r10_simplex_DV_1.5.vars.hom.ge5kb.gq10.varlocs.althap.sort.bed -b\
hg002v0.7_matpat_r10_simplex_DV_1.5.allvars.sort.bed >\
hg002v0.7_matpat_r10_simplex_DV_1.5.vars.hom.ge5kb.gq10.vars_on_alt_hap.bed

# join vars with alt coordinates for variants without a variant on the
# alt haplotype:
```

```
awk '{OFS="\t"; print $4, $0}'\
hg002v0.7_matpat_r10_simplex_DV_1.5.vars.hom.ge5kb.gq10.varlocs.bed | sort -k1,1 >\
hg002v0.7_matpat_r10_simplex_DV_1.5.vars.hom.ge5kb.gq10.varlocs.tmp.bed
awk '{OFS="\t"; print $4, $0}'\
hg002v0.7_matpat_r10_simplex_DV_1.5.vars.hom.ge5kb.gq10.no_vars_on_alt_hap.bed |\
sort -k1,1 > tmp.bed
join tmp.bed\
hg002v0.7_matpat_r10_simplex_DV_1.5.vars.hom.ge5kb.gq10.varlocs.tmp.bed |\
awk '{OFS="\t"; print $2, $3, $4, $5, $6, $7, $8, $10, $11, $12, $13, $14}' | sort\
-k1,1 -k2,2n -k3,3n >\
hg002v0.7_matpat_r10_simplex_DV_1.5.vars.hom.ge5kb.gq10.varlocs.no_vars_on_alt_hap.w
ithaltcoords.bed

# use pull_filtered_vcf_lines.pl script to pull appropriate VCF records from
# initial DeepVariant output:
pull_filtered_vcf_lines.pl >\
hg002v0.7_matpat_r10_simplex_DV_1.5.vars.hom.ge5kb.ge10reads.af95.no_vars_on_alt_hap
.vcf
```

This resulted in a list of 1,633 sites with lost heterozygosity in the v0.7 assembly which were correctible using the corresponding DeepVariant call. Reassuringly, this set of corrections included all of the incorrect alleles (true positive corrections) in the Merqury- and Strand-seq-flagged regions which had been considered correct by curators in the previous curation step. This points to long-read discovery of sites with misphased alleles using DeepVariant being a more effective method for correction discovery than Merqury and Strand-seq phasing.

##### Small errors discovered from short-read homozygous calls

To discover small indel errors where the assembly was heterozygous while the sample appeared to be homozygous (i.e., places where a new, erroneous allele had probably been introduced into the assembly on one haplotype only), we examined two sets of DeepVariant v1.5 homozygous non-reference calls. All short reads were aligned with BWA MEM to both paternal and maternal haplotype assemblies independently, causing autosomal reads to “pile up” on either the maternal or the paternal sequence of v0.7. These haplotype-specific assemblies consisted of single haplotypes of the v0.7 assembly combined as well as decoy sequences from the opposite haplotype’s sex chromosome (i.e., chrX added to the v0.7 paternal haplotype, or chrY added to the maternal haplotype), as well as the mitochondrial sequence and the sequence of EBV used to immortalize the HG002 cell line added to both hapotype-specific references. The resulting alignments are subsequently referred to as “all-to-one” alignments.

The first set of short reads aligned consisted of 100x coverage of paired 2x150 small insert reads from Element Biosciences, and the second set of reads consisted of 55x coverage of single-ended 1x100 Onso reads from PacBio (**Table S1**). We first merged the Element and Onso DeepVariant call sets produced by DeepVariant, and then extracted only the homozygous non-reference variant calls (which are equivalent to suggested corrections to the reference haplotype) that appeared in both sets of calls:

```
# do the intersect with bcftools merge (so we can see which were in which)
```

```
bcftools merge SBB_PB/hg002_ssr_DV.vcf.gz \
element_SI/hg002v0.7_mat_element_PCR_free_2x150_100X_DV_1.5.vcf.gz | awk -F"\t" \
'$1~/#/ || ($10~/1\/1/ && $11~/1\/1/) {print}' | bgzip -c  > \
element_SI_and_SBB_PB.mat.merge.hnr.vcf.gz
bcftools merge SBB_PB/hg002v0.7_pat_SSR_DV_1.5.vcf.gz \
element_SI/hg002v0.7_pat_element_PCR_free_2x150_100X_DV_1.5.vcf.gz | awk -F"\t" \
'$1~/#/ || ($10~/1\/1/ && $11~/1\/1/) {print}' | bgzip -c  > \
element_SI_and_SBB_PB.pat.merge.hnr.vcf.gz

# find calls that are in both element and SBB sets:
bedtools intersect -a \
element_SI/hg002v0.7_pat_element_PCR_free_2x150_100X_DV_1.5.hnr.vcf -b \
SBB_PB/hg002v0.7_pat_SSR_DV_1.5.hnr.withheader.vcf > \
element_SI_and_SBB_PB.pat.noheader.vcf
bedtools intersect -a \
element_SI/hg002v0.7_mat_element_PCR_free_2x150_100X_DV_1.5.hnr.vcf -b \
SBB_PB/hg002_ssr_DV.hnr.vcf > element_SI_and_SBB_PB.mat.noheader.vcf
```

This approach yielded 4,203 corrections on the maternal haplotype and 4,104 on the paternal.

##### Small errors discovered from Element heterozygous calls

We next used parental assemblies to phase heterozygous Element calls based on the allele observed in the maternal (HG004) or paternal (HG003) haplotype. To generate a list of proposed corrections from heterozygous DeepVariant calls on Element read alignments to one haplotype at a time ("all-to-one" alignments), we utilized diploid hifiasm (v0.18.9-r527) maternal (HG004) and paternal (HG003) assemblies generated from PacBio HiFi and Oxford Nanopore reads and phased with HiC. These assemblies of HG002's parents were aligned to their corresponding haplotype of HG002v0.7, and variants were then called with dipcall. Heterozygous sites in the parent that were seen to be homozygous in HG002 were used to determine which of the parent's two haplotypes ("hap1" or "hap2") had been inherited by the child. This yielded bed-formatted files against HG002v0.7 showing which of each parent's assembly haplotypes were inherited at any spot in the v0.7 assembly.

In regions where DeepVariant made a heterozygous call against the HG002 assembly, we then selected the allele which was consistent with the inherited haplotype from the respective parental assembly (e.g. HG003 for paternal haplotype), but not observed in the other parent's assembly.

```
# Sample commands for correcting maternal haplotype:
# use HiFi+ONT+HiC HG004 hifiasm assembly to phase element hets on mat that
# are consistent with the inherited HG004 haplotype.
# the regions where hap1 and hap2 from hg4 were inherited are in separate files,
# but they could be combined. They are also separated by whether the element GT
# was 0/1 or 1/2, but the final GT is always 1/1 after selecting the variant
# from hg4 and normalizing. In exploratory work, these
# seemed generally reliable, but likely contain a small number of errors where
# the hg4 assembly was wrong, particularly in homopolymers

# Find regions where each hg4 haplotype was inherited by clustering variants on
# each haplotype. Since the initial simpler merging had a few errors, we used a more
```

```
# stepwise approach: merge variants within 10kb and keep regions >20kb.
# merge remaining regions within 100kb and keep >200kb. merge remaining
# regions within 1Mb and keep >10Mb. Finally merge remaining regions with
# 70Mb
# USES BEDTOOLS v2.30.0, BCFTOOLS v1.17

gunzip -c hg4vhg2mat.pair.vcf.gz | awk '{FS=OFS="\t"} {if($11 ~ /^1\/1/) \
print $1, $2-1,$2+length($4)}' | mergeBed\ -i stdin -d 10000 | awk '$3-$2>20000' |\
mergeBed -i stdin -d 100000 | awk '$3-$2>200000' | mergeBed -i stdin -d 1000000 |\
awk '$3-$2>1000000' | mergeBed -i stdin -d 7000000 >\
hg4vhg2mat.pair.hap1inherited.v2.bed

gunzip -c hg4vhg2mat.pair.vcf.gz | awk '{FS=OFS="\t"} {if($10 ~ /^1\/1/) \
print $1, $2-1,$2+length($4)}' | mergeBed -i stdin -d 10000 | awk '$3-$2>20000' |\
mergeBed -i stdin -d 100000 | awk '$3-$2>200000' | mergeBed -i stdin -d 1000000 |\
awk '$3-$2>1000000' | mergeBed -i stdin -d 7000000 >\
hg4vhg2mat.pair.hap2inherited.v2.bed

subtractBed -a hg4vhg2mat.pair.hap1inherited.v2.bed -b\
hg4vhg2mat.pair.hap2inherited.v2.bed  > hg4vhg2mat.pair.hap1inherited.nohap2.v2.bed
subtractBed -a hg4vhg2mat.pair.hap2inherited.v2.bed -b\
hg4vhg2mat.pair.hap1inherited.v2.bed  > hg4vhg2mat.pair.hap2inherited.nohap1.v2.bed


#Intersect with Element-DV het calls where all data is aligned to maternal
#element vcf from #https://s3-us-west-2.amazonaws.com/human-
pangenomics/T2T/HG002/assemblies/polishing/HG002/v0.7/variants/DeepVariant/all_to_on
e_element_PCRfree/hg002v0.7_mat_element_PCR_free_2x150_100X_DV_1.5.vcf.gz

# also add calls from maternal vs maternal to ignore hets that are already on
# the other haplotype, so likely not errors
dvc get --rev c40d5cf \ https://github.com/ndwarshuis/hg2-t2t-benchmark-
dev.git/pipelines/hg2-trio-dipcall/results/dipcall/hg2_mat_verkko
gunzip -c hifiasm/hg2_mat_verkko/hg2.fixed.vcf.gz |\
awk '{FS=OFS="\t"} {if(!($1 ~ /^#/)) $1=$1"_matERNAL"; print}' | bgzip -c >\
hifiasm/hg2_mat_verkko/hg2.fixedmat.vcf.gz
tabix hifiasm/hg2_mat_verkko/hg2.fixedmat.vcf.gz

# merge vcfs (will have errors in a few places where complex vars
# have differing representations
bcftools merge hg002v0.7_mat_element_PCR_free_2x150_100X_DV_1.5.vcf.gz \
hg4vhg2mat.pair.vcf.gz hg2_mat_verkko/hg2.fixedmat.vcf.gz --force-samples >\
hg4vhg2matvelementDVmatmerge.vcf

# Find element-DV 0/1 calls that match hg4 hap1 where it was inherited,
# and the variant is not in the maternal haplotype
intersectBed -a hifiasm/hg4vhg2matvelementDVmatmerge.vcf -b \
hg4vhg2mat.pair.hap1inherited.nohap2.v2.bed -header | awk '$1 ~ /^#/ \
|| ($10 ~ /^0\/1/ && $11 ~ /^1\/1/  && !($13 ~ /^1/))' |\
bcftools view -s results/dipcall/hg4/hg2.hap1.bam --trim-alt-alleles |\
bcftools norm -f assembly.v0.7.fasta > hg4vhg2mat_hg2elementhet01_hap1inherited.vcf

# Find element-DV 1/2 calls that match hg4 hap1 where it was inherited,
# and the variant in hg4 is not in the maternal haplotype
intersectBed -a hifiasm/hg4vhg2matvelementDVmatmerge.vcf -b\
hg4vhg2mat.pair.hap1inherited.nohap2.v2.bed -header | awk '$1 ~ /^#/ \
|| ($10 ~ /^1\/2/ && (($11 ~ /^1\/1/  && !($13 ~ /^1/)) || \
```

```
($11 ~ /^2\/2/  && !($13 ~ /^2/))))' | bcftools view -s \
results/dipcall/hg4/hg2.hap1.bam --trim-alt-alleles | \
bcftools norm -f assembly.v0.7.fasta > hg4vhg2mat_hg2elementhet12_hap1inherited.vcf

# Find element-DV 0/1 calls that match hg4 hap2 where it was inherited,
# and the variant is not in the maternal haplotype
intersectBed -a hifiasm/hg4vhg2matvelementDVmatmerge.vcf -b\
hg4vhg2mat.pair.hap2inherited.nohap1.v2.bed -header | awk '$1 ~ /^#/ \
|| ($10 ~ /^0\/1/ && $12 ~ /^1\/1/  && !($13 ~ /^1/))' |\
bcftools view -s results/dipcall/hg4/hg2.hap2.bam --trim-alt-alleles | \
bcftools norm -f assembly.v0.7.fasta > hg4vhg2mat_hg2elementhet01_hap2inherited.vcf

# Find element-DV 1/2 calls that match hg4 hap2 where it was inherited,
# and the variant in hg4 is not in the maternal haplotype
intersectBed -a hifiasm/hg4vhg2matvelementDVmatmerge.vcf -b \
hg4vhg2mat.pair.hap2inherited.nohap1.v2.bed -header | awk '$1 ~ /^#/ \
|| ($10 ~ /^1\/2/ && (($12 ~ /^1\/1/  && !($13 ~ /^1/)) || \
($12 ~ /^2\/2/  && !($13 ~ /^2/))))' | \
bcftools view -s results/dipcall/hg4/hg2.hap2.bam --trim-alt-alleles | \
bcftools norm -f assembly.v0.7.fasta > hg4vhg2mat_hg2elementhet12_hap2inherited.vcf
```

This approach yielded a total of 9,061 corrections on the maternal haplotype and 9,819 corrections on the paternal.

#### Corrections to larger regions

##### T2T-Polish-discovered issues

T2T-Polish[59] was used to discover regions of HG002v0.7 with unusually high or low coverage or high amounts of read clipping. First, both PacBio HiFi Sequel reads called with DeepConsensus v1.1 and “R9” Oxford nanopore reads called with guppy remora v6.1.2 (**Table S1**) were aligned separately to the entire diploid v0.7 assembly with winnowmap2[110] (“all-to-all” alignments), after which “issue” bed files were created with T2T-Polish:

```
# create file of repetitive k-mers for winnowmap:
meryl count k=15 hg002v0.7.fasta.gz output merylDB  # add "compress" for homopolymer
compression
meryl print greater-than distinct=0.9998 merylDB > repetitive_k15.txt

# align HiFi and ONT reads with winnowmap2:
winnowmap --MD -W repetitive_k15.txt -ax map-pb -I12g -t$cpus \
hg002v0.7.fasta.gz hifi_dcv1.1.fastq.gz
winnowmap --MD -W repetitive_k15.txt -ax map-ont -I12g -t$cpus \
hg002v0.7.fasta.gz ont_guppy_6_1_2_remora.fastq.gz

# run T2T-Polish scripts to find coverage, quality, and clipping issues:
# generate microsatellite bed files in directory "pattern"
cd pattern
$T2TPolish/pattern/microsatellites.sh hg002v0.7.fasta.gz
# create assembly bed files for "low_support.sh", called by issues.sh:
awk -F"\t" '{OFS="\t"; print $1, 0, $2}' v0.7.fasta.fai > v0.7.bed
cat merqury/v0.7_hybrid/*_only.bed | bedtools merge -i - > v0.7.err.bed
touch v0.7.exclude.bed # no excluded regions
awk -F"\t" '{OFS="\t"; print $1, 0, 10000; print $1, $2-10000, $2}' \
```

```
v0.7.fasta.fai > v0.7.telo.bed
# run issues.sh:
$T2TPolish/coverage/issues.sh hg002v0.7_hifi_dcv1.1.pri.paf \
hg002v0.7_hifi_dcv1.1.pri v0.7 HiFi
$T2TPolish/coverage/issues.sh \
hg002v0.7_guppy_6_1_2_remora.pri.paf hg002v0.7_guppy_6_1_2_remora v0.7 ont
```

The union of these two issue bed files (one for HiFi, one for ONT) contained 344 regions covering roughly 2 megabases of consensus: 35 issues were based on ONT evidence, 337 on HiFi evidence, and 28 were flagged using both platforms.

##### Issues discovered with Flagger/SecPhase

Flagger[10,114] is a read-mapping-based pipeline developed for evaluating phased genome assemblies. It works by taking long reads mapped to the assembly, computing read depth of coverage along the assembly and using Gaussian Mixture Model (GMM) to infer which parts of the assembly are potentially problematic. It can infer three types of misassemblies; erroneous blocks (regions with misjoins or high base-level error rate), false duplications and collapsed blocks. It also incorporates mapping quality of read alignments as an additional signal to detect false duplications.

Flagger v0.2 was run on the same read alignments used for the T2T-Polish analyses (see "T2T-Polish-discovered issues"), after which regions which were flagged for both the HiFi alignments and the ONT alignments were gathered using bedtools intersect, resulting in 159 assembly regions (called "Flagger Intersect Regions" in the issues github repository), 34 of which overlapped with at least one of the 344 (union of HiFi and ONT) T2T-Polish-called regions (these Flagger calls are available on AWS at https://s3-us-west-2.amazonaws.com/human-pangenomics/index.html?prefix=T2T/HG002/assemblies/qc/flagger/).

##### Patches to problematic regions using a newer assembly (HG002v0.8)

All patch corrections to HG002v0.7's larger problematic regions used sequence from a later assembly that had been run with newer read datasets (**Table S1**) in May, 2023 using verkko version 1.3.1 (commit 0268963fb2eda147e9dfb8d4b0291fa06d47b4bb). This assembly was labeled HG002v0.8, and its scaffolds, labeled with human chromosome assignments, are available on AWS (source data and assembly URL in **Table S1**).

In preparation for applying HG002v0.8 patches to HG002v0.7, a set of 137 "problem regions" were compiled (see below). Of these, regions which could be successfully lifted from HG002v0.7 to HG002v0.8 in one piece after adding 2000 bases of buffer sequence to both the 5' and 3' end were curated by viewing their read alignments in IGV to assess whether the HG002v0.8 consensus represented a significant improvement over the HG002v0.7 consensus.

These 137 regions were composed of the following sets: First, 18 non-acrocentric problem regions that were both called by "Flagger" as collapses, duplications, or errors and marked as issues by the T2T-Polish pipeline. So-called "Intersect" regions were then determined by taking the intersections of regions tagged by the program Flagger as "Col", "Dup", or "Err" and regions also marked as an issue by the T2T-Polish pipeline when run on both HiFi and ONT primary alignments of reads to the entire diploid v0.7 assembly. Second, an additional twelve regions that contained a high density of DeepVariant homozygous non-reference calls on (a) three different long read alignment sets (HiFi/DeepConsensusv1.1, Duplex ONT, and R10 simplex ONT) ("all-to-all" alignments) or (b) alignments of high-accuracy short reads, both Element and Onso, to single haplotypes of the v0.7 assembly ("all-to-one" alignments) were included to potential patching. The final set of potential patch regions were those that had an unusual amount of sequence difference between v0.7 and v0.8. These were 10kb windows of v0.7 which had uniquely liftable regions to v0.8, but which also had at least 200 bases of difference between the two assemblies within the liftover chain. There were 122 of these intervals, 15 of which overlapped with one of the 30 regions in the first set described above. After curation, 74 of the large regions found with these methods were correctable using patches from the HG002v0.8 assembly.

#### Final assembly corrections applied to v0.7

The corrections described above fell into the following categories:

- "False Hets": this category included short read homozygous calls and long read homozygous calls (7,849 corrections)
- "False Homs": this category included parent assembly-phased short read heterozygous calls and phase-switched and falsely homozygous long read heterozygous calls (11,726 corrections)
- "Het sites with errors": this category included short read heterozygous calls where one allele disagreed with HG002v0.7 (7,965 corrections)
- "Large structural errors": this category includes the large regions identified by T2T-Polish and flagger and were improved in HG002 v0.8 (74 corrections, each replacing an average of 5,081 bp and adding or subtracting, on average, 2,080 bp)

Of the 27,540 small corrections, 97 fell within the boundaries of the 74 larger patch corrections described above and were therefore not included. This resulted in a set of 27,517 small and large corrections which were applied to the HG002v0.7 assembly using bcftools:

```
cat ../../assemblies/v0.7.fasta | bcftools consensus -c \
hg002v0.7_to_hg002v0.9.chain corrections_large_and_small.withheader.vcf.gz \
> v0.9.fasta
```

The distribution of sizes for this correction set shows that the vast majority of corrections were one-base deletions from HG002v0.7 (**Figure S2**). The resulting corrected assembly, named T2T-HG002v0.9, was released in July, 2023.

### Categories of issues/corrections to v0.9 (round 2)

#### Corrections to problem areas in “difficult” regions of T2T-HG002v0.9

##### Correction of errors identified in centromeres

Errors in the centromeric regions within T2T-HG002v0.9 assembly were identified by running NucFreq (v0.1) with the following command:

```
	NucPlot.py {input.bam} {output.png} --regions {centromeric_region_coordinates} -r
{repeatmasker.out}
```

NucFreq plots were visually inspected to identify potential collapses in sequence, misjoins, or other types of assembly errors by looking for deviations in the first and second most common base relative to the mean. Seven centromeres were identified as having potential misassemblies: chr4_MATERNAL, chr4_PATERNAL, chr5_PATERNAL, chr6_MATERNAL, chr6_PATERNAL, chr13_PATERNAL, and chr17_MATERNAL. To patch the assembly errors in these centromeres, we generated a hifiasm v0.19.5 assembly using the following command:

```
hifiasm -o {output_dir_prefix} -t{num_of_threads} --ul $(cat {ont_fastq.fofn} | \
sed -z 's/\n/,/g;s/,$/\n/') -1 {mat.yak} -2 {pat.yak} \
$(cat {pacbio_hifi_fastq.fofn})
```

We aligned HG002v0.9 centromeric contigs to the HG002 hifiasm assembly using minimap2 v2.28 using the following parameters: -I 15G -a --eqx -x asm20 -s 5000 and extracted the corresponding centromeric regions that spanned the assembly errors in both the HG002v0.9 assembly and hifiasm assembly using seqtk (v1.4). Then, we ran RepeatMasker (v4.1.0) on each of the fasta files using the following parameters: {-species human -dir {output_dir} -pa {num_of_threads}. We also ran StringDecomposer (v1.0.0) on each of the fasta files using α-satellite monomer inputs specific to each chromosome and derived from the T2T-CHM13 v2.0 genome. Using the StringDecomposer outputs, we manually patched the HG002v0.9 genome assembly with the hifiasm assembly. We validated the successful repair of these regions by aligning HG002 PacBio HiFi data to the HG002v0.9 assembly containing the repaired centromeric contigs with the following command:

```
pbmm2 align --log-level DEBUG --preset SUBREAD --min-length 5000 -j 8.
```

We then ran NucFreq on each centromeric region using the same command as above and assessed whether the centromeric regions had first and second most common base frequencies similar to the mean of the region. This confirmed that the following centromeres were successfully repaired: chr4_MATERNAL, chr4_PATERNAL, chr5_PATERNAL, chr13_PATERNAL, and chr17_MATERNAL. Two centromeres still had assembly errors (chr6_MATERNAL, chr6_PATERNAL) that were unable to be repaired.

The 19 regions corrected on seven HG002v0.9 centromeres had the following v0.9 coordinates: chr4_MATERNAL:49232779-49280263, chr4_MATERNAL:51637922-51647612, chr4_PATERNAL:52166546-52240961, chr4_PATERNAL:52712713-52997607, chr5_PATERNAL:46790217-46791543, chr5_PATERNAL:47576796-47581721, chr5_PATERNAL:49782320-49787412, chr5_PATERNAL:51147396-51149093, chr5_PATERNAL:52219891-52225667, chr6_MATERNAL:62162402-62167836, chr6_PATERNAL:59153776-59162947, chr6_PATERNAL:59355877-59361991, chr6_PATERNAL:61597046-61614878, chr6_PATERNAL:62187141-62191384, chr6_PATERNAL:62281724-62324340, chr6_PATERNAL:62686192-62728299, chr6_PATERNAL:62762429-62774151, chr13_PATERNAL:11176318-11202477, chr17_MATERNAL:25766051-25799993

##### Correction of errors identified in telomeres

Reads from Oxford Nanopore Technologies (ONT) telomere enrichment sequencing (Telo-seq) were generated using R10 chemistry with v4.3 SUP basecalling. Since the telomere analysis was begun before the release of v0.9, this analysis was completed using the v0.7 assembly and then carried over to determine corrections to v0.9. Reads were initially mapped to the HG002v0.7 phased reference genome using minimap2 (v2.26), and single nucleotide polymorphisms (SNPs) and insertion-deletion variants (indels) were called using FreeBayes (v1.3.7). Variants were filtered to retain only homozygous calls using bcftools (v1.17), and a consensus sequence was generated incorporating these variants into the reference sequence. The ONT telo-seq reads were subsequently re-mapped to this consensus sequence, and final corrections were made manually by curation using the Integrative Genomics Viewer (IGV v2.16.2) and Geneious Prime (v2023.1).

Once this analysis was completed, the corrected telomere consensus sequences were compared to the newly-released HG002v0.9 by first extracting 100,000 base pairs from each end of each chromosome into a FASTA formatted file, then aligning the curated telomere sequences to them with minimap2 using the parameters:

```
minimap2 -x asm5 -g 50000 -n 0 -f 0 teloseqs.fasta v0.9.telomere.fasta >>
p_teloarms_ONTv14_to_v0.9.minimap2.out
```

After filtering the resulting alignments to require the tag “dv:f:0.00”, a shell script was used to generate a VCF patching the correct coordinates into the v0.9 assembly see the software repository for this manuscript for the exact commands performed to do the patching[104].

##### Extension of v0.9 rDNA flanking sequences into the rDNA gap

With the exception of the rDNA array on the paternal copy of Chr13 (whose length was corrected using ultra-long ONT reads, see the section “Lengthening of paternal Chr13’s rDNA array using spanning ONT reads”), the rDNA arrays of v0.7 contained Ns approximating the length of each array flanked by low-accuracy consensus. Generation of new, more accurate flanking sequence for the rDNA arrays was done with the script “improve_gaps_ont.py”[104], which iteratively extends unique region into the tangles when there is a clear majority (best > 1.5* second_best) of ONT reads which are anchored on the unique region and support the extension.

These new contigs were then aligned to the T2T-HG002v0.9 assembly with minimap2 and the alignments were used to choose the best matches between extended contigs and flanking sequences.

From the minimap2 results, high identity mappings were extracted:

```
# find contig to chromosome mapping from minimap2 output:
grep 'dv:f:0.000' patchv2_to_0.9.out | \
awk -F"\t" '{OFS="\t"; print $6, $1}' | sort | uniq | \
sed 's/_0/_P/' | sed 's/_1/_Q/' | awk '$1!="chr13_PATERNAL" {print}' \
> patch_v2_contig_map.txt
```

A fasta-formatted file of flanking sequences surrounding the rDNA arrays was created:

```
export BED=$1
export ASSEMBLYFASTA=/data/Phillippy/projects/HG002_diploid/assemblies/v0.9.fasta
export NACROS=`cat $BED | wc -l`
export FLANKFILE=`echo $BED.flank.fasta | sed 's/.bed//'`
echo $NACROS" lines in "$BED
rm -f $FLANKFILE
for i in `seq 1 $NACROS`; do
        export CHROM=`head -$i $BED | tail -1 | awk -F"\t" '{print $1}'`
        export START=`head -$i $BED | tail -1 | awk -F"\t" '{print $2}'`
        export END=`head -$i $BED | tail -1 | awk -F"\t" '{print $3+1}'`
        export ENDP=`echo $END + 2000000 | bc`
        echo "Found " $CHROM 1 $START $END $ENDP
        (echo ">"$CHROM"_P"; samtools faidx $ASSEMBLYFASTA $CHROM:1-$START | tail -n
+2) >> $FLANKFILE
        (echo ">"$CHROM"_Q"; samtools faidx $ASSEMBLYFASTA $CHROM:$END-$ENDP | tail -
n +2) >> $FLANKFILE
done
samtools faidx $FLANKFILE
```

Then each high-identity rDNA-matching contig sequence was aligned to its corresponding v0.9 flanking sequence one contig at a time:

```
export FULLASSEMBLY=v0.9.fasta
export ASSEMBLY=rDNA_N_coords.v0.9.flank.fasta
export PATCHSEQS=patch_v2/rdna_patch_v2.fasta
# map has v0.9 chrom with _[PQ], then contig from Dima's file
export PATCHMAP=patch_v2_contig_map.txt
export POUTPUT=pside_patch_to_v0.9.minimap2.out
export QOUTPUT=qside_patch_to_v0.9.minimap2.out
rm -f $POUTPUT $QOUTPUT
for contig in `awk '{print $1}' $PATCHSEQS.fai`; do
    export CONTIG=$contig
    export CHROMEND=`awk '$2==ENVIRON["CONTIG"] {print $1}' $PATCHMAP`
    export CHROM=`echo $CHROMEND | sed 's/ATERNAL_P/ATERNAL/' | sed
's/ATERNAL_Q/ATERNAL/'`
    export END=`echo $CHROMEND | sed 's/.*_//'`
    export CHROMLEN=`awk -F"\t" '$1==ENVIRON["CHROM"] {print $2}' $FULLASSEMBLY.fai`
    export LEN=`awk -F"\t" '$1==ENVIRON["CHROMEND"] {print $2}' $ASSEMBLY.fai`
    #export START=`echo $LEN-100000 | bc`;
    echo "CONTIG: "$CONTIG " CHROMEND: "$CHROMEND "END: "$END " CHROM: "$CHROM "
LENGTH: "$CHROMLEN "CHROMENDLEN: "$LEN
```

```
        samtools faidx $ASSEMBLY $CHROMEND > assembly.fasta
        samtools faidx $PATCHSEQS $CONTIG > patchseq.fasta
        if [[ $END == "P" ]]; then
           export OUTPUT=$POUTPUT
        elif [[ $END == "Q" ]]; then
           export OUTPUT=$QOUTPUT
        fi
        minimap2 -x asm5 -g 50000 -n 0 -f 0 patchseq.fasta assembly.fasta | \
           grep 'dv:f:0.000' >> $OUTPUT
    done
```

The resulting minimap2 PAF records were used to extract patch coordinates to input to the script “create_patch_vcf.py”[104], which creates a correction VCF file which was included in the final creation of the v1.0 assembly:

```
export V9FASTA=v0.9.fasta
export RDNAFASTA=rdna_patch_v2.fasta
export PPATCHCOORDS=pside_patch_to_v0.9.minimap2.out
export QPATCHCOORDS=qside_patch_to_v0.9.minimap2.out
export V9NCOORDS=rDNA_N_coords.v0.9.bed
export COORDFILE=v0.9.rdna_patch_v2.coords.txt
rm -f $COORDFILE
export VCFFILE=v0.9.rdna_patch_v2.vcf
rm -f $VCFFILE
for contig in `awk -F"\t" '{print $6}' $PPATCHCOORDS`; do
    export CONTIG=$contig
    export CHROMSIDE=`awk -F"\t" '$6==ENVIRON["CONTIG"] {print $1}' $PPATCHCOORDS`;
    export CHROM=`echo $CHROMSIDE | sed 's/L_P/L/' | sed 's/L_Q/L/'`
    export V9START=`awk -F"\t" '$6==ENVIRON["CONTIG"] {print $3+1}' $PPATCHCOORDS`;
    export PATCHCOORDS=`awk '$6==ENVIRON["CONTIG"] {print $8+1" "$7}' $PPATCHCOORDS`;
    export PATCHSTART=`echo $PATCHCOORDS | sed 's/ .*//'`
    export PATCHEND=`echo $PATCHCOORDS | sed 's/.* //'`
    export V9END=`echo "$V9START + $PATCHEND - $PATCHSTART" | bc`
    echo -e "$CHROM\t$V9START\t$V9END\t$CONTIG\t$PATCHSTART\t$PATCHEND\t+" >> $COORDFILE
done

for contig in `awk -F"\t" '{print $6}' $QPATCHCOORDS`; do
    export CONTIG=$contig
    export CHROMSIDE=`awk -F"\t" '$6==ENVIRON["CONTIG"] {print $1}' $QPATCHCOORDS`;
    export CHROM=`echo $CHROMSIDE | sed 's/L_P/L/' | sed 's/L_Q/L/'`
    export V9QSTART=`awk -F"\t" '$1==ENVIRON["CHROM"] {print $3+1}' $V9NCOORDS`
    export V9FLANKEND=`awk -F"\t" '$6==ENVIRON["CONTIG"] {print $4}' $QPATCHCOORDS`;
    export PATCHCOORDS=`awk '$6==ENVIRON["CONTIG"] {print $8+1" "$7}' $QPATCHCOORDS`;
    export PATCHSTART=`echo $PATCHCOORDS | sed 's/ .*//'`
    export PATCHEND=`echo $PATCHCOORDS | sed 's/.* //'`
    export V9PATCHEND=`echo "$V9FLANKEND + $V9QSTART - 1" | bc`
    export V9PATCHSTART=`echo "$V9PATCHEND - $PATCHEND + $PATCHSTART" | bc`
    echo -e "$CHROM\t$V9PATCHSTART\t$V9PATCHEND\t$CONTIG\t$PATCHSTART\t$PATCHEND\t-" >>
$COORDFILE
done

# extract patch VCF:
python3 create_patch_vcf.py -c $COORDFILE -r $V9FASTA -p $RDNAFASTA --vcf $VCFFILE
```

#### Lengthening of paternal Chr13's rDNA array using spanning ONT reads

We discovered that two reads from an ultra-long Dorado-called ONT dataset (read names 03fb2d40-804f-418c-9195-872db2ea628e and 0bba627b-004a-42b0-8ff9-a76f9ca9ec6f) spanned the chr13_PATERNAL rDNA array when aligned to the v0.9 assembly. The paternal Chr13 rDNA array had been assembled with two copies of the rDNA unit in the v0.7 assembly, and had remained that way in the v0.9 assembly. The new spanning reads were aligned with clipping by minimap2 because they contained more copies of the rDNA repeat unit than the v0.9 reference. Comparison of the two spanning reads to each other as well as to the assembly with ModDotPlot[115] showed that the ONT reads actually had six copies of the repeat unit between the non-rDNA ends, i.e., four more units than the v0.9 assembly has (**Figure S11**).

We found that a verkko assembly done with ONT duplex reads[116] had assembled the entire chr13_PATERNAL rDNA array with the same number of copies as the spanning ultralong ONT reads, so we extracted that section of the duplex assembly with 500 kb flanks and used it as a patch to the v0.9 assembly, determining the coordinates by matching the flanking regions of v0.9 with the flanks of the duplex assembly using minimap2.

### Correction of small errors using trio-based Onso, standard and long-insert Element, Illumina, and HiFi DeepVariant calls

We again utilized DeepVariant calls (**Table S2**) to apply trio-based approaches to phase small corrections before applying them to v0.9. Three variant sets were filtered and included in the corrections made to v0.9 to obtain v1.0.1:

1. Taking advantage of the availability of new high-accuracy, higher depth-of-coverage short reads from the entire family trio, DeepTrio was run on Onso and Element reads from HG002, HG003, and HG004 aligned separately to either the maternal+Y of the paternal+X v0.9 references (**Table S1, Table S2).** High-confidence HG002 variants which appeared in both the Onso and the Element call sets were filtered for Mendelian consistency and to remove calls in the VDJ regions or in regions with high depth of coverage, as well as indel calls of 10 or more bases, which were observed to be potential somatic variants. This yielded 1,306 corrections.
2. To rescue short read-based calls that were filtered out of the first group, we aligned Onso, standard and long-insert Element, and Illumina reads from HG002, HG003, and HG004 to the separate maternal+Y and paternal+X v0.9 fasta files ("all-to-one" alignments) (**Table S1**). Then we called variants using DeepTrio (**Table S2**), merged and genotyped the gvcfs using GLnexus, and phased the HG002 variants from each technology using rtg-tools-3.12.1/rtg mendelian with the --Xphase --all-records options. Variant alleles phased to maternal or paternal haplotype with GQ and DP greater than 20 in all 3 individuals and DP less than double the median coverage in HG002 were identified as potential errors. We removed any variants in IGH, IGL, or IGK regions because they rearrange somatically. We also called variants with DeepVariant from HiFi reads aligned to the combined maternal and paternal haplotypes ("all-to-all" alignments). Based on manual curation, we kept phased short read variants if they had a homozygous or heterozygous HiFi call with the same allele, or if they were 1 bp indels in

homopolymers where HiFi is less reliable. Example commands used to do this are available in the software repository for this manuscript[104]. This yielded 2,998 corrections.
3. Finally, 1,505 homozygous variant calls in the HiFi "all to all" DeepVariant set above (**Table S2)** were filtered to require GQ>=20 and DP<75 and included as additional corrections.

#### Final assembly corrections applied to v0.9

When the three categories of small corrections were combined and filtered to remove variants that lie within the larger patched regions, 4,482 small variants remained (displayed in blue in the correction size histogram in **Figure S2**).

Once v1.0 was created, it was discovered that we had inadvertently introduced ambiguity codes for three base pairs into the assembly due to heterozygous sites in the VCF file that was used as input to "bcftools consensus". These three sites were at the following (v1.0) positions:

chr1_PATERNAL:73256576 K
chr3_PATERNAL:42065937 Y
chr8_MATERNAL:126720079 M

The decision was made to correct each of these three sites to what looked to be the most likely ATGC nucleotide. This resulted in the chr1_PATERNAL K and the chr3_PATERNAL bases being replaced with T's, and the chr8_MATERNAL base being replaced with an A. The resulting assembly after these three corrections were applied was designated T2T-HG002v1.0.1 and was released and submitted to Genbank in October, 2023.

### Categories of issues and corrections to v1.0.1 (round 3)

The polishing and patching of T2T-HG002v1.0.1 to create v1.1 ("round 3") attempted to address issues that were raised by several QC tools. Specifically, we applied filtered variant call set corrections created by DeepVariant[57] and DeepPolisher[48] and also created and applied assembly patches, where possible, to the following problem regions: 1) issues submitted to the github "HG002-issues" repository[108], 2) so-called "NIST error exclusion regions" (see the section "DeepVariant calls and NIST exclusion regions", below), 3) regions with large numbers of DeepPolisher corrections which hadn't survived filtering, 4) regions surrounding unresolved Sniffles[58] suspect regions discovered by a new version of Flagger, v0.4.0 (see "Results of re-running Flagger on the T2T-HG002v1.0.1 assembly").

#### Patching v1.0.1 with consensus from two alternative assemblies

Because there were suspected erroneous regions of v1.0.1 for which we were unsure of the correct HG002 consensus sequence, we attempted to use two newer alternative assemblies' consensus to repair suspect v1.0.1 regions. Specifically, in cases where the two new assemblies agreed with each other but disagreed with the v1.0.1 assembly, we constructed a patch to correct v1.0.1 with the newer sequence. To do this, we used two python scripts called "retrieve_patchseqs_from_bam.py" and "create_patch_vcf.py"[104], which parse minimap2 alignments of two alternative assemblies to the unpolished reference to develop a patch for a particular region of the unpolished assembly. If the two newer assemblies have identical consensus sequences to each other, and both have a different consensus from the unpolished assembly, retrieve_patchseqs_from_bam.py creates a VCF-formatted correction using the alternative assemblies' shared sequence as the new allele. We refer to this process as "assembly patching".

The first assembly we used for assembly patching was a recent assembly run with the Verkko2 assembler and phased with trio data (**Table S12**). The second assembly was the "lc24 medaka-polished" assembly of HG002 released by Oxford Nanopore at the London Calling 2024 conference (**Table S12**). Both of these assemblies successfully spanned about two thirds of the suspect or problem regions we evaluated, and both confirmed the v1.0.1 assembly consensus in about one third of those cases. In another third, the two assemblies suggested a common correction for v1.0.1 and a patch was applied. The final third of regions spanned by both assemblies revealed differing sequences between the two assemblies, so no correction was made in those cases.

#### Polishing and patching v1.0.1 to create v1.1

In polishing v1.0.1, DeepPolisher was run to produce VCF files with suggested corrections that were then filtered and applied directly to the v1.0.1 reference. The DeepPolisher calls were obtained by running DeepPolisher as described in the section "DeepPolisher calls on HG002v1.0.1", below, and randomly selected DeepPolisher calls were manually curated along with the proposed corrections in other categories. Because curation of these calls revealed a large number of false positives among "DeepPolisher only" calls, only 840 (out of 2,066 total) DeepPolisher calls which agreed with corrections in the DeepVariant call set were applied. The rest of the DeepPolisher calls were clustered into regions which were later considered for potential patching (see below).

As we had done in the previous two rounds, we also ran DeepVariant on multiple sets of reads aligned to the new v1.0.1 assembly, including read sets that had been used in previous rounds, as well as newly available "Q28" ONT and ONT R10 duplex reads (**Table S2)**. The resulting calls were assessed for reliability and accuracy, then used to cross-reference with the DeepPolisher calls as described above, and also used to delineate "exclusion regions" where errors in the v1.0.1 assembly were suspected (see "DeepVariant calls and NIST exclusion regions").

To ensure that the corrections suggested by DeepPolisher, DeepVariant and our patching process were of high quality, we performed manual curation of randomly selected examples from various sets of corrections that overlapped with each other or were distinct, determining whether subsets of our algorithmic corrections could be confirmed as reliable or not by manual curation. Based on the results of this curation, all corrections were applied from the DeepVariant Element-based calls and all successful assembly patches derived from NIST excluded regions (see the section "DeepVariant calls and NIST exclusion regions"). In addition, any correction seen in two or more of the curated correction sets was applied. Corrections suggested only by DeepPolisher had been found to be unreliable in a majority of the ten examples we curated, but in an attempt to “rescue” possibly valid corrections in the DeepPolisher-only set, we attempted to use the patching method described above to correct the DeepPolisher-only regions, adding a 100 base pair buffer to each side of call locations and running the retrieve_patchseqs_from_bam.py script[104]. Of 1,226 DeepPolisher-only corrections, 346 (179 paternal) had a shared alternative consensus in the Verkko2/trio and medaka assemblies to the HG002v1.0.1 assembly, and so a patch was applied, but in another 386 (174 paternal) the two assemblies both had consensus that agreed with the v1.0.1 sequence so no patch was applied. Another 426 (195 paternal) DeepPolisher problem regions had two differing consensus sequences in the two assemblies, so no patch was applied, and finally 68 (25 paternal) regions could not be spanned by both assemblies, making patching impossible.

#### Methods used to discover regions for attempted patching

*DeepPolisher calls on HG002v1.0.1*
40× PacBio HiFi DCv1.2 reads for HG002 were obtained from the HPRC, and aligned to the HG002v1.0.1 diploid assembly using minimap2 with parameters -a -x map-hifi --cs --eqx -L -Y -I8g. In order to correct read phasing in long stretches of homozygosity in these alignments, the PHARAOH pipeline[48] was run with default parameters, using as input 40× ONT UL >100kb HG002 reads from the HPRC aligned to each haplotype of the HG002v1.0.1 assembly with minimap2 and parameters `-a -x map-ont --cs --eqx -L -Y`. DeepPolisher was run with docker version google/deepconsensus:polisher_v0.0.8_12122023 and model checkpoint 665. Polishing edits were filtered according to the following filters: GQ > 20 for 1bp insertions, GQ > 12 for 1bp deletions, and GQ > 5 for all other edit sizes, as recommended in Mastoras et al.[48].

*Github “Issue” submissions*
Issues submitted to the HG002-issues github repository[108], with the exception of those in the rDNA arrays and gaps, were also evaluated for potential patching. Of 29 (13 paternal) regions flagged as issues on the github site, the patching process was able to correct 15 (7 paternal) while 10 (4 paternal) had disagreeing consensus in the two assemblies, 3 (1 paternal) remained unspanned by one or both assemblies, and one paternal region had complete agreement between the patch assemblies and v1.0.1.

*Re-run of Flagger on the T2T-HG002v1.0.1 assembly*
We used the Flagger-v0.4.0 pipeline that includes a new module for detecting annotations with coverage biases[114]. We were motivated by the fact that some human satellites have exhibited read coverage biases which depend on both the satellite family and the sequencing platform. For example, ONT R10.4.1 Ultra-Long reads have been shown to have upward coverage bias in HSat-1A. The pipeline takes a list of satellite annotations in BED format, compares the median coverage for each annotation to the genome-wide coverage and determines whether it exhibits a coverage bias. This allows Flagger to fit the GMM with an independent set of parameters for each biased annotation, improving the accuracy of misassembly detection in those regions.

To assess assembly quality, we mapped long reads from three sequencing platforms, HiFi Revio (3 flow cells, ~100× coverage), ONT R10 Duplex (~80×), and ONT R10 Ultra-Long (~120×), to the v1.0.1 assembly using Winnowmap (**Table S1)**. We also provided the censat and segdup annotation to the Flagger pipeline for detecting arrays with biased coverage and also stratifying final results. All the bed files used to run Flagger are available in an archived version of the Flagger github repository[114]. The WDL for running Flagger-0.4.0 and instructions for running it are available in the software archive for this manuscript[104].

*DeepVariant calls and NIST exclusion regions*

The proposed DeepTrio corrections to T2T-HG002v1.0.1 were similar to those used in polishing v0.9, using the same standard insert Element and HiFi reads from the trio aligned separately to each haplotype of HG002v1.0.1 (“all-to-one” alignments), as well as HiFi, ultralong ONT, and duplex ONT aligned to combined haplotypes (“all-to-all” alignments) (**Table S2**). Several categories of variants were used to generate the “NIST exclusion regions” by adding 50bp to each side of each of the following categories of variants and merging regions within 1000 bp:

1. Element DeepTrio calls phased to a haplotype and in a homopolymer or dinucleotide tandem repeat
2. Element DeepTrio calls phased to a haplotype and not in a homopolymer or dinucleotide tandem repeat but supported by any of the long read callsets
3. Element DeepTrio calls not phased to a haplotype but not in the opposite haplotype and supported by any of the long read callsets
4. HiFi DeepTrio calls phased to a haplotype and not in a homopolymer or dinucleotide tandem repeat
5. Heterozygous or homozygous DeepVariant calls in HiFi and either ONT dataset and <8 bp in size (because larger heterozygous variants tended to be winnowmap2 alignment issues)
6. Homozygous DeepVariant calls in HiFi and either ONT dataset
7. Heterozygous or homozygous DeepVariant calls in both ONT datasets, not in homopolymers, and <8 bp in size (because larger heterozygous variants tended to be winnowmap alignment issues)

Separate from the exclusions bed file containing these regions, we also attempted to identify *de novo* and mosaic variants and created a separate bed file for these because both are mostly mutations which arose in the cell line and are more likely to vary between cell line batches and not in the iPSC HG002 line. We used TNscope with 300x Illumina WGS from the NIST reference material batches of HG002, HG003, and HG004 aligned separately to each HG002 haplotype. We treated the combined parents as “normal” and HG002 as “tumor”. After curating both small and larger variants, they generally seemed to be true de novos/mosaics if HG002 DP>150 and HG003 and HG004 VAF=0, though many are low VAF (~3-5%) in HG002. We separated those that match the other haplotype and lifted them over to the other haplotype since they are either *de novo* or mosaic on the other haplotype. The remaining are likely mosaic but not clear as to which haplotype so we excluded them on both haplotypes. We also used HiFi Revio reads from the trio aligned separately to each haplotype with DeepSomatic in a similar way. Upon curation, these tended to be most likely to be true mosaic or *de novo* variants if they were also in

DeepVariant calls from long reads aligned to both haplotypes and not in parental DeepVariant calls or in homopolymers. 50bp were added to each side of these potential mosaic and *de novo* variants, and regions within 1000bp were merged to get 2,686 regions across both haplotypes.

An example of a simplified process for this which might be more generally applicable to polishing other assemblies with Element or Onso reads from a trio, where the only filtering was to select 1 and 2 bp indels in homopolymers or dinucleotide tandem repeats with HG002's GQ>20 and DP>20 and DP<double the mean coverage (100 in this case), is presented in the software archive for this manuscript[104].

#### Total corrections made in three rounds of polishing

VCF files containing all corrections made to HG002v0.7, HG002v0.9, and HG002v1.0.1 (i.e., all of the corrections applied to HG002v0.7 in the three rounds of polishing resulting in HG002v1.1) are available as VCF-formatted files on the HG002-issues github site[108]. These files were used to calculate the statistics reported in Table **S3** regarding the types of corrections applied in the three rounds of polishing, and correction lengths for the small corrections are displayed in **Figure S2**.

### Evaluation of T2T-HG002v1.1

#### Strand-seq assembly evaluation

We used the Strand-seq data generated from HG002[24] (**Table S1**) to evaluate the directional and structural contiguity of the HG002v1.1 assembly. Strand-seq data for HG002 are available via the Genome in a Bottle[2] and the human pangenomic AWS portal. First, selected high quality Strand-seq libraries (n=131) were aligned to the HG002v1.1 assembly using BWA-MEM[117] (v0.7.17-r1188), after which reads were sorted by samtools[118] (v1.15.1) and duplicate reads were marked by sambamba[119] (v1.19). To detect putative misassembly breakpoints in T2T-HG002v1.1 we concatenated read alignments across all 131 Strand-seq libraries to create a high coverage Strand-seq read profile across the whole genome[120]. We ran breakpointR[121] on such high coverage Strand-seq data to detect any recurrent strand-state changes that are indicative of genome misassemblies using runBreakpointr function (set parameters: pairedEndReads = FALSE, windowsize = 50000, binMethod = "size", genoT = 'binom', background = 0.1, peakTh = 0.25, minReads = 50). Specifically we were looking for regions where all Strand-seq reads are genotyped as homozygous inverted ('ww'; 'HOM' - all reads mapped in minus orientation) or heterozygous inverted ('wc', 'HET' - approximately equal number of reads mapped in minus and plus orientation). Homozygous inverted regions are indicative of assembly misorientation while heterozygous inverted regions are mostly marking normal heterozygous inversions in a given individual. In rare cases heterozygous inverted regions might represent assembly chimerism if present at the ends of the assembled contigs (chromosomes).

**Figure S3** shows an ideogram in which regions are genotyped as either in complete agreement with the assembly (reference, ‘cc’) or as heterozygous (‘wc’). Heterozygous regions are usually caused by low mappability regions such as short acrocentric arms, centromeres, and repetitive regions. A subset of these mark positions of heterozygous inversions highlighted by arrowheads. We did not observe any region genotyped as homozygous inverted and thus we concluded there is no Strand-seq evidence of  misorientation present in these assemblies.

#### k-mer based evaluation

The final round of evaluation was made using reads from the latest sequencing platforms available. Prior rounds of curation and evaluation had shown PacBio Revio with SPRQ and Element AVITI UltraQ to have the most consistent bases for homopolymers and microsatellite repeats compared to PacBio Revio HiFi (without SPRQ) or Illumina. We used GenomeScope2 and overall genome coverage to evaluate sequencing platforms in an effort to avoid sequencing biases, in particular to find a platform that complements the GA-microsatellite dropouts in PacBio long-reads with minimum sequencing errors.

Using meryl v1.4.1, k=31 mers were collected from the following read sets for evaluation:

- PacBio SPRQ on Revio, 2 cells sequenced with 30 hours (100x)
- PacBio HiFi on Revio, DeepConsensus v1.2, 2 cells sequenced with 24 hours (70x)
- Element AVITI UltraQ Standard 400bp insert, 2x150 (80x)
- Illumina HiSeq2500 2x250, GIAB (60x)
- PacBio Onso 2x150 sequenced on Revio (45x)

Next, we used `GenomeScope2` (commit fdeb89178d506c9af2c5d0d103e0135a164889a3) to evaluate the error rate within each set of reads.

```
$tools/genomescope2.0/genomescope.R -i $pf.k31.hist -k 31 -o $pf.gs2 --fitted_hist
```

For PacBio SPRQ reads, -l 50 was set to help find the correct kcov in order to obtain a more accurate error rate.

```
$tools/genomescope2.0/genomescope.R -i $pf.k31.hist -k 31 -o $pf.gs2 --fitted_hist -l 50
```

To compensate for the known GA-microsatellite dropouts in PacBio long-reads, coverage dropouts were evaluated on HG002v1.1 for short-reads with `bedtools genomecov` to find overall regions with less than 2 reads aligned. For these short reads, we used the bam files described in the next section “Mapping based evaluation”, and ran bedtools genomecov with the usage:

```
# regions with coverage < 2x
bedtools genomecov -bga -ibam $pf.dedup.bam | awk '$4<2' | bedtools merge -i - >
$pf.dedup.cov_lt_2.bed
# exclude rDNA gaps
bedtools subtract -A -a $pf.cov_lt2.bed -b v1.1.gap.bed > $pf.cov_lt2.no_gap.bed
```

The overall results are summarized in the table below (and in **Table S17**):

| Platform | Het (%) | Kcov (het. peak) | Error rate (%) | Model fit | Genome size est. | Bps with coverage < 2x |
|---|---|---|---|---|---|---|
| Element Aviti UltraQ | 0.25% | 34.0 | 0.06% | 0.82 | 2,908,208,748 | 2,180,013 |
| Illumina HiSeq 2x250, | 0.27% | 30.7 | 0.26% | 1.06 | 2,903,028,543 | 1,012,551 |
| PacBio Onso | 0.29% | 22.5 | 0.03% | 0.74 | 2,904,805,881 | 12,216,638 |
| PacBio Revio with SPRQ | 0.28% | 50.6 | 0.08% | 1.08 | 2,866,227,392 | |
| PacBio Revio HiFi | 0.23% | 34.0 | 0.09% | 0.42 | 2,923,638,455 | |

From the GenomeScope2 results, we noted that the SPRQ chemistry had a slightly lower error rate than the HiFi Revio reads without SPRQ. Among short-reads, PacBio Onso data had the smallest error rate, but we also found it had the largest sequencing dropouts in GA-enriched regions compared to other short-read platforms, consistent with other PacBio platforms. Therefore, 31-mers from the next best short-read platform, Element Aviti UltraQ, were chosen to act as a complement to SPRQ read-derived 31-mers for the most accurate and complete representation of the genome. A hybrid database was built, of the same type as was used for evaluating T2T-CHM13[59].

```
meryl union-sum [ greater-than 1 AVITI_UltraQ_Std.k31.meryl ] [ greater-than 1
Revio_SPRQ_30hr_PacBio.k31.meryl ] output SPRQ_union_UltraQ.meryl
```

Subsequently, Merqury (commit ed8c3ba3ea8897d9151a7da8772cd5d83fbce474) was used to re-evaluate HG002v1.1 and all prior assembly versions and benchmark assemblies to obtain a QV score.

```
$MERQURY/_submit_merqury.sh SPRQ_union_UltraQ.meryl $asm1.fa.gz $asm2.fa.gz $out
```

The *_only.bed files contain the locations of 31-mers in the v1.1 assembly that are not found in the `SPRQ_union_UltraQ.meryl` set, and represent likely errors in the assembly. A final bed file consisting of these "error kmer" loci was created by merging regions within 5kb of each other, and this file (containing 314 regions with a total of 205,108 base pairs, and 293 regions with a total of 183,143 base pairs outside the rDNA arrays, **Table S6**) was used in the final issues track (see "Consolidated issues track").

```
cat *_only.bed | sort -k1,1V -k2,2n | bedtools merge -d 5000 -i - >
v1.1.sprq_elmt_hybrid.error.mrg5kb.bed
```

For haplotype evaluation, hamming distance was collected from Merqury results using HG002 Illumina HiSeq2500 2x250 31-mers, along with 31-mers from HiSeq2500 2x250 reads for the parental genomes, HG003 (paternal) and HG004 (maternal).

```
out=${name}.ilmn
hapmers=$out.hapmers.count
```

```
cat $hapmers | awk 'NR>1 { if ($3 > $4) { err=$4; } else { err=$3; } total=$3+$4; \
  if (total==0) { print $0"\t"err"\t"total"\t0.00"} \
  else { print $0"\t"err"\t"total"\t"(100*err)/total } }' > $hapmers.hamming
# Per-haplotype hamming distance
grep $asm1 $hapmers.hamming | awk -v asm=$asm1 '{err+=$(NF-2); total+=$(NF-1);} END {print
asm"\t"(100*err)/total}'
if ! [[ -z $asm2 ]]; then
  grep $asm2 $hapmers.hamming | \
  awk -v asm=$asm2 '{err+=$(NF-2); total+=$(NF-1);} END {print asm"\t"(100*err)/total}'
  cat $hapmers.hamming | \
  awk -v asm="Both" '{err+=$(NF-2); total+=$(NF-1);} END {print asm"\t"(100*err)/total}'
fi
```

After inspection, the majority of discovered switch errors were found to reside in immunoglobulin genes (IG) or TCRs, and therefore were removed from our list of ‘switch errors’. The IG and TCR genes were identified as described below using gene annotations on hg38, lifted to the HG002v1.1 assembly. Excluding these loci, there were 803 regions with a total of 32,014 base pairs, or 794 regions with a total of 31,603 base pairs outside the rDNA arrays.

```
wget https://s3-us-west-2.amazonaws.com/human-
pangenomics/T2T/HG002/assemblies/annotation/VDJregions/HG002.VDJ.lifted_to_v1.1.bed
bedtools subtract -A -a v1.1.hap_switches.bed -b HG002.VDJ.lifted_to_v1.1.bed >
v1.1.hap_switches.noVDJ.bed
```

### Mapping based evaluation

The following read datasets were mapped to HG002v1.1 using the same parameters as for long and short reads in the previous rounds of polishing:

PacBio reads
- PacBio Revio HiFi DeepConsensus v1.2, 3 cells, one 30hr run and two 24 hr runs
- PacBio Revio with SPRQ, 2 cells, 30 hr

ONT reads
- ONT R10.4 ULK q28 (downloaded from epi2me, “An experimental extremely high-accuracy, ultra-long sequencing kit”, 2023)
- ONT R10.4 ULK lc24 (downloaded from epi2me, “Nanopore-only T2T assembly of a human genome”, 2024)

Short-reads
- Element AVITI UltraQ Standard 400bp insert, 2x150 (80x)
- Illumina HiSeq2500 2x250, GIAB (60x)

The PacBio reads were filtered to include only primary alignments with no supplementary alignments. The ONT data were filtered to include primary alignments, discarding reads shorter than 10 kbp. In addition, the q28 and lc24 reads were filtered for mapped identity >85% and >95%, respectively, to remove spurious alignments of lower quality reads using filt_bam_len_idy.py (https://github.com/arangrhie/T2T-Polish/blob/master/coverage/filt_bam_len_idy.py).

It should be noted that three of these read datasets (the PacBio Revio with SPRQ reads, the ONT R10.4 ULK “lc24” reads, and the Element AVITI UltraQ reads, were not available until after the release of T2T-HG002v1.1, and therefore provide an independent assessment of the accuracy of the HG002v1.1 assembly.

#### T2T-Polish issues

After filtering the bam files, paf files were re-generated, and issues.sh from the T2T-Polish pipeline (https://github.com/arangrhie/T2T-Polish/blob/master/coverage/issues.sh) was run twice, once with the PacBio alignments and once with the ONT alignments, using the SPRQ+Element hybrid 31-mers described above.

An intersection of the PacBio and ONT issues.bed files was considered as the final set of ‘T2T-Coverage issues’ to avoid coverage biases posed in GA enriched regions of the PacBio reads.

```
issues.sh hg002v1.1_pb_hifi_sprq.paf PB_HiFi_SPRQ v1.1 HiFi /path/to/pattern
issues.sh hg002v1.1_ont_q28_lc24.paf ONT_Q28_LC24 v1.1 ONT /path/to/pattern

# merge within 1kb
bedtools merge -d 1000 -i hg002v1.1_pb_hifi_sprq.issues.bed >
hg002v1.1_pb_hifi_sprq.issues.mrg1k.bed
bedtools merge -d 1000 -i hg002v1.1_ont_q28_lc24.issues.bed >
hg002v1.1_ont_q28_lc24.issues.mrg1k.bed

bedtools intersect -u -a hg002v1.1_pb_hifi_sprq.issues.mrg1k.bed -b
hg002v1.1_ont_q28_lc24.issues.mrg1k.bed | sort -k1,1V -k2,2n | grep -v "chrM" >
pb_and_ont.issues.mrg.bed
```

Outside of the rDNA arrays, these T2T-Polish-discovered issues consist of 3 regions with a total of 44,652 base pairs.

#### NucFlag

NucFlag v0.3.3 was applied to the HiFi and ONT read data to generate the first and 2nd most frequent tracks and an output bed file using the following parameters:

```
nucflag \
-i hg002v1.1_pb_hifi_sprq.bam \
--output_cov_dir nucflag_freq \
-o nucflag_PB_HiFi_SPRQ.bed \
-p 24 \
-t 24
```

First and second most frequent allele tracks were generated from the bed output in the nucflag_freq as below and uploaded to the browser:

```
for seq in $(cut -f1 $sizes)
```

```
do
 echo $seq
 inputs=`ls $in_dir/${seq}* | sort -k1,1V`
 zcat $inputs | awk -v seq=$seq -v out=$out \
   'BEGIN { header="fixedStep chrom="seq" start=1 step=1"; \
  print header >> out"_1st.cov.wig"; print header >> out"_2nd.cov.wig" } \
  {if ($1!="position") {print $2 >> out"_1st.cov.wig"; print $3 >> out"_2nd.cov.wig"; } }'
done
wigToBigWig ${out}_1st.cov.wig $sizes ${out}_1st.cov.bw
wigToBigWig ${out}_2nd.cov.wig $sizes ${out}_2nd.cov.bw
```

For ONT reads, an experimental version of NucFlag branch feature/v1.0 was applied based on communication with the author. This version detects indel mismatches in addition to SNP based mismatches for more sensitive flagging of misassembly.

```
git clone https://github.com/logsdon-lab/NucFlag.git --branch feature/v1.0 NucFlagv1_0 --depth 1
cd NucFlagv1_0
make venv && make build && make install
source venv/bin/activate

nucflag -i hg002v1.1_ont_q28_lc24.bam \
-f /data/Phillippy/projects/HG002_diploid/assemblies/v1.1.fasta \
-o nucflag_Q28_LC24.indel.bed \
-c /data/Phillippy/tools/NucFlag/NucFlagv1_0/nucflag_ont.toml  \
-t $SLURM_CPUS_PER_TASK
```

An intersection of the two missassembly bed files (`nucflag_PB_HiFi_SPRQ.bed` and `nucflag_Q28_LC24.indel.bed`) was used for further evaluation.

```
bedtools intersect -u -a pb_hifi_sprq/nucflag_PB_HiFi_SPRQ.bed -b
ont_q28_lc24/nucflag_Q28_LC24.indel.issue.bed | grep -v "chrM" | sort -k1,1V -k2,2n >
NucFlag_PB_ONT.indel.issues.bed
bedtools merge -d 100 -i NucFlag_PB_ONT.indel.issues.bed >
NucFlag_PB_ONT.indel.issues.mrg100bp.bed
```

Outside of the rDNA, this intersected set of NucFlag regions had 237 regions with 4,897,713 base pairs.

#### Visualization of short-read alignments

Short-reads were mapped using the T2T-Polish bwa pipeline ([https://github.com/arangrhie/T2T-Polish/blob/master/bwa/bwa.sh](https://github.com/arangrhie/T2T-Polish/blob/master/bwa/bwa.sh)).

```
bwa mem -t $cpu $ref $r1 $r2 > $tmp/$out.sam
samtools fixmate -m -@$cpu $out.sam $out.fix.bam
samtools sort -@$cpu -O bam -o $out.bam -T $out.tmp $out.fix.bam
samtools index -@$cpu $out.bam
# mark duplicates and remove them
samtools markdup -r -@$cpu $out.bam $out.dedup.bam
```

```
samtools index -@$cpu $out.dedup.bam
# collect primary alignments
samtools view -@$cpu -F0x100 -hb -o $tmp/$out.dedup.pri.bam $out.dedup.bam
```

These short-read bam files were used, in addition to the merged PacBio and ONT bam files used in the T2T-Polish coverage and NucFlag analysis, for visual inspection of randomly chosen regions using IGV. The IGV session files used for this are available via the HG002-issues github repository[108].

#### Prior known issues

Issues which had been flagged for patching while making corrections to HG002v1.0.1 but were not correctable (see "Polishing and patching to create v1.1 from v1.0.1") were lifted from HG002v1.0.1 to HG002v1.1 (see "Chain files to and from the HG002v1.1 assembly") and included in the track file "hg002v1.1_issues_and_excluded_regions.merged.bed", which can be viewed and downloaded in the HG002v1.1 browser as the "v1.1 Suspicious Regions" track in the "Assembly and Validation" group. This set of prior known issues, carried over from v1.0.1, was included in the creation of the consolidated "v4" issues track for T2T-HG002v1.1 (see next section) that is displayed in the browser and was used to exclude regions from consideration whenever using T2T-HG002v1.1 as a genome benchmark for the analyses in this manuscript. Briefly, the prior known issue regions fall into the following categories: 1) NIST exclusion regions discovered using DeepTrio, 2) Regions called by DeepPolisher, 3) Uncorrected Sniffles SV calls, and 4) issues submitted through the HG002-issues github repository[108].

#### Consolidated Issues Track

Five BED files were integrated to produce the final "v4" issues track for v1.1. The following files described in the above sections were included for merging:

- `NucFlag_PB_ONT.indel.issues.mrg100bp.bed`
- `v1.1.sprq_elmt_hybrid.error.mrg5kb.bed`
- `v1.1.hap_switches.noVDJ.bed`
- `pb_and_ont.issues.mrg.bed`
- `hg002v1.1_issues_and_excluded_regions.merged.bed`

Regions in these five files were merged using "bedtools merge -d 1000" into a grand "dirty" BED file of issues, after which each merged regions was assigned to a category by comparison back to the component BED files with bedtools intersect, removing regions from the "dirty" file and assigning them to the largest category with which they overlapped.

After merging and categorization, 3,172 v1.1 regions were flagged as issues, comprising 38.8 Mbp (0.65% of diploid HG002, excluding MT) (**Table S6 and Table S7)**. Note that due to the inclusion of padded extensions and merging across error-free regions, as well as the inclusion of possible false-positive quality issues predicted by the various programs, the total bases flagged in hg002v1.1_issues.v4.bed (available on AWS at https://s3-us-west-

2.amazonaws.com/human-pangenomics/T2T/HG002/assemblies/annotation/assemblyissues/hg002v1.1_issues.v4.bed) is likely to be an over-estimate of the error in the v1.1 assembly. In this manner, to provide a reliable benchmark, we erred on the side of caution and flagged any suspicious regions in the assembly.

#### Annotation and browser resources for T2T-HG002v1.1

We have made the two haplotypes of T2T-HG002v1.1 available in the “T2T Genomes” UCSC browser assembly hub as a centralized repository for annotations of the T2T-HG002 assembly (**Figure S4**).

#### Gene annotation

We generated annotations for the T2T-HG002v1.1 assembly (hereafter HG002v1.1) by mapping genes and transcripts from the T2T-CHM13 annotation onto HG002v1.1. The reference annotation used here (JHU RefSeqv110 + Liftoff v5.2) was originally created by mapping the RefSeq (v110) reference annotation of the GRCh38.p14 assembly onto the T2T-CHM13v2.0 assembly using Liftoff[62] and Comparative Annotation Toolkit (CAT)[122], followed by manual curation. To annotate HG002v1.1, we adopted a two-pass approach to handle unusually challenging regions separately. These regions include the V(D)J gene segments and the ribosomal DNA (rDNA) arrays, which have features that tend to cause errors when trying to map them from one assembly to another. To avoid these errors, we created masked versions of the target genomes, in which the V(D)J regions and the rDNA arrays were replaced with Ns.

To annotate putative V(D)J regions, we used Liftoff to project V(D)J gene annotations from the RefSeq GFF of the T2T-CHM13 assembly onto each haplotype of the v1.0.1 assembly. These annotations were then lifted again to the corresponding haplotypes of HG002v1.1. The set of V(D)J genes included TRA, TRB, TRG, IGL, and IGH. For the IGK locus, where both constant regions were ablated due to the presence of IGKDEL, we defined the locus using the start coordinate of IGKDEL and the end coordinate of IGK. rDNA arrays were identified by aligning a reference 45S rDNA sequence to the assembly using nucmer with parameters --maxmatch -l 31 -c 100. Only alignments longer than 1,000 bp and with greater than 96% sequence identity were retained, resulting in 51 putative 45S rDNA copies on the MAT chromosomes and 81 on PAT. Note that each 45S rDNA copy encompasses 18S, 5.8S, and 28S rDNA subunits, as it encodes a precursor rRNA that is subsequently processed into mature rRNA subunits.

We treated each haplotype of the HG002v1.1 assembly (MAT and PAT) as a separate haploid genome and processed each independently (including chrX in the MAT run and chrY in the PAT run), because Liftoff is designed to work with haploid assemblies. For the first pass in our process, we used Liftoff to map all features from the reference annotation, excluding additional gene copies, rDNA array rRNA genes, and V(D)J gene segments, onto a masked version of the MAT and PAT assemblies. We ran Liftoff with the parameters -chroms chroms.mat/pat.txt -copies -sc 0.95 -exclude_partial -polish. This initial pass produced annotations of 59,441 genes on MAT and 57,770 genes on PAT, encompassing protein-coding genes, lncRNAs, and other gene types.

The second pass focused solely on mapping the rDNA arrays. In total, 219 rDNA units, comprising 876 rRNA genes, were extracted from the T2T-CHM13 reference annotation. For this mapping step, we also included 7 rRNA gene annotations that occur outside the typical rDNA array structure but still within the acrocentric arms. To ensure that rRNA genes were mapped as complete units, consisting of 45S, 18S, 5.8S, and 28S genes, in that order, we added custom "unit" features to the rDNA annotations. These "unit" features served as parent elements to the four rRNA genes, ensuring that the four genes would be mapped together. To enforce this structural constraint, we used the -f features.txt parameter in Liftoff to restrict annotation to regions where all four rRNA genes co-occur in the correct order. All other Liftoff parameters remained identical to those used in the first pass. This second pass resulted in the annotation of 21 rDNA arrays (84 rRNA genes) on MAT and 28 arrays (112 rRNA genes) on PAT.

The annotations generated by the first and second passes of Liftoff were merged and sorted using gffread[123] with parameters -O -F –keep-exon-attrs. Before merging, we used bedtools[124] intersect with the -wa -wb flags to confirm that no genes from the first-pass annotation overlapped with those from the second pass. In total, this two-pass process yielded 59,525 genes on MAT and 57,882 genes on PAT.

After running Liftoff, we applied several post-processing steps to improve the quality of the annotated genes and transcripts. One key step involved correcting coding sequence (CDS) features to ensure that their lengths were divisible by three, thereby maintaining proper codon structure. Liftoff does not enforce this constraint, and factors such as minor mapping artifacts, assembly issues, or biological variants (e.g., small insertions or deletions of 1–2 bp) can result in the annotation of CDS features whose lengths are not divisible by three, necessitating trimming. While some of these cases may reflect true frameshifting variants, others arise from technical limitations or ambiguities. We manually inspected a subset of these cases but did not perform a systematic analysis of the underlying sequence contexts; therefore, not all trimmed CDSs should be interpreted as functionally compromised or frameshifted.

To restore the correct codon structure and ensure that the annotated CDS accurately reflects the translated region of a protein-coding gene (with the final three bases corresponding to the last codon), we identified CDS features with lengths not divisible by three and trimmed the excess bases, where bases were trimmed from the end if the first three bases form a valid start codon, and trimmed from the start otherwise. This correction was applied to 586 CDS features on MAT and 534 on PAT. Note that CDS is a transcript-level feature. The sequence responsible for a problematic CDS annotation often spans exon(s) shared by multiple isoforms of the same gene, necessitating trimming across all affected transcripts. Accordingly, the number of genes affected by this trimming procedure is much smaller (e.g., 219 in MAT).

We then extended CDS features that were missing a stop codon to the first in-frame stop codon downstream of the Liftoff-mapped translation termination site. While premature termination codons (PTCs) lead to truncated proteins and potential loss of function (LoF), downstream stop codons that produce elongated proteins are generally considered less deleterious. This procedure resulted in extending 299 CDS features on MAT and 311 on PAT.

To further improve gene annotation, we used Lifton's extra-copy search submodule (Chao et al., 2025) to run miniprot[64], aligning protein sequences from the GRCh38 MANE (v1.4) annotation set to the genome. Protein-to-genome alignments are particularly useful for identifying genes with low nucleotide-level conservation but preserved protein sequences. After generating alignments with miniprot, we compared the resulting annotations to the existing Liftoff-based HG002v1.1 annotations. We filtered out any features that overlapped ≥10% of an existing gene or spanned more than two adjacent gene loci, ensuring that no two genes were annotated in the same genomic location.

Because protein-to-genome alignments only capture CDS regions, we manually added exon, transcript, and gene features to preserve the correct hierarchical annotation structure. If miniprot identified an additional copy of a gene already annotated by Liftoff, we retained the same gene identifier and marked the copy using the "extra_copy_number" attribute. Otherwise, we assigned a new gene identifier. This procedure added 30 genes to MAT and 21 to PAT.

Lastly, to assign gene identifiers, we introduced a T2T-HG002v1.1-specific ID system for all genes and transcripts in the annotation. Unlike reference gene curation efforts such as RefSeq[125], GENCODE[126], or CHESS[127], which sometimes use shared IDs across genomes, this assembly-specific system ensures that each ID refers to a unique object within the HG002v1.1 assembly. Gene IDs follow the format: hg002_[chromosome]_[haplotype]_[gene ordinal number]
where the gene ordinal number reflects the gene's position on a given chromosome counting from left to right; e.g., the 50th gene will have ordinal number 50. Transcript IDs follow the format: [gene_id].[transcript_ordinal_number] where the transcript ordinal number indicates the order of isoforms for a given gene. All features are also tagged with their source gene or transcript ID (i.e., identifiers from the T2T-CHM13 reference annotation), enabling users to track them back to their source in the reference. The current release of the gene annotation excludes the mitochondrial chromosome and V(D)J gene segments.

#### Identification of haplotype-specific genes

We compared haplotype-specific gene copy numbers using HUGO gene symbols to ensure that we were counting only duplicate genes. Genes were considered copies of each other if they shared the same HUGO symbol (**Table S8**). Using these copy numbers, we identified genes exclusive to only one of the two T2T-HG002v1.1 haplotypes, referred to as haplotype-specific genes. In total, we found 832 distinct genes exclusive to MAT, of which 24 were autosomal, and 79 exclusive to PAT, of which 31 were autosomal. As expected, the majority of haplotype-specific genes were on sex chromosomes.

Since Liftoff/LiftOn, when used to project gene annotations from T2T-CHM13 onto HG002v1.1, is primarily based on sequence similarity, it is possible that close paralogs were annotated in place of a specific gene, potentially resulting in false positive haplotype-specific gene calls. To address this, we aligned the protein sequences encoded by each putative haplotype-specific gene to all proteins on the opposite haplotype using BLASTP. Alignments were filtered using a stringent e-value threshold of $1.0\times10^{-10}$, ≥ 90% identity, and ≥ 90% query/target coverage. If a gene had more than one match on the other haplotype, it was excluded from the haplotype-specific set.

After filtering, the final list of haplotype-specific genes included 14 MAT-only autosomal genes, and 12 pat-only autosomal genes (**Table S11**).

### FIRE analysis of Fiber-seq data for HG002

Using Fiber-seq data generated on HG002 (SRX24951052), Fiber-seq Inferred Regulatory Elements (FIRE) v0.04 was applied to identify regulatory elements from long-read Fiber-seq data using the methods described in Vollger *et al.*[74]. In brief, FIRE utilizes semi-supervised machine learning to identify MTase-sensitive patches (MSPs) that represent regulatory elements on single chromatin fibers. The methodology utilizes the Mokapot framework and XGBoost algorithms to classify MSPs as likely regulatory elements, assigning each FIRE element an estimated precision value that indicates the probability of being a true regulatory element. This approach provides single-molecule resolution of chromatin accessibility, analogous to a long-read version of DNaseI/ATAC-seq.

Peak calling was performed by identifying FIRE score local maxima with FDR values below a 5% threshold, with peak boundaries determined by the median start and end positions of underlying FIRE elements. False discovery rates were calculated by shuffling fiber locations across the genome and recalculating FIRE scores, defining FDR as the ratio of bases with shuffled scores above a threshold to bases in unshuffled data. The analysis generated multiple track types, including standard FIRE peaks, wide peaks (merged regions within one nucleosome length), coverage tracks showing MSPs, FIREs, and nucleosomes, as well as haplotype-resolved percent accessibility measurements.

### Repeats and transposable element sequences

Transposable element-derived sequences were annotated using RepeatMasker v4.1.7-p1 and a custom library comprising of curated models in Dfam 3.7[128] and those generated as part of the

T2T-CHM13[26], ape sex chromosomes[34], and HG002 chromosome Y[24] analyses. The aforementioned repeat models were appended to the existing RepeatMasker library as follows:

```
#Make a directory for the new libraries
mkdir ~/TEproject/RMplusY_XY_CHM13/

#Copy the RepeatMasker libraries to a new location
cp -r /usr/local/RepeatMasker-4.1.2-p1/Libraries/ ~/TEproject/RMplusY_XY_CHM13/

#Append the embl file to the existing library
#famdb.py -i ~/TEproject/RMplusY_XY_CHM13/Libraries/RepeatMaskerlib.h5 append
RMplusY_XY_CHM13.embl --name 'RMplusY_XY_CHM13_library'

#Run RepeatMasker with the appended library
RepeatMasker -libdir ~/TEproject/RMplusY_XY_CHM13/Libraries/RepeatMaskerlib.h5 -s -species
v1.1_hg002.fasta -pa 14 -a
```

This process allows the most sensitive and comprehensive search stages for human repeat detection when using the species flag, but also includes the curated repeats from other primate projects not currently in the Dfam database.

### Centromeric satellite sequences

Centromeric satellites were annotated using the CenSat workflow. The workflow is written in workflow description language (WDL), and full code plus additional documentation can be accessed via github (https://github.com/kmiga/alphaAnnotation/tree/main).

Alpha satellites were annotated using a modified version of a HumAS-HMMER (https://github.com/enigene/HumAS-HMMER). This workflow annotates alpha satellite monomers from a database of hidden markov models (HMMs). Alpha monomer annotations were merged into summary bins (active array, inactive higher order repeat (HOR), diverged HOR, and monomeric).

Ribosomal arrays were annotated using HMMs based on the first and last 700 bp of the rDNA repeat unit (GenBank accession U13369.1)[129] as well as two of the rDNA genes; 18S (NCBI Reference Sequence XR_007084227.1) and 5.8S (NCBI Reference Sequence XR_007084259.1)[130]. These annotations were then merged to create a complete summary annotation. Scaffolding gaps (sequences of Ns) were annotated using Seqtk gap (https://github.com/lh3/seqtk) to provide complete annotation coverage of rDNA arrays where the rDNA gaps exist. This annotation of T2T-HG002v1.1's rDNA sequences was used for the locations of the rDNAs for other rDNA analyses in this manuscript (e.g., "Evaluation of HG002v1.1", and "Fraction of HG002v1.1 that is inaccessible to variant-based benchmarks").

Classical satellites (HSATII and HSATIII) were annotated using a script previously described in Altemose et al. 2022[27], which uses a database of human specific kmers. Annotations were then merged to create a summary annotation over regions where strand switching breaks the annotation.

The final subset of centromeric satellites (HSat1A, HSat1B, bSats and gSats, and smaller species such as SST1, SATR, and ACRO) were annotated using RepeatMasker (Smit, AFA, Hubley, R & Green, P. RepeatMasker Open-4.0. 2013-2015 http://www.repeatmasker.org). Relevant satellite monomers were extracted from the RepeatMasker output and merged to summarize larger arrays. Strand information for all cenSat satellites, where available, were recorded in a separate file hg002v1.1.SatelliteStrandv2.0.bed.

The final CenSat annotation set was created by compiling and automatically curating these satellite annotations described above, including resolving overlaps, filtering out small satellite arrays (<2kb) and merging incomplete annotations. The final step identified the centromere transition (CT) regions, which were determined by merging satellite annotations within 2MB and intersected with the location of the active alpha satellite array. This provided tiled annotation of regions within the centromere transition that don't have satellite annotations.

#### Segmental duplications

Segmental duplications (SDs) in the HG002v1.1 assembly were annotated as follows: The maternal and paternal haplotype genomes were comprehensively masked in terms of repeats using TRF[131] v.4.1.0, RepeatMasker[132] v.4.1.5 , and Windowmasker[133] (v2.2.22) with the commands:

```
asm=hg002v1.1.fasta
trf $asm 2 7 7 80 10 50 2000 -l 30 -h -ngs
RepeatMasker   -s   -e   ncbi   -xsmall   -species   human $asm
windowmasker -mk_counts -mem 16384 -smem 2048 -infmt fasta -sformat obinary -in $asm -out
asm.count &&; windowmasker-infmt fasta -ustat asm.count -dust T -outfmt interval -in $asm -out
asm.interval
```

For the maternal haplotype, we concatenated all autosomes from this haplotype with chromosome X and Y to also consider interchromosomal SDs from the two sex chromosomes. Likewise, paternal autosomes and the two sex chromosomes were examined for SDs altogether. Using the repeat-softmasked genomes, SDs were analyzed using SEDEF[134] (v1.1). SDs were further filtered for the pairwise sequence identity >90%, length > 1 kbp, and satellite content <70%.

#### Subtelomeric regions

The annotation of human subtelomeres includes the identification and organization of subtelomere repeat elements (SREs). The SREs are subtelomeric DNA segmental duplications, defined as genomic DNA segments greater than 1 kb and greater than 90% similar in nucleotide sequence that are present in two or more subtelomeres. Subtelomeres are defined operationally as the most distal 500kb human DNA segments at each chromosome end. SRE regions contain mosaic patchworks of segmental duplications called paralogy blocks (pblocks), which bear high similarity to discrete segmental duplication segments occurring in multiple subtelomeres[135,136]. The identity and organization of paralogy blocks is highly polymorphic and haplotype-specific at many subtelomeres[137], and may play a role in cis-regulation of single telomere and haplotype-specific telomere lengths[138,139]. Representative pblock sequences from GRCh38 were used to identify most pblocks from HG002v1.1 SRE regions, but eleven new SRE pblocks were also identified in the HG002v1.1 assemblies.

Representative sequences for existing GRCh38-derived pblocks were masked (with Repeat Masker (Smit, AFA, Hubley, R & Green, P. RepeatMasker Open-4.0. 2013-2015 http://www.repeatmasker.org) and Tandem Repeats Finder[131] software run under default parameters) and then aligned to the reference sequence using BLASTn v.2.13.0+[140], requiring a minimum of 90% identity and 100 bp alignment length. Groups of alignments (localized, ordered, co-directional, greater than 1kb chain-length) form a mapping location of a pblock to the reference. If a pblock aligned only partially with a given segment of the reference sequence, then if possible, it is extended by pairwise alignment of the unmasked p-block sequence to the flanking segment(s) of the reference. If neighboring pblocks are overlapping, then the one with the highest percent identity to the reference in the overlapping area is selected.

After all GRCh38-derived pblocks were mapped to HG002v1.1, remaining subtelomere regions of HG002v1.1 were investigated for potential new pblocks. These regions included gaps greater than 1kb between existing mapped pblocks within a subtelomere, and the subtelomere reference areas towards the centromeric side of the subtelomere region (<500kb), where no existing pblock mapped. The investigation was carried out by masking (with Repeat Masker and Tandem Repeats Finder) these candidate pblock regions and aligning them to the complete HG002v1.1 reference. If a segment of a subtelomere maps to multiple subtelomere areas (<500 kb from telomeres), it was annotated as a valid new subtelomeric pblock.

#### Representation of additional HG002 rDNA units

The rDNA arrays are the most complex regions to assemble in the human genome and, as a result, are the only gaps remaining in the HG002v1.1 reference. To provide a representation of additional copies of HG002's rDNA to the benchmark, we've run ribotin, a tool developed specifically for rDNA array assembly[47], develop branch, commit d8a73739d5f7a5de3d27904a18e53d5bbfdf14e9.

```
ribotin-ref -t 10 -r rDNA.KY962518.1.fasta -i m84005_220827_014912_s1.hifi_reads.fastq.gz -i
m84005_220919_232112_s2.hifi_reads.fastq.gz -i m84011_220902_175841_s1.hifi_reads.fastq.gz --nano
all_pass.vhg002v1.fastq.gz --approx-morphsize 45000 -o ribo_epime/
```

Ribotin outputs sequences which represent the most frequent rDNA units in the sample and a graph that represents all different rDNA units present in the sample with their relative order. The rDNA morph consensus sequences and the graph showing their relative order are available at https://s3-us-west-2.amazonaws.com/human-pangenomics/index.html?prefix=T2T/HG002/assemblies/annotation/rdna/hg002_rdnamorphs_v0.1/.

### Creation of a T2T-HG002v1.1 ideogram track

Unless otherwise noted, all input files, along with the conversion tools bigBedToBed and liftOver, were obtained from the UCSC Genome Browser. The T2T-chm13v2.0 (hs1) cytoBandMapped track file was obtained from https://hgdownload.soe.ucsc.edu/gbdb/hs1/cytoBandMapped/ and converted to bed format. Cytoband coordinates were first lifted over to hg002v1.1.mat and hg002v1.1.pat separately using chain files CHM13v2.0_to_hg002v1.1.mat.chain and CHM13v2.0_to_hg002v1.1.mat.chain (see the section "Chain files to and from the HG002v1.1 assembly"). The CenSat annotation track (see "Centromeric satellite sequences") was obtained from [https://github.com/hloucks/CenSatData/tree/main/HG002/v1.1](https://github.com/hloucks/CenSatData/tree/main/HG002/v1.1). The perl script Refine_CytoBand_Liftover.pl[141] was generated to extract centromere, telomere, rDNA, and heterochromatin boundary information from the CenSat track and fasta index files, then to use this information to adjust the boundaries of these regions in the lifted over cytoband tracks, as follows. Centromere band boundaries were placed at the most distal ends of the "active_hor" array annotation (or split HOR arrays, e.g. on chr3 and 4) on each chromosome. The p->q transition was placed at the midpoint between these centromere band boundaries. Centromere-adjacent band coordinates were adjusted to be contiguous. The last band on each chromosome was adjusted to have its end position match the chromosome end position. rDNA stalk band coordinates were adjusted to match the boundaries of the rDNA arrays. The 1q12, 9q12, and Yq12 boundaries were adjusted to the ends of the large HSat2/3 arrays in those regions, and adjacent band coordinates were adjusted accordingly. For any remaining bands with boundaries that failed to lift over, their coordinates were fixed to preserve the relative sizes of that band and its neighboring bands relative to their sizes in T2T-chm13v2.0. For validation, non-gap band sizes were confirmed to be preserved after liftover, and adjacent band coordinates were checked to be contiguous.

### Heterozygous variants and estimated windowed heterozygosity

Comparison of the maternal and paternal haplotypes of autosomes and the pseudoautosomal (PAR) regions of the sex chromosomes is sensitive to the methods used to align homologous sequences to each other as well as to the decisions made when filtering alignments of repetitive regions. In the absence of prior work prescribing methods for this, heterozygous variants and estimates of heterozygosity were computed as follows:

A FASTA file of all maternal HG002v1.1 autosome sequences was aligned to the corresponding file of paternal sequences with minimap2 version 2.26 with the command:

```
minimap2 -a -t2 -x asm5 v1.1.pataut.fasta.gz v1.1.mataut.fasta.gz | samtools view -O BAM | \
samtools sort --threads 2 -T v1.1.mataut_vs_v1.1.pataut.tmp -O bam -o
v1.1.mataut_vs_v1.1.pataut.sort.bam
```

A second "reverse" BAM file was created using the opposite alignment (paternal chromosomes to maternal). The two BAM files were then passed to the GQC script "gethets"[104] to produce a

BED file of heterozygous variant locations and a heatmap-colored track of windowed variant counts.

```
gethets --bam1 v1.1.mataut_vs_v1.1.pataut.sort.bam --bam2 v1.1.pataut_vs_v1.1.mataut.sort.bam --
ref1 v1.1.pataut.fasta.gz --ref2 v1.1.mataut.fasta.gz  --prefix v1.1.hets.non1to1.200k --non1to1
--windowsize 200000 --heatmap
```

In addition, the sex chromosomes were compared using the same minimap2 parameters, and the two directed BAM files were again passed to the gethets program with the same parameters to find heterozygous positions and calculate heterozygosity in the pseudoautosomal regions.

The gethets script filters the alignments passed to it as follows.

1. For each of the two directed BAM files (maternal to paternal, paternal to maternal):
   - Include only alignments that are designated as “primary”
   - Alignments must cover at least 10,000 bp along the reference (target)
2. Find pairs of corresponding alignments. For each included alignment in the first BAM:
   - If a reciprocal matching alignment in the second BAM exists (for which the target has the same start and end coordinates as the alignment’s query coordinates, and the query has the same start and end coordinates as the target start and end, pair the two alignments
   - If no reciprocal alignment exists in the second BAM file, examine all alignments in the second BAM file for which the covered query region intersects the first alignment’s target region. If the intersecting target region’s endpoints in the first file align to the same opposite haplotype positions in both the first and the second BAM file, use the sub-alignments in the intersecting portion as corresponding alignments
3. For all pairs of corresponding alignments, tally heterozygotes as any mismatch, insertion, or deletion within the alignment’s CIGAR string. In addition, calculate the number of these heterozygous positions in windows sized according to the parameter passed to the gethets program.

A BED track reporting all heterozygous positions between the maternal and paternal haplotypes in HG002v1.1, as well as a windowed BED file with heatmap-colored heterozygosity values is displayed and available for download in the UCSC browser assembly hub for HG002v1.1.

### Chain files to and from the HG002v1.1 assembly

Chain files for lifting coordinates to and from the HG002v1.1 assembly were created using the "nf-LO" pipeline described in Rhie et al., Nature 2023[24]. Briefly, we used nextflow to run nf-LO[142] between the HG002v1.1 genome and a haploid target genome (e.g., GRCh38 or CHM13v2.0), then split the chains at all locations where there were unaligned segments longer than 1kbp or gaps longer than 10kbp. Only alignments between homologous chromosomes were retained. The package rustybam[143] was then used to trim overlapping portions of the chains, resulting in one-to-one alignments. The package chaintools[104] was used to invert chains to obtain chain files for the reverse direction.

Chain files used to lift coordinates to and from HG002v1.1 for this manuscript are displayed and available for download in the UCSC browser assembly hub for HG002v1.1.

```
nextflow run nf-LO/main.nf --source $SOURCEFASTA --target $TARGETFASTA --outdir . -profile local
--aligner minimap2 --max_cpus 2 --max_memory 64Gb -resume python chaintools/split.py -c
./chainnet/liftover.chain -o $PREFIX-split.chain python chaintools/to_paf.py -c $PREFIX-
split.chain -t $SOURCEFASTA -q $TARGETFASTA -o $PREFIX-split.paf
awk '{short1=$1; short2=$6; gsub("_MATERNAL", "", short2); gsub("_PATERNAL", "", short2);
if(short1==short2) {print}}' $PREFIX-split.paf > $PREFIX-samechr-split.paf
cat $PREFIX-samechr-split.paf | rb break-paf --max-size 10000 | rb trim-paf -r | rb invert | rb
trim-paf -r | rb invert > $PREFIX.paf
paf2chain -i $PREFIX.paf > $PREFIX.chain
python chaintools/invert.py -c $PREFIX.chain -o $PREFIX.inverted.chain
```

### Fraction of HG002v1.1 inaccessible to variant-based benchmarks

To determine the fraction of the non-rDNA HG002 genome that was not included in the GIAB v4.2.1 read-based small variant benchmark, we lifted the "high confidence regions" of v4.2.1 (available at https://ftp-trace.ncbi.nlm.nih.gov/giab/ftp/release/AshkenazimTrio/HG002_NA24385_son/NISTv4.2.1/GRCh38/HG002_GRCh38_1_22_v4.2.1_benchmark_noinconsistent.bed) onto each haplotype of the HG002v1.1 assembly separately using the GRCh38 to HG002v1.1 haplotype chain files described in “Creation of chain files to and from the HG002v1.1 assembly”.

The total bases in the lifted regions and in the v1.1 genome are presented in **Table S16.** In particular, of the 5,965,910,578 base pairs of non-rDNA sequence in the T2T-HG002v1.1 assembly, only 5,081,210,845 base pairs, or 85.2%, are covered by lifted “high confidence” regions of the GIAB v4.2.1 variant benchmark. In contrast, the “non-excluded” regions of the T2T-HG002v1.1 assembly cover 5,960,629,742, or 99.9%, of the 5,965,910,578 base pairs of the non-rDNA portions of T2T-HG002v1.1 (**Table S4**).

### GIAB v4.2.1 missed regions in various annotated regions of HG002v1.1

We loaded chromosome sizes and the lifted-over GIAB confident regions into separate GRanges objects and used the setdiff function from the GenomicRanges package to identify regions of the HG002 autosomes missing from the GIAB confident set. To assess whether these missing bases overlapped biologically functional elements, we imported gene annotations (HG002v110.JHU.v0.5.withheader.gff), coding sequence (CDS) features, and cenSat annotations (see “Gene annotation”, above) into GRanges objects and intersected them with the missing regions. Set intersections were visualized using the eulerr package in R, where values were converted into named vectors and fitted to Euler diagrams, and the final venn diagram is shown in **Figure 1B**.

### Use of HG002v1.1 as a genome benchmark to evaluate assemblies, reads and variant call sets

### Evaluation of test assemblies using GQC

The availability of a near-perfect diploid benchmark for HG002 allows users to evaluate haploid and diploid genome assemblies by comparing them to the benchmark. To facilitate this and other comparisons, we developed a software tool called GQC, and archived the version used for this manuscript's analyses in the manuscript's software repository[104]. GQC is implemented in python, and includes libraries for aligning sequences and parsing the alignments to determine continuity and consensus accuracy at a local and genome-wide level.

#### Pre-phasing diploid assemblies with GQC

To report quality statistics about assemblies, GQC first pre-phases the assembly scaffolds into maternal and paternal regions ("phase blocks") based on the presence of haplotype-specific kmers present in only one haplotype of the v1.1 benchmark ("assembly-hapmers"). This is done to find phase switches, i.e., places in test assembly scaffolds where the consensus changes from maternal to paternal sequence, or vice-versa. Phase blocks are calculated first by calculating and then mapping the assembly-hapmers to the test assembly using FASTK v1.1 (Gene Myers, https://github.com/thegenemyers/FASTK, accessed February 14, 2025 and archived in the software archive for this manuscript[104]. The assembly-hapmer locations are then used as the observables of a two-state hidden Markov model ("HMM") and the Viterbi algorithm is then used to predict the most probable underlying benchmark haplotype "state" of the assembly at each locus. The predicted states are then used to write a BED-formatted file of phase block regions within each scaffold.

Briefly, the hidden Markov model has a state $S_i$ at each assembly-hapmer position $i$, where $S_i$={'mat', 'pat'} is the haplotype of the position's phase block and is maternal or paternal. In addition, a state $S_i$ "emits" an observable $O_i$ that is also one of "mat" or "pat" with probabilities $1 - \alpha$ (if the state and the observable represent the same haplotype) or $\alpha$ (if the state emits an observable from the opposite haplotype). To allow for phase changes, the model allows the underlying state $S_i$ to alter its value between adjacent assembly-hapmer positions with probability $\beta$ (for a switch to the opposite haplotype state) and correspondingly, the model assigns a probability of $1 - \beta$ for $S_{i+1}$ to remain in the same haplotype state at the following marker. By examining the size of resulting haplotype blocks and the rate of opposite haplotype markers within blocks, we chose $\alpha = 0.05$ and $\beta = 0.01$ as default values for GQC to pre-phase the assembly scaffolds it evaluates. When adjacent assembly-hapmers have a change in state in the resulting state chain, the position exactly between the two hapmer positions is chosen as the boundary between the two phase blocks.

After determining the scaffold phase blocks, GQC aligns all assembly scaffolds separately to the maternal part of the v1.1 benchmark and to the paternal part, then considers phased regions of the assembly one by one, using only the alignments of that region to their same-haplotype region based on the pre-phasing. Since the alignments to the single-haplotype assembly frequently extend beyond the boundaries of the predicted assembly phase blocks, only the portion of each alignment that lies within the phase block is considered in the evaluation (see "Evaluation of assembly accuracy within alignments to the benchmark").

#### Evaluation of assembly's long-range continuity

In addition to reporting on a test assembly's standard continuity metrics (contig and scaffold N50/L50, NG50/LG50, and auNG), GQC calculates statistics with regard to the portions of the assembly's scaffolds that are aligned to the genome benchmark. While N50, NG50, etc. are calculated based on assembly contig and/or scaffold lengths, the aligned statistics NGA50/LGA50, etc. are dependent on the alignments themselves, which are determined using minimap2 with the "asm5" preset by default. Because minimap2 can sometimes be unpredictable with regard to how large an indel it will incorporate into its alignments, GQC breaks alignments at the locations of indels which are 10,000 base pairs or larger in size, and joins consecutive same-strand alignments if the endpoint of the first is closer than 10,000 base pairs to the starting point of the next along both the test assembly sequence and the benchmark. Once alignments are split and combined in this way, NGA, LGA, and auNGA statistics are calculated from the lengths in a manner similar to their counterparts.

#### Evaluation of assembly accuracy within alignments to the benchmark

Within each region of the assembly's phased scaffolds, the alignment to the haplotype matching the region's phase block is used to tally the assembly's errors. Any substitution, insertion, or deletion within the alignment is listed in a BED-formatted output file of errors (and optionally, a VCF-formatted file), and classified as either a "phasing error" when the assembly allele matches the benchmark sequence on the opposite haplotype, or a "consensus error" if the assembly allele doesn't match either benchmark haplotype. The number of errors of each type are used to calculate phred-scaled quality values reported in the program's general statistics output file.

#### Evaluation of mononucleotide run accuracy

In addition to being listed as errors in the consensus analysis, mononucleotide errors are evaluated specifically for the program's mononucleotide accuracy statistics. The HG002v1.1 assembly has 1,990,653 mononucleotide runs of length ten or greater, and for each of these runs that are included within a same-haplotype alignment of the phased assembly scaffolds to the benchmark, runs are classified as either correct (the complete run is in the scaffold and has the same number of bases as the benchmark), wrong haplotype (the complete run is in the scaffold but has a different number of bases than the matching haplotype, but the same number as the benchmark's opposite haplotype), wrong length (the complete run is in the scaffold but has a different number of bases than either haplotype of the benchmark), or erroneous (the run contains other sequence beside the single base within the mononucleotide sequence).

#### Assemblies used for comparison and evaluation with HG002v1.1

We used other high quality assemblies of HG002 for validation of HG002v1.1 as well as for demonstration of HG002v1.1's value as a genome benchmark. Our motivation in selecting this particular group of assemblies was to evaluate using the benchmark the improvement of assemblies over several years' time. Details of these assemblies' availability for download and citations of manuscripts describing them are listed in **Table S12**. Briefly, the HG002 assemblies used in this work's evaluation were:

1. The "Ash1v2.0" haploid assembly[53] with contigs filtered to include only the main chromosomes:
```
for chrom in `awk -F"\t" '{print $1}' Ash1_v2.0.fa.fai | grep 'chr' | \
grep -v 'random' | sort`; do samtools faidx Ash1_v2.0.fa $chrom >> \
Ash1_v2.0.mainchroms.fasta; done
```

2. The hifiasm diploid assembly published in Cheng *et al.*, 2021[54].

3. The "year1hprc" diploid assembly downloaded from the HPRC AWS site[10,41]. This assembly was created with the hifiasm assembler using data similar to other samples that were part of the HPRC project's first data release.

4. The verkko diploid assembly published in Rautiainen *et al.*, 2023.

5. The diploid assembly "lc24_medaka_6b4" which was announced at Oxford Nanopore's 2024 London Calling meeting. This was a verkko2 assembly run entirely on ONT reads from multiple separate sequencing kits, including the use of "6b4" chemistry designed to improve the accuracy of homopolymer runs.

### Evaluation of sequencing read accuracy using GQC

#### Datasets used to demonstrate read accuracy benchmarking

Four read datasets were included in the read benchmarking results presented in this manuscript. Information on where they were obtained is in **Table S12**. GQC readbench results, as well as AWS links to the aligned BAM files used to generate them, are in **Table S14.** The first read set evaluated was the high accuracy, ultra-long Oxford nanopore sequence dataset presented by Epi2Me at the Nanopore Community meeting in December, 2023). The announcement of this dataset describes the dataset as:

> Ultra-long libraries of native DNA from GM24385 (HG002) were prepared using a modified Ultra-Long DNA Sequencing Kit V14 motor protein and experimental high-accuracy run conditions. Sequencing was performed on a PromethION instrument to obtain 125 Gbp of sequencing data passing quality filters (read Q-score > Q10, ), with a read length N50 of 91 kbp. This data was basecalled using a bespoke dorado model to yield a median accuracy of Q26.4.

The second read set used in the read benchmarking analysis were HiFi reads sequenced by Pacific Biosystems in May, 2024 with 24 hour movies on a Revio ICS 13 flowcell, then called with DeepConsensus v1.2. The third read set was sequenced by Element Biosciences using their UltraQ (Q50) chemistry with a 400 base pair insert size, and the fourth set was a dataset of PCR-free whole genome sequencing reads, sequenced in 2020 on the Illumina NovaSeq platform, and described in Baid et al. 2020 bioRxiv (https://www.biorxiv.org/content/10.1101/2020.12.11.422022v1).

The “readbench” command in the GQC package (Nancy F. Hansen,  (2025) allows users to evaluate aligned sequence reads for substitution and insertion/deletion discordance rates.

#### Calculation of substitution and small insertion/deletion discrepancies

From a user-supplied BAM file, GQC readbench reports all discrepancies (single nucleotide substitutions and indels) within alignments of reads to the benchmark genome. If the allele displayed on a read matches the opposite haplotype’s allele at a heterozygous site of HG002, the error is tagged as a “phasing” error.

#### Coverage assessment

Based on a BAM file’s coverage and read lengths, GQC readbench’s “--arrivalratecoverage” option sets a bin size estimated to result in a mean 1,000 read starts per bin, and then reports the number of read starts in each non-overlapping bin of that size which is fully included within the non-excluded regions of the HG002v1.1 genome benchmark. These bin arrival rates can then be used to plot cumulative coverage curves like the ones in **Figure 4E** of this manuscript. In addition, GQC’s “--bincoverage” option reports the average coverage of primary read alignments across bins of a user-specified size as well as the GC content within each bin in BED format, using a cumulative sum operation on read start and stop “events” for efficiency, following the methods used by the program mosdepth[144]. These values, printed in BED format, can be used to make plots like the one in **Figure 4F** of this manuscript.

#### Assessment of short tandem repeat accuracy

For regions of the diploid benchmark that are annotated as homopolymer, dinucleotide, trinucleotide, and tetranucleotide runs, GQC readbench assesses the accuracy of a sequence read for that run as follows:

1. First it determines the read positions aligned to the five bases immediately flanking the short tandem repeat (STR) on each side. If there aren’t aligning bases due either to deletion or alignment clipping, it extends the STR-aligned read sequence to include adjacent bases so there are five bases on each side in addition to the STR sequence.
2. If the sequence within the STR is an exact repeat of the motif, it counts the number of bases within the STR sequence and compares it to the benchmark run length. If it matches, and the five flanking bases on each side match the benchmark’s flanking

bases, the read is counted as “CORRECT”. If the run length matches but there are differences in the flanking bases, the read is counted as “FLANKERROR”.
3. If the sequence within the STR is an exact repeat of the motif but its length is different from the benchmark that it is aligned to, and the opposite haplotype of the benchmark is a run of the same length as the read, it counts the read as a “HET” error. If the read STR length is different but doesn’t match the opposite benchmark haplotype, it is counted as a “LENGTHERROR”.

GQC readbench writes files with the read classification for each assessed STR for each assessed read. It then creates a plot of STR accuracy as a function of run length and a histogram plot showing the frequencies of different length errors.

#### Evaluation of base quality scores

If the user-supplied BAM file contains base quality scores, GQC readbench will tally binned accurate bases and errors (reporting substitutions and indels separately) by quality score as it processes alignments. It reports these counts, along with an “effective” phred-scaled quality score for each bin, in a separate output file. QV bin accuracy tallies for the various read data sets were used to generate the plots in **Figure 4D**.

### Evaluation of variant call sets via “constructed genomes”

#### Converting gVCF files to VCF files with high confidence region BED files

Variant callers like DeepVariant output gVCF files containing entries with an "END=" value in the INFO field and a "0/0" genotype with a GQ score for the sample for regions that DeepVariant is calling as a match to the reference. In addition, entries with a "PASS" in the filter field report a genotype and GQ score at variant positions similar to those in a traditional VCF file.

For the non-variant gVCF entries that include an "END=" value, we included the region beginning at the entry's position and ending at its END value in the high confidence region BED file. For passing genotype calls without END values, we also included the region from the entry's position with length equal to the length of the reported REF allele as a high confidence region. Regions with a genotype of "./." were **not** included in the high confidence region BED file.

The gVCFs used in this analysis are publicly available from the NCBI trace archive (**Table S12**).

The commands used to create the high confidence BED files are:

```
# Homozygous reference sites with genotype quality at least $MINQ:
bcftools query -i 'GT="RR" && INFO/END!="."' -f '[%CHROM\t%POS0\t%INFO/END\t%GQ\n]' $GVCF | awk -
F"\t" '$4>=ENVIRON["MINQ"] {OFS="\t"; print $1, $2, $3}' | awk -F"\t" '$2>=0 {print} $2<0
{OFS="\t"; print $1, 0, $3}' > $HIGHCONFBED
# add passing variant/genotype call regions that have GQ at least $MINQ:
```

```
bcftools query -i 'INFO/END="."' -f '[%CHROM\t%POS0\t%END\t%GQ\n]' $GVCF | awk -F"\t"
'$4>=ENVIRON["MINQ"] {OFS="\t"; print $1, $2, $3}' | awk -F"\t" '$2>=0 {print} $2<0 {OFS="\t";
print $1, 0, $3}' >> $HIGHCONFBED
```

High confidence regions were sorted and merged to remove overlaps/redundancy:

```
sort -k1,1 -k2,2n -k3,3n $HIGHCONFBED | bedtools merge -i - > $HCMERGEBED
```

The gVCF file's header and all entries with non-reference genotypes that are also not "./." are included in the VCF variant file that is subsequently used to alter the genome reference file to create a "variant-constructed genome".

The command used to create the VCF file of variants is:

```
gunzip -c $GVCF | awk -F"\t" '$1~/^#/ || ($9~/^GT/ && $10!~/^0\/0/ && $10!~/^\.\/\./) {print}' |
awk -F":" '$1~/#/ || $2=="GQ" && $8>=ENVIRON["MINQ"] {print}' | \
bgzip -c > $VARIANTVCF
```

#### Phasing variant call sets

Because the variant call sets were unphased, we used HiFi reads (**Table S12**) to phase the VCF-formatted variants created in the previous step with HapCUT2[89]. This was to ensure that the variant-constructed genome would represent two single haplotypes, at least on a local level.

HapCUT2 was run using the following commands:

```
extractHAIRS --pacbio 1 --realign_variants 1 --ref $REF --bam $BAM --VCF $VARIANTVCF --out
$PREFIX HAPCUT2 --fragments $PREFIX --VCF $VARIANTVCF --output $PREFIX.haplotype
```

#### Creating and benchmarking a variant-constructed genome

To create two separate haplotypes for the two VCF alleles at autosomal sites, the phased diploid VCF file was split into two haploid VCF files using the following commands:

```
gunzip -c $VCF | grep -v 'RefCall' | awk -F"\t" '$1!~/#/ && $1!~/chrY/ {$NF=gensub(/([0-
9]+)([|/]).*/,"\\1\\2\\1","g", $NF)} {OFS="\t"; print}' | bgzip -c > $HAP1VCF
gunzip -c $VCF | grep -v 'RefCall' | awk -F"\t" '$1!~/#/ && $1!~/chrX/ {$NF=gensub(/([0-
9]+)([|/])([0-9]+).*/,"\\3\\2\\3","g", $NF)} {OFS="\t"; print}' | bgzip -c > $HAP2VCF
```

These haploid VCF files were used with bcftools consensus to create two FASTA-formatted haploid genome files from the reference file against which the variants were called:

```
bcftools consensus -c $CHAIN1 -f $REF -s $SAMPLE $HAP1VCF > $NEWREFPREF.hap1.fasta
bcftools consensus -c $CHAIN2 -f $REF -s $SAMPLE $HAP2VCF > $NEWREFPREF.hap2.fasta
```

and the resulting chain files (produced because the "-c" option was used with bcftools consensus) were used to create BED-formatted files for the high confidence regions against each of the new haploid genomes:

```
liftOver $HQBED $CHAIN1 $NEWBED1 $UNMAPPEDBED1
liftOver $HQBED $CHAIN2 $NEWBED2 $UNMAPPEDBED2
```

Finally, the high-confidence BED files were used to mask their respective genome FASTA files using "bedtools maskfasta":

```
bedtools subtract -a $NEWREFPREF.genome.hap1.bed -b $NEWBED1 > $LQBED1
bedtools subtract -a $NEWREFPREF.genome.hap2.bed -b $NEWBED2 > $LQBED2

bedtools maskfasta -fi $NEWREFPREF.hap1.fasta -bed $LQBED1 -fo $NEWREFPREF.hap1.highconf.fasta
bedtools maskfasta -fi $NEWREFPREF.hap2.fasta -bed $LQBED2 -fo $NEWREFPREF.hap2.highconf.fasta
```

These variant-constructed genome haplotypes ($NEWREFPREF.hap1.highconf.fasta and $NEWREFPREF.hap2.highconf.fasta) were then compared to the hg002v1.1 benchmark and evaluated using the methods described in the section "Evaluating genome assemblies with GQC".

To obtain the values plotted in **Figure S5A-D** and for calculating the figures reported in the main paper, the following shell commands were run on the GQC output files:

```
for file in `ls
GRCh38_GQC/GRCh38_Revio_DV_HiFiPhased.mingq[1234]0/GRCh38_*mingq[1234]0/*.benchcovered.v1.1.bed
CHM13_GQC/CHM13_Revio_DV_HiFiPhased.mingq[1234]0/CHM13_*mingq[1234]0/*.benchcovered.v1.1.bed`; do
  export GENSTATFILE=`echo $file | sed \ 's/.benchcovered.v1.1.bed/.generalstats.txt/'`
  export QUAL=`echo $file | sed 's/.*mingq//' | sed 's:\..*::'`
  export REF=`echo $file | sed 's/_.*//'`
  export COVERED=`awk -F"\t" '{sum += $3-$2} END {print sum}' $file`
  export TOTERRORS=`grep 'Total errors in alignments' $GENSTATFILE | awk '{print $NF}'`
  export SNPERRORS=`grep 'Total substitution errors in alignments' $GENSTATFILE | awk '{print
$NF}'`
  export INDELERRORS=`grep 'Total indel errors in alignments' $GENSTATFILE | awk '{print $NF}'`
  echo -e $REF"\t"$QUAL"\t"$COVERED"\t"$TOTERRORS"\t"$SNPERRORS"\t"$INDELERRORS
done
```

To consider only the consensus quality in the regions of HG002 covered by both the GRCh38-constructed and the CHM13-constructed genomes, the following shell commands were run:

```
for grch38file in `ls
../GRCh38_GQC/GRCh38_Revio_DV_HiFiPhased.mingq[1234]0/GRCh38_*mingq[1234]0/*.benchcovered.v
1.1.bed`; do
  export QUAL=`echo $grch38file | sed 's/.*mingq//' | sed 's:\..*::'`
  export CHM13FILE=`echo $grch38file | sed 's/GRCh38/CHM13/g'`
  export GRCH38ERRORS=`echo $grch38file | sed
's/.benchcovered.v1.1.bed/.errortype.v1.1.bed/'`
  export CHM13ERRORS=`echo $GRCH38ERRORS | sed 's/GRCh38/CHM13/g'`
  echo $QUAL $grch38file $CHM13FILE $GRCH38ERRORS $CHM13ERRORS
  bedtools intersect -a $grch38file -b $CHM13FILE -u >
intersected_covered_regions.gq$QUAL.bed
  awk '$(NF-1)=="SNV" {OFS="\t"; print $1, $2, $3}' $GRCH38ERRORS >
grch38_snv_errors.gq$QUAL.bed
  awk '$(NF-1)=="INDEL" {OFS="\t"; print $1, $2, $3}' $GRCH38ERRORS >
grch38_indel_errors.gq$QUAL.bed
  awk '$(NF-1)=="SNV" {OFS="\t"; print $1, $2, $3}' $CHM13ERRORS >
chm13_snv_errors.gq$QUAL.bed
  awk '$(NF-1)=="INDEL" {OFS="\t"; print $1, $2, $3}' $CHM13ERRORS >
chm13_indel_errors.gq$QUAL.bed
  bedtools intersect -a grch38_snv_errors.gq$QUAL.bed -b
intersected_covered_regions.gq$QUAL.bed -u > grch38_snv_errors.gq$QUAL.intersected.bed
```

```
  bedtools intersect -a grch38_indel_errors.gq$QUAL.bed -b
intersected_covered_regions.gq$QUAL.bed -u > grch38_indel_errors.gq$QUAL.intersected.bed
  bedtools intersect -a chm13_snv_errors.gq$QUAL.bed -b
intersected_covered_regions.gq$QUAL.bed -u > chm13_snv_errors.gq$QUAL.intersected.bed
  bedtools intersect -a chm13_indel_errors.gq$QUAL.bed -b
intersected_covered_regions.gq$QUAL.bed -u > chm13_indel_errors.gq$QUAL.intersected.bed
done

for qual in `echo "10 20 30 40"`; do
  export INTERSECTEDCOV=`awk -F"\t" '{sum += $3-$2} END {print sum}'
intersected_covered_regions.gq$qual.bed`;
  export GRCH38ERRORS=`cat grch38_indel_errors.gq$qual.intersected.bed
grch38_snv_errors.gq$qual.intersected.bed | wc -l`;
  export CHM13ERRORS=`cat chm13_indel_errors.gq$qual.intersected.bed
chm13_snv_errors.gq$qual.intersected.bed | wc -l`;
  echo -e $qual"\t"$INTERSECTEDCOV"\t"$GRCH38ERRORS"\t"$CHM13ERRORS
done
```

### Direct comparison between genome-based and variant-based benchmarking errors

#### Variant benchmark generation and comparison using hap.py

Genome benchmarking and variant benchmarking are theoretically based on the same information, but it is not obvious how they can be compared due to their different representations of accuracy. To explore this, we generated variants using the curated HPRC assembly[40] (**Table S12**) and compared them to a variant benchmark using hap.py. We then compared these variant benchmarking results to accuracy reports for the same assembly evaluated against the T2T-HG002v1.1 genome benchmark.

We generated the genome benchmarking results against HG002v1.1 by running an early version of GQC called “q100bench” on alignments of the curated HPRC assembly to the v1.1 benchmark. The results of this analysis are similar to the results described in the section “Evaluation of test assemblies using GQC”.

To perform the variant benchmarking using hap.py, we aligned the HPRC assembly to GRCh38 with minimap2 (v2.28-r1209) and custom parameters -z200000,10000,200 and called variants using dipcall. Then we ran hap.py using benchmarking best practices[12] to compare the HPRC assembly-based variant calls to a variant benchmark we had generated as part of creating the “HG2-T2TQ100” assembly-based variant benchmark with https://github.com/usnistgov/defrabb available at https://ftp-trace.ncbi.nlm.nih.gov/ReferenceSamples/giab/data/AshkenazimTrio/analysis/NIST_HG002_DraftBenchmark_defrabbV0.019-20241113/).

This hap.py output and the genome benchmarking results described above were used as input for the next two analyses.

#### Overall comparisons between genome benchmarking QV and variant benchmarking precision/recall

We first asked how the overall performance metrics for genome (QV) and variant benchmarking (F1, precision, recall, variants per base, etc) compared to each other for the curated HPRC assembly. We filtered the hap.py output from comparing the HPRC haplotypes to the HG2-T2TQ100 variant benchmark to a subset of stratifications in complex repeats and hard-to-map regions. We then used the same stratifications to subset the genome benchmarking errors after having projected them onto GRCh38, including only those that landed in the small variant benchmarking regions. Errors of length 1 were deemed “SNVs” and the rest “INDELs” which we then compared likewise to the same categories in the variant benchmark across each stratification.

Within each stratification, metrics were computed as follows. The numbers of false negatives (FNs) and false positives (FPs) were taken from the columns “TRUTH.FN” and “QUERY.FP” in the hap.py output. FN+FP was merely the sum of these two. These counts per base were obtained by normalizing them to Subset.IS_CONF.Size in the hap.py output, which corresponds to the number of base pairs within the indicated stratification. QV (genome error per base) was obtained by also normalizing the number of genome errors that fell within the stratification on GRCh38 to Subset.IS_CONF.Size. Precision, Recall, and F1 were taken as-is from the columns “METRIC.Precision”, “METRIC.Recall”, and “METRIC.F1_Score” in the hap.py output. These three metrics were then inverted (1 - metric) to make them directionally similar to QV (i.e., after phred-scaling, higher corresponds to fewer errors). Each metric was then phred-scaled and plotted within each stratification and for SNVs and INDELs. These results were plotted in **Figure 5** and **Figure S6**.

#### Base-level comparison between genome errors and variants

We next asked how variants and genome errors compared to each other at an individual level; for instance, we observed that a single genome benchmarking error often corresponded to multiple variant benchmarking errors, and it is not immediately obvious why this happens. For this analysis, we only considered small variants and not structural variants with respect to GRCh38 (>50bp). In addition, since hap.py does not phase its output, we added phasing back into the hap.py output by using ‘bcftools merge’ to merge it with the phased VCF file created by dipcall. After merging, we directly compared simple variants which had an unambiguous representation, or replayed clusters of variants within ~10bp of each other in order to determine correct phasing. This recovered the phasing of 99.998% (4558745/4558833) of variants.

These variants (in vcf format) were then “unzipped” into two bed-like files corresponding to variants against GRCh38 for each haplotype; specifically we generated files which contained, for each variant position, the REF allele and the ALT allele for both the T2T-HG002v1.1 and the HPRC haplotype (either mat or pat). Hereafter these “split” variants are referred to as “variant alleles” to distinguish from variants in VCF format which often represent a variant relative to two haplotypes. This format enabled us to easily check if the two assemblies had a variant relative to each other (and thus if a genome error should be expected at that position) and if either had a variant relative to GRCh38. Each variant allele was then padded with 50bp on either side.

Next, we needed to project genome errors from the T2T-HG002v1.1 assembly coordinate system to the GRCh38 coordinate system. We first generated .paf files by aligning each of the v1.1 assembly haplotypes to GRCh38 using minimap (with '-c --paf-no-hit --cs -z200000,10000 -xasm5'). We then projected the genome errors using the projection script "project_blocks_multi_thread.py" from the flagger tool[114]. Note that each genome error bed file from GQC (each file corresponded to one of the two HPRC assembly haplotypes) contained coordinates for its evaluated HG002v1.1 haplotype (paternal or maternal); therefore, each error bed file was split into either haplotype, resulting in four bed files that were projected (two for "like-haplotype" alignments, and two for "cross-halotype" alignments where the HPRC assembly mapped to the opposite HG002v1.1 haplotype). These "cross-alignments" were relatively few and analysed separately. We then subset the remaining projected variants to the small variant regions, which excludes structural variants (those >50bp) (https://giab-data.s3.amazonaws.com/defrabb_runs/20241009_v0.018_HG002Q100v1.1/results/draft_benchmarksets/GRCh38_HG002-T2TQ100v1.1-dipz2k_smvar-excluded/GRCh38_HG2-T2TQ100-V1.1_smvar_dipcall-z2k.benchmark.bed). We also excluded any genome errors which had a difference of >50bp between the HG002v1.1 and HPRC assemblies.The remaining genome errors were then compared to the variants.

Next, we intersected variant alleles and genome errors (now both in bed format) using 'intersectBed -a <A> -b <B> -loj'. We performed this intersection both ways (i.e., with "A" and "B" being either variant/genome error or the reverse) for both haplotypes in order to obtain overlaps and non-overlaps in either category. Note that the 50 base pair padding added to the variants meant that this intersection actually found variant alleles/errors "near" each other. For genome errors and variant alleles which intersected, we next attempted to "match" either with the other by first applying the truth and query variants to GRCh38, and then substituting the projected sequence from the T2T-HG002v1.1 and HPRC assemblies to their mapped coordinates in GRCh38 (essentially treating the each genome error as a "variant" relative to GRCh38). A "match" was obtained if the resulting sequence from both truth/T2T-HG002v1.1 and query/HPRC were equal. Note that a "match" may require multiple variants and/or genome errors to be considered at once. In practice, ~43% of variant alleles/errors intersected one-to-one; matches were trivial to determine in these cases. For more complex cases, we used a combinatorial optimization algorithm to replay different combinations of errors and variant alleles. ~11% of these variant alleles contained multiple error/variant allele combinations which matched one-to-one that happened to be near each other. The results of this matching process were summarized in **Figure S7A,B** and **Figure S8A.**

Lastly, we intersected these match results (per haplotype) with the original hap.py vcf file which contained the benchmarking results. This allowed us to determine the number/type of genome errors that matched with the number/type of variants, as well as whether or not a true error (i.e., mismatch between T2T-HG002v1.1 and HPRC) actually resulted in a FP or FN label from hap.py. In total, 41545 variants encoded for a genome error, 27393 had a perfect match, 6761 intersected with a genome error but failed the match algorithm (potential match) and the remainder did not have a hit (due to failed liftover, complex variant representation, potential bugs in hap.py, errors in variant benchmark, or missing genome errors in GQC). In 17628/27393 (64.4%) of cases, one variant matched with one match on one or both haplotypes, which was the basis for the "1 Line" categories in **Figure S7D,E**. In the remaining cases, a group of matched variants alleles/errors was spread across multiple lines in the hap.py vcf file. 7616/27393 (27.8%) of these were relatively simple variants that hap.py represented as two lines for some reason (for example, a "0|1" and "." for HG002v1.1 and HPRC respectively on one line followed by a "1|0" and "1|1" on the next line was likely a collapse that could have been written as "1|2" and "1|1"). These formed the basis for the "2 Line" categories in **Figure S7D,E**. All other cases (2149/27393) were deemed "complex" (example in **Figure S9**). Assigning each variant/variant group with its matches was performed according to the definitions in **Table S15** on the basis of their GT fields.

##### Equivalencies between variant benchmarking errors and genome benchmarking errors

In general, we expect that genome and variant benchmarking should be roughly equivalent for non-structural variants that lie within the confident regions of the GIAB benchmark, with the caveat that finding the correspondence between the two types of errors requires first projecting the genome errors onto GRCh38 coordinates which may be an imperfect operation. Within the variant benchmark regions, we found a few interesting relationships between genome and variant performance metrics (**Figure S7D,E** and **Figure S8C,D**): 1) what GQC calls "phasing" errors often appear as genotype errors ("collapse errors" in the assembly) where one such error corresponds to either one FN variant (collapse to reference) or one FN and one FP (collapse to non-reference), 2) a minority of phasing errors appear as phase switch errors, which are typically counted as true positive variants in hap.py but could be separately counted as errors by phasing benchmarking tools, 3) ~8% of genome errors fall in complex variants, primarily in tandem repeats and homopolymers (**Figure S7C**), so that a single genome benchmarking error can cause at least two FP or FN variant calls, (example in **Figure S10**). 4) Consensus errors most often corresponded to at least one FP variant.

##### Categories of variants and their relation to the genome errors provided by GQC

*Collapses*
One common type of variant-based error is known as a "collapse" where the true genotype and the test genotype are heterozygous and homozygous, respectively, and one allele is shared between the two. Usually this results when a given region has a difference between the two haplotypes but lacks enough reads to support this difference, resulting in the true sequences being "collapsed" into one sequence in the test genome. The collapsed sequence may match one of the true sequences or neither. In the latter case, we found these were almost always the result of the truth having two insertions/deletions (indels) of differing lengths and the collapsed

query sequencing having an intermediate length (**Figure S8B**); we thus called these “average collapses” below.

14538/30505 genome errors (47.7%) matched with collapses, and 70.8% of these were phasing genome errors (**Figure S8C**). 75.0% of these were “exact collapses” where one allele matched, and therefore each of these variants matched with exactly one genome error (**Figure S8D**). Within this subcategory, 95.2% were spread on one line with either one FN or one FN and FP, and the remainder were on two lines with one FN and FP. For those with one line in the VCF file, variants with only FN are cases where the collapse is to the reference allele (i.e., truth is 1|0, query is 0|0) and variants with FN/FP are those that collapse to alternate (1|0 to 1|1). Curation showed that collapses spread across two lines were likely cases involving a single-nucleotide polymorphism (SNP) and an indel, which hap.py will split into two lines to account for the two different variant types. 25.0% of genome errors which matched with a collapse variant matched in pairs (one for each haplotype), and these corresponded to “average collapses” as described above.

This analysis shows how variants and genome errors can be equated differently depending on the reference, which may lead to different benchmark performance metrics. For collapses, one genome error will correspond to one FN if the variant collapses to reference, but will also include an FP if the variant collapses to alternate. An average collapse will generally equate to one FP and one FN (89.5% in this case); when this isn’t true the variant is likely part of a larger, more complex cluster of variants that cause the labels to be counted differently.

*Sequencing Errors*

This variant category encompassed a variety of genotypes (**Table S15**) all of which were likely caused by an error in the underlying sequencing platforms. 100% of these (11962 variants) occurred in either a homopolymer or a tandem repeat, and 97.2% of these were INDELs (**Figure S7C**). These likely arose due to differences in length between the truth and query sequences. For instance, reverse collapses (so named because they have the opposite genotype manifestation relative to truth and query) were likely in a homopolymer that had a homozygous length in the truth but was either too long or too short on one haplotype in the query.

*Misphases*

A small number of variants (648) matched perfectly except their phasing was flipped which we called “Misphases”. hap.py marked these as true positives due to the fact that vcfeval (the comparison engine within hap.py) ignores phasing when comparing variants. Unsurprisingly, 87.2% of the corresponding genome errors were phasing errors in GQC, and all included two genome errors since both haplotypes mismatched the truth. This is the simplest instance where vcfeval will undercount true errors due to it not taking phasing into account. There are other instances where vcfeval scored a variant as TP even though the phasing did not match; in our data these were relatively complex cases and thus appeared under the “complex” heading (**Figure S7D,E**).

We also identified variants that were likely due to both a phasing flip and a sequencing error, which we called “Misphase/seq errors.” (ie 0|2 vs 2|1, so one allele mismatches, and the matching allele has the wrong phase). These were the only variant category for which the genome error type was “MIXED” (ie, there were two genome errors on each haplotype, one of which was a consensus error and one of which was a phasing error) (**Figure S7D,E**).

#### Supplementary Figures

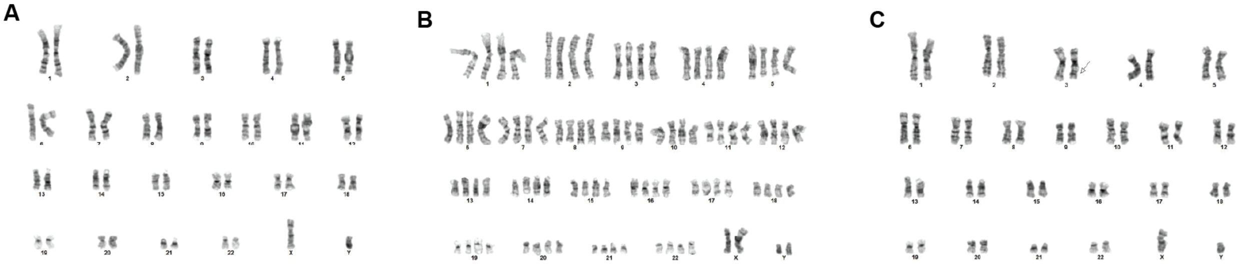

**Figure S1: Cytogenetic analysis of the GM24385/HG002 cell line. (A)** 53 of 73 metaphase cells (73%) showed a normal male karyotype (46,XY). **(B)** 13 cells (18%) showed a tetraploid karyotype (92,XXYY). **(C)** 3 karyotyped cells (4%) showed an inversion on chromosome 3 (46,XY,?inv(3)(q26.3q29), shown with an arrow).

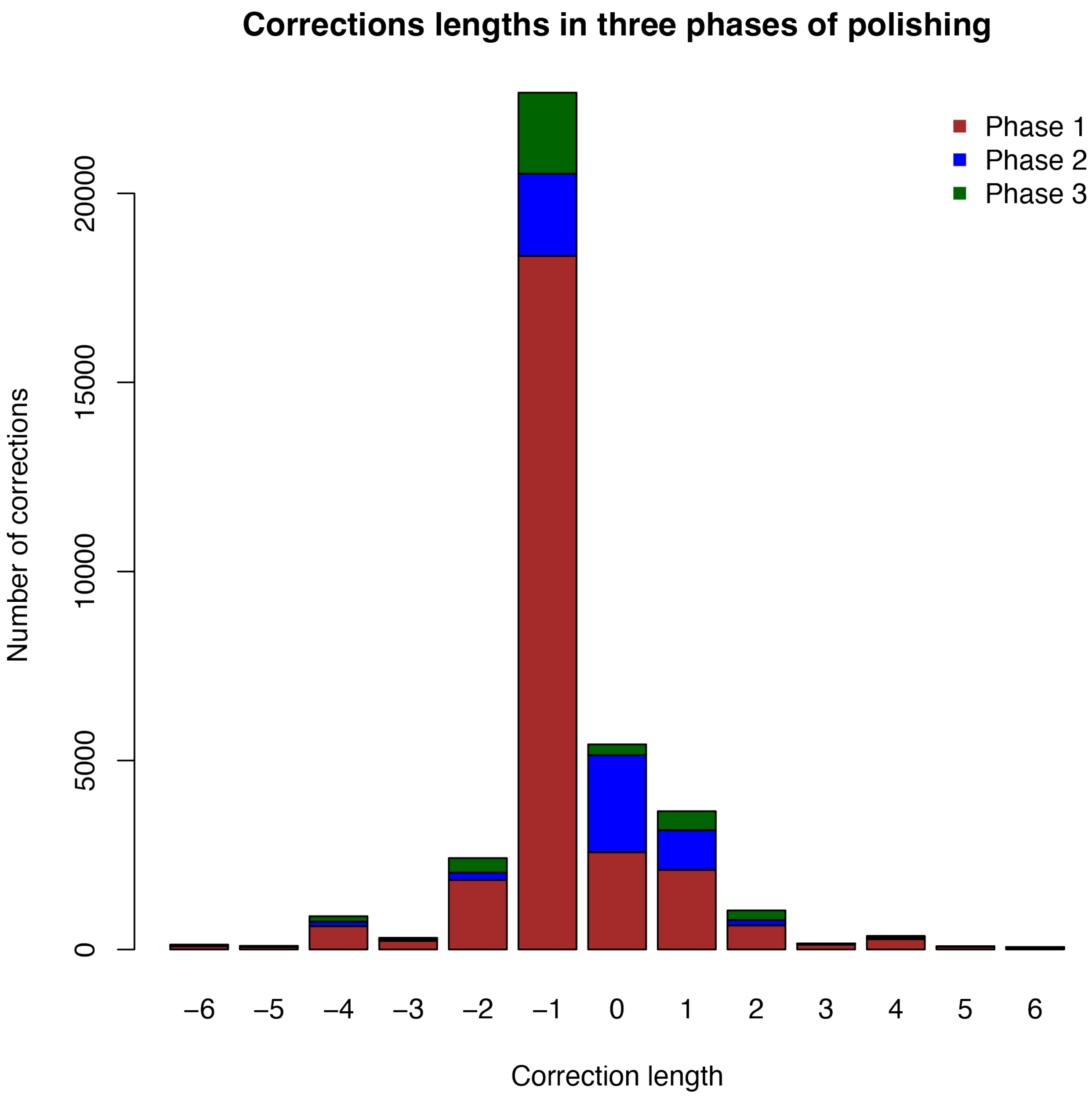


**Figure S2: Lengths of small corrections made in three rounds of polishing.** Histogram of correction lengths in polishing HG002v0.7 to create HG002v0.9 (Phase 1), in polishing HG002v0.9 to create HG002v1.0.1 (Phase 2), and in polishing HG002v1.0.1 to create HG002v1.1 (Phase 3). The length of a correction is defined as the difference between the new (corrected) allele length and the original (uncorrected) allele length. Roughly the same number of single nucleotide corrections (length 0) were made in Phases 1 and 2 (2574 and 2555, respectively), with a smaller number (291) in Phase 3. The plot shows that the large majority of corrections made overall were deletions of single bases, usually shortening homopolymer sequences that were found to be one base too long.

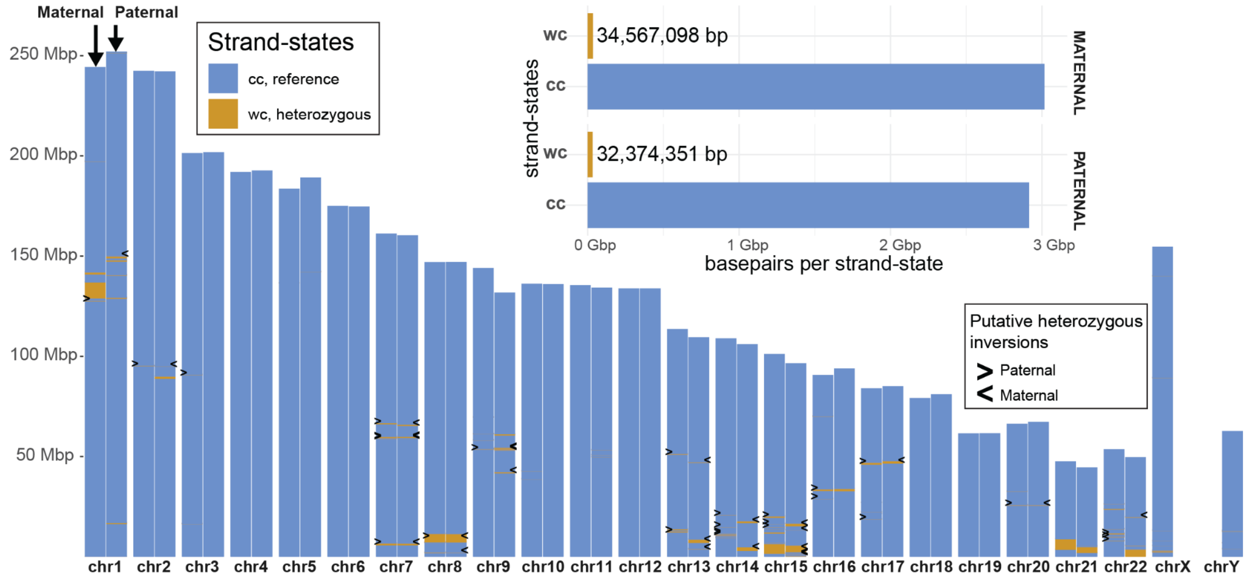


**Figure S3: Strand-seq assembly evaluation of HG002v1.1.** An ideogram shows Strand-seq-based genotyping marking regions genotyped as in complete agreement with the assembly (reference, 'cc') and regions genotyped as heterozygous ('wc'). Heterozygous regions are generally caused by low mappability regions such as short acrocentric arms, centromeres, and repetitive regions. A subset of these mark positions of heterozygous inversions, highlighted by arrowheads. We did not observe any region genotyped as homozygous inverted, suggesting that no misorientation is present in these haplotype assemblies.

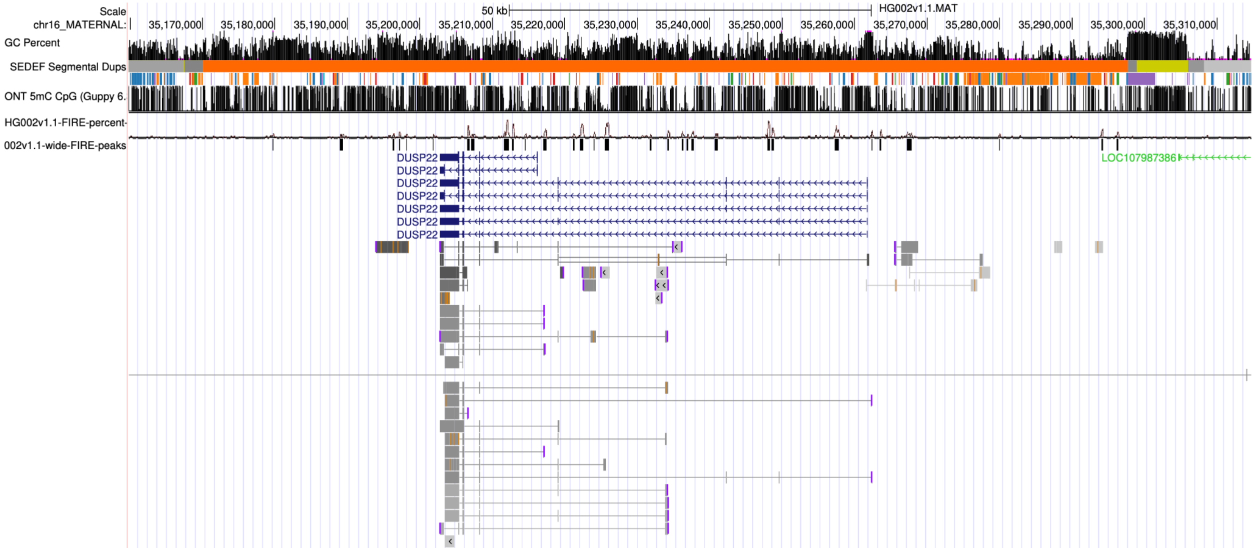


**Figure S4: UCSC Browser hub for T2T-HG002v1.1.** The browser includes diploid gene annotation and functional genomics tracks, as well as sequence-based annotations (GC percent, CpG islands, repetitive elements, segmental duplications, subtelomere repeats, and centromeric satellites) and validation tracks (known issues, depth of coverage, NucFreq output, etc.). Chain file tracks are also included, showing the position of all heterozygous sites and enabling quick locus switching between the two HG002 haplotypes, T2T-CHM13, and GRCh38. Shown here is the maternal haplotype of HG002 Chr16 focused on the *DUSP22B* gene from **Figure 2C**. Browser tracks shown from top to bottom: GC percent, segmentally duplicated regions (orange representing a highly similar duplication), RepeatMasker annotations, ONT-based methylation percentage, Fiber-seq accessibility, Fiber-seq peak calls, gene models, and PacBio Kinnex transcripts. The methylation, Fiber-seq, and transcriptome data jointly support the expression of this additional *DUSP22* copy in the HG002 LCL.

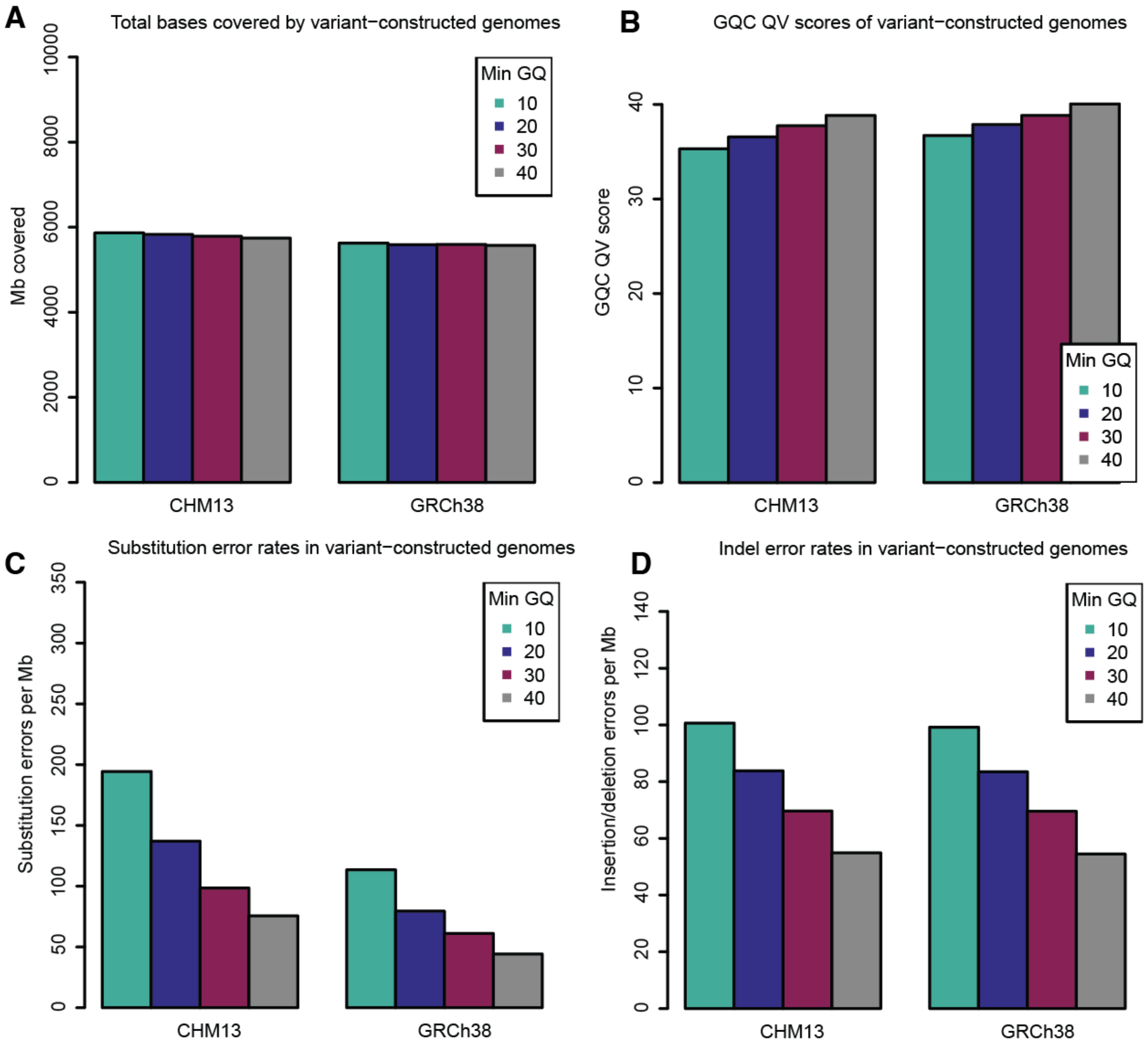


**Figure S5: GQC quality statistics for variant-constructed genomes using different minimum genotype quality cutoffs. (A)** Aligned coverage of v1.1, or total genome benchmark bases covered by aligned sequence in the variant-constructed genome. **(B)** GQC quality scores (phred-scaled) for variant-constructed genomes calculated as -10 times the log, base 10 of the total number of discrepancies divided by the total alignment length. **(C)** Substitution error rate in variant-constructed genomes per megabase of aligned sequence. **(D)** Indel error rate in variant-constructed genomes per megabase of aligned sequence.

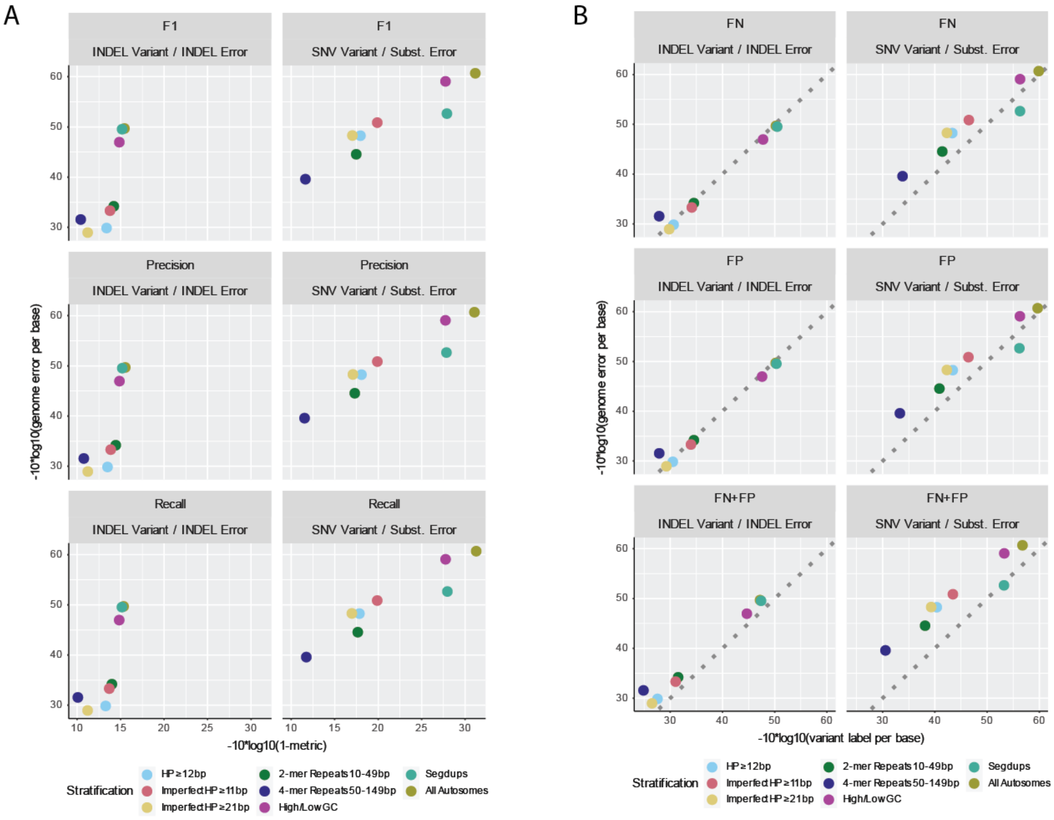


**Figure S6: Global performance metric comparison between variant and genome benchmarking across various stratifications**. **(A)** Genome errors per base (as reported by GQC) vs F1, precision, or recall reported as a variant benchmarking performance metric (phred-scaled). Genome errors were restricted to those which can be successfully projected onto GRCh38 GIAB small variant benchmark regions. Furthermore, SNV and INDEL designations for genome errors were based on comparison of an HG002 assembly to the benchmark while variant benchmarking SNV and INDEL designations were based on comparison of HG002 to GRCh38. HP = homopolymer. **(B)** Phred-scaled genome benchmarking errors per base vs. phred-scaled false negative and false positive variants per base for SNVs/Substitution Errors or INDELs/INDEL Errors. Points above the dotted line indicate variant benchmark had more errors relative to genome benchmark.

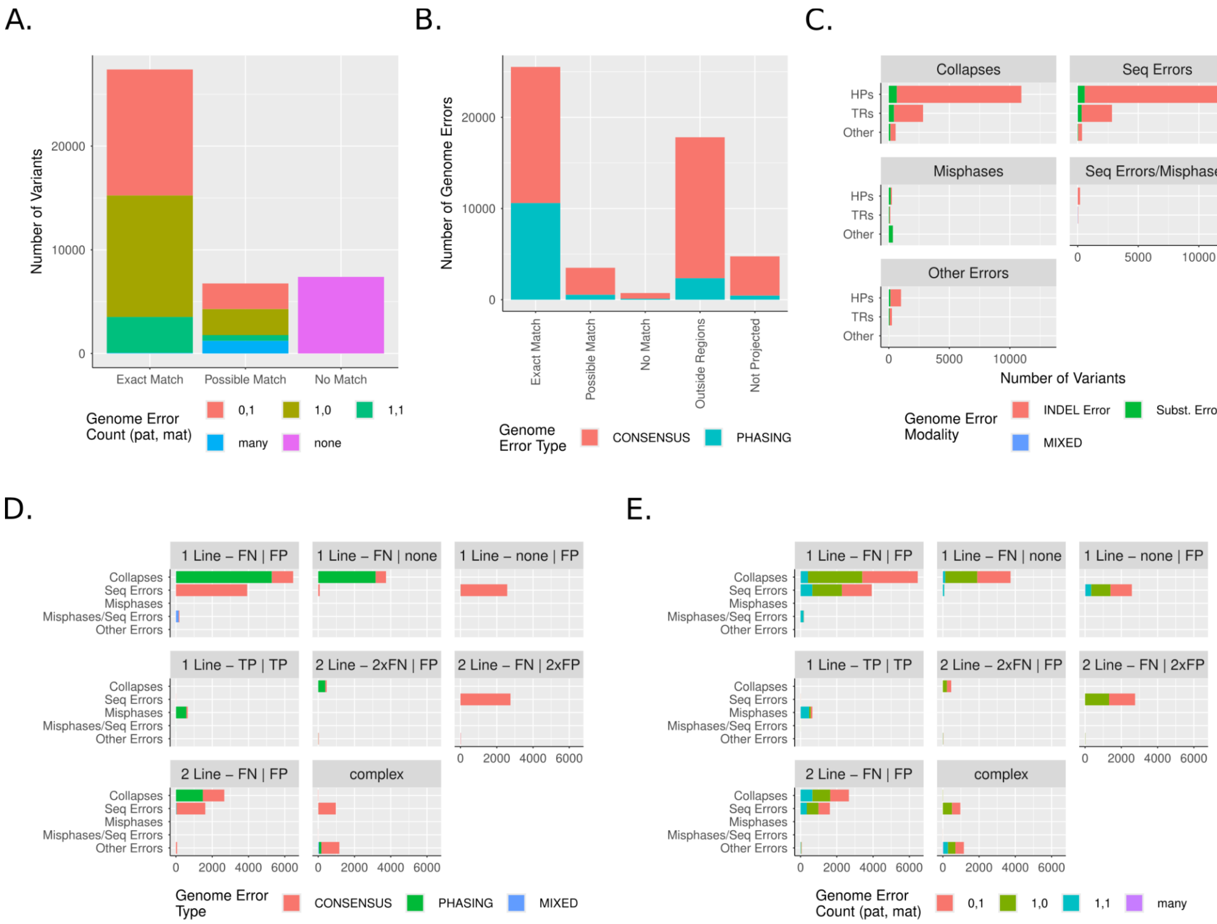


**Figure S7: Variants and their corresponding genome benchmarking errors. (A)** Variant-based errors grouped by those that matched with a genome error ("Exact Match"), intersected but failed to match with a genome error ("Possible Match") or intersected with no genome errors ("No Match") colored by how many genome errors were part of the intersection. **(B)** Genome errors grouped by those that matched, intersected, or did not intersect with a variant analogous to **(A)**; in addition, genome errors that projected from T2T-HG002v1.1 to GRCh38 coordinates but landed outside the benchmarking regions ("Outside Regions") or failed to lift over entirely ("Not Projected"). **(C)** Variants grouped by presence in homopolymers (HP), tandem repeats (TRs), or other stratifications. "Substitution Errors" are differences between the T2T-HG002v1.1 and HPRC assemblies for which both alleles are only a single base, and "INDEL Errors" are everything else. (**D-E)** Variants with at least one matching genome error grouped by variant type (y axis) and label/number of lines (header) and either colored by the type of matching genome error(s) **(D)** or number of matching genome errors **(E)**.

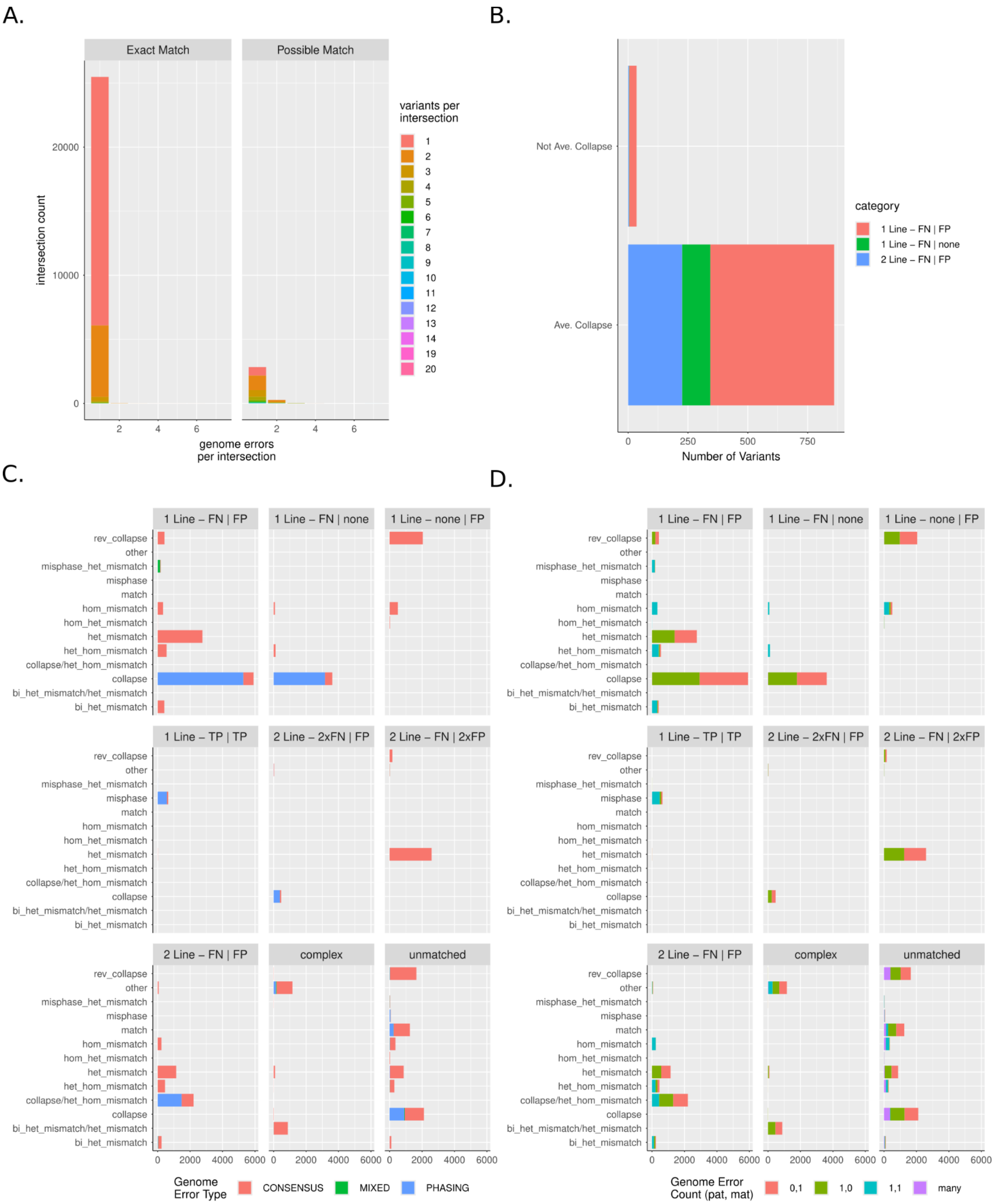


**Figure S8: Variants match results and corresponding genome errors with granular categorization** **(A)** counts of variants and genome errors per intersection that either match (left) or do not match (right); in this case variants are split across both haplotypes so homozygous variants are included twice. **(B)** Counts of variants that are het_hom_mismatches that are “average collapses” or not **(C-D)**

Corresponding categories analogous to **Figure S7C,D** broken down into finer subcategories that correspond directly to variant genotype. See **Table S15** for how these categories are defined.

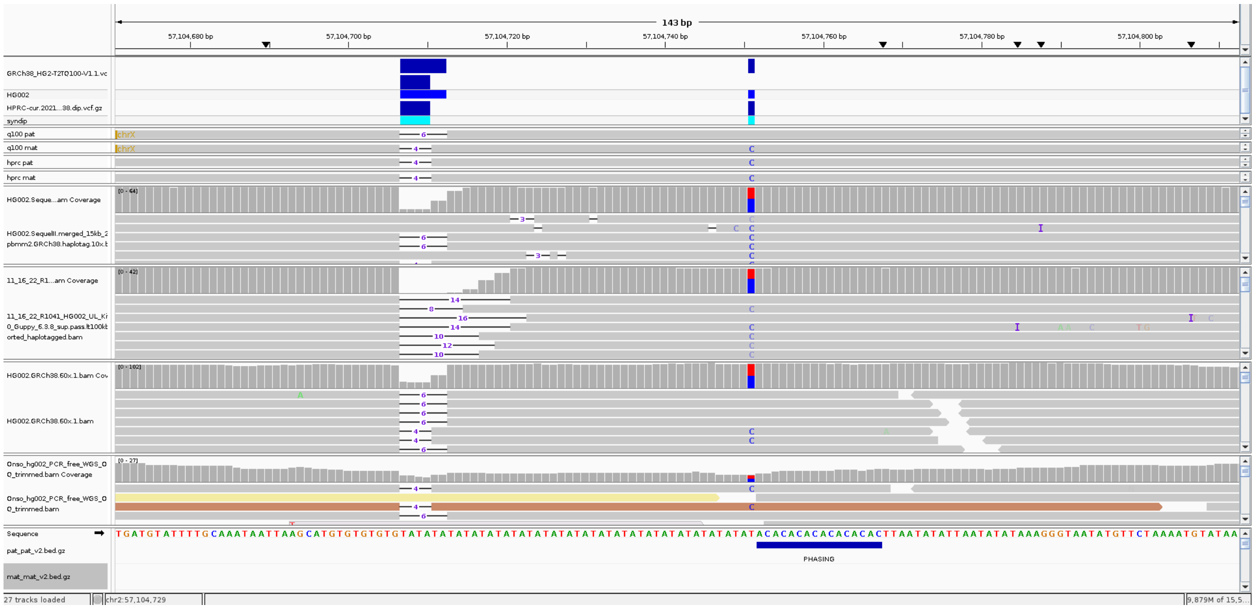


**Figure S9: Example on chr2:57,104,564-57,104,848 (GRCh38)** showing a collapse genome error which is represented as 3 variants. The net effect is that AC is inserted on the HPRC paternal assembly relative to the paternal T2T-HG002v1.1 assembly. The three variants are described in **Table S18: Example of a complex variant.** The query deletes TATA relative to reference. The truth deletes TATA and converts the last T to a C on the maternal haplotype and deletes TATATA in the paternal haplotype. This complexity arises because the maternal T2T-HG002v1.1 haplotype and both HPRC haplotypes insert AC but remove TATA relative to GRCh38 reference.

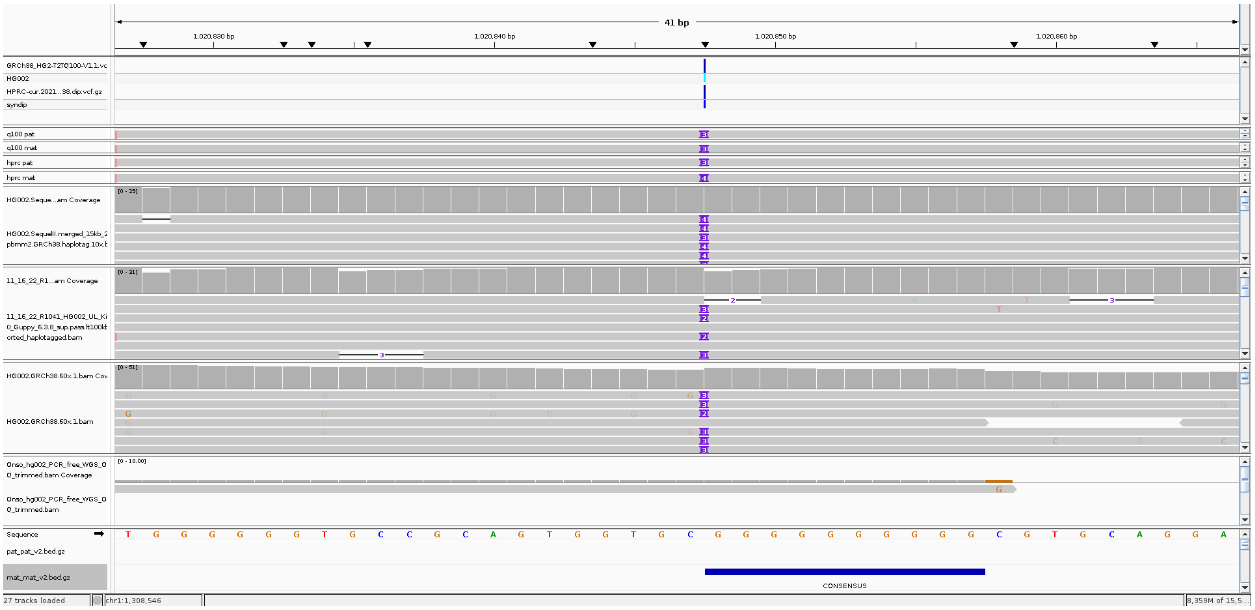

**Figure S10: Example on chr1:1,020,827-1,020,866 (GRCh38)** showing collapse genome error which is equivalent to a FP and FN variant on the maternal haplotype. The net effect is that one G is inserted on the HPRC maternal assembly relative to the maternal haplotype of HG002v1.1.

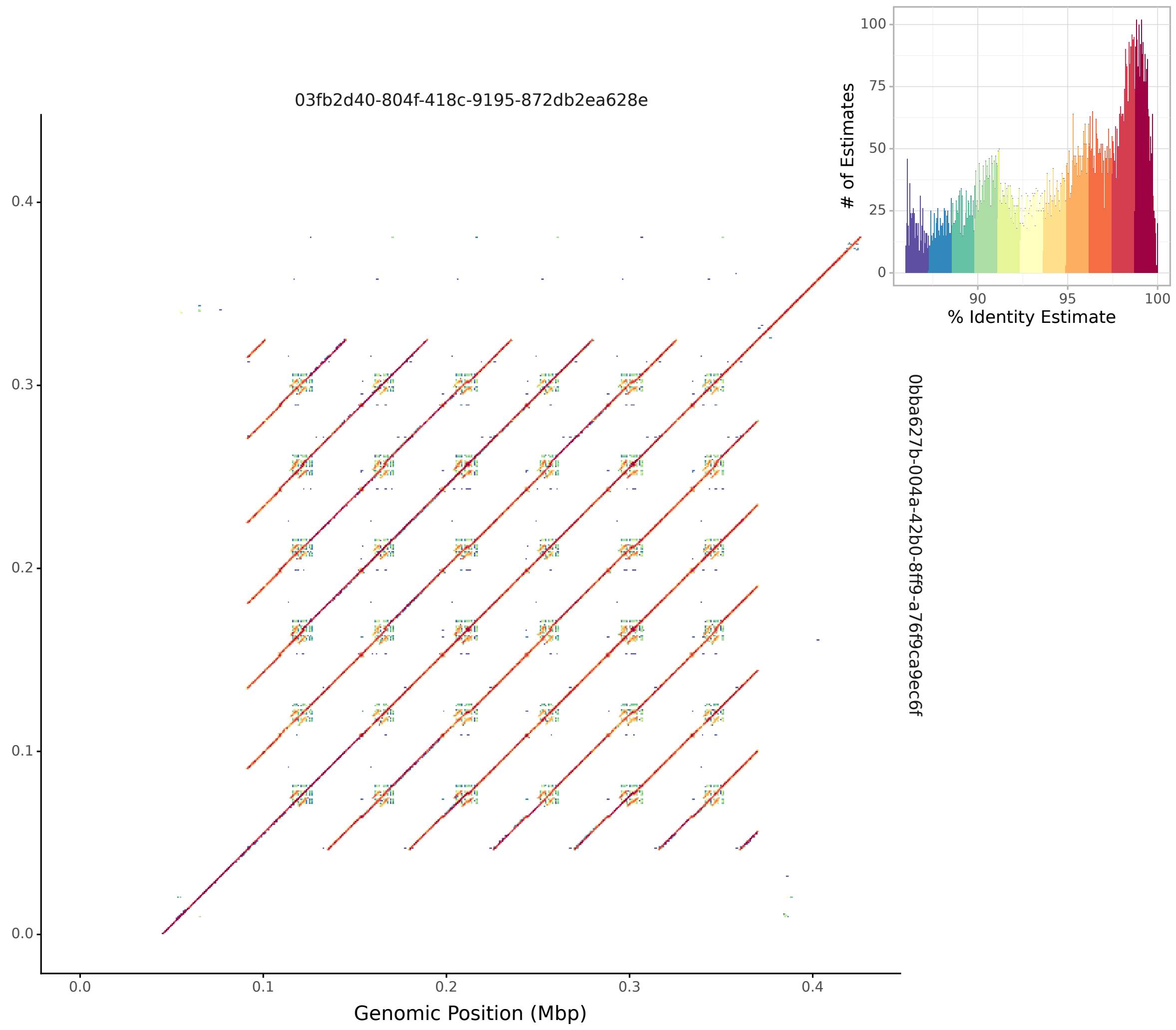


**Figure S11: ModDotPlot plot of two ONT ultralong reads which span the paternal chr13 rDNA array.** Ultralong reads indicate six copies of the rDNA repeat unit, longer than the v0.9 consensus of chr13_PATERNAL's acrocentric p-arm.
